# Design of Transmembrane Mimetic Structural Probes to Trap Different Stages of γ-Secretase-Substrate Interaction

**DOI:** 10.1101/2021.08.05.455313

**Authors:** Sanjay Bhattarai, Sujan Devkota, Michael S. Wolfe

**Affiliations:** Department of Medicinal Chemistry, University of Kansas, Lawrence, 66045, KS, USA

**Keywords:** aspartyl protease, membrane proteins, peptidomimetics, helical peptides, transition-state analogs

## Abstract

The transmembrane domain (TMD) of the amyloid precursor protein of Alzheimer’s disease is processively cut by γ-secretase through endoproteolysis and tricarboxypeptidase “trimming”. We recently developed a prototype substrate TMD mimetic for structural analysis—composed of a helical peptide inhibitor linked to a transition-state analog—that simultaneously engages a substrate exosite and the active site and is pre-organized to trap the carboxypeptidase transition state. Here we developed variants of this prototype designed to allow visualization of transition states for endoproteolysis, TMD helix unwinding, and lateral gating of substrate, identifying potent inhibitors for each class. These TMD mimetics exhibited non-competitive inhibition and occupy both exosite and active site as demonstrated by inhibitor cross competition experiments and photoaffinity probe binding assays. The new probes should be important structural tools for trapping different stages of substrate recognition and processing via ongoing cryo-electron microscopy with γ-secretase, ultimately aiding rational drug design.

## Introduction

γ-Secretase is a founding member of a class of intramembrane-cleaving proteases (I-CLiPs), enzymes with membrane-embedded active sites that hydrolyze substrate transmembrane domains (TMDs) within the hydrophobic interior of the lipid bilayer.^1^ To accomplish this seemingly paradoxical process, γ-secretase and other I-CLiPs must recognize substrate TMD, allow lateral entry into an internal water-containing active site, and bend or unwind the α-helical TMD to make the scissile amide bond available for cleavage. How these enzymes carry out such intricate substrate recognition and processing is unclear. Among I-CLiPs, γ-secretase is unique in being composed of four different membrane proteins, with presenilin as the catalytic component of an aspartyl protease.^2, 3^ Upon assembly with the other components (nicastrin, Aph-1 and Pen-2),^4–6^ presenilin undergoes autoproteolysis^3, 7^ to an N-terminal fragment (NTF) and C-terminal fragment (CTF), with each presenilin subunit contributing one of the two catalytic aspartates to what is now the active γ-secretase complex.

The γ-secretase complex hydrolyzes >100 known substrates, including the amyloid precursor protein (APP) of Alzheimer’s disease and the Notch family of developmental signaling receptors.^8^ TMD cleavage of APP toward the production of 42-residue amyloid β-protein (Aβ) is involved in the pathogenesis of Alzheimer’s disease (AD),^9^ while TMD cleavage of Notch receptors is an essential step in their proto-oncogenic signaling pathways.^10^ Thus, γ-secretase inhibitors and modulators have potential as treatments for AD and cancer.^11–14^ An additional level of complexity of γ-secretase involves processive proteolysis. The enzyme first carries out endoproteolysis near the cytosolic end of the APP TMD, at ε cleavage sites, to release the APP intracellular domain, with 48- or 49-residue Aβ peptide intermediates further processed by γ-secretase through a tricarboxypeptidase activity.^15^ Previously we and others developed transition-state analog inhibitors^16–20^ (TSAs, e.g. **1**, Fig. 1) that target the active site^21, 22^ as well as potent helical peptide inhibitors (HPIs, e.g. **2**)^23, 24^ that bind a substrate-docking exosite on γ-secretase.^25^ Use of these chemical probes revealed that the two sites are distinct but in close proximity.^25^ TSA probes also suggested active-site pockets that accommodate the side chains of three substrate residues C-terminal to the cleavage site (residues P1’, P2’ and P3’),^20^ and more recent studies support this three-pocket model for dictating tripeptide trimming.^26^

**Figure 1.**
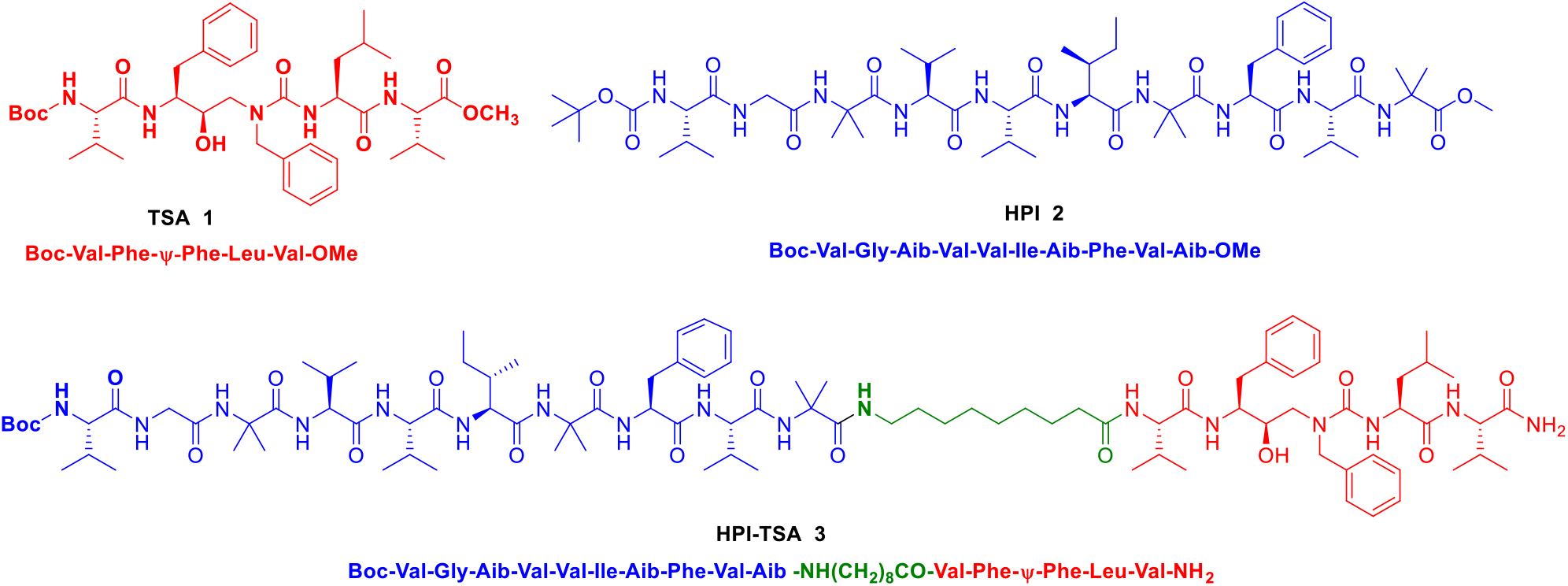
Structures of selected peptidomimetic γ-secretase inhibitors. Compound **1** is a hydroxyl ethyl urea (Ψ)-containing transition state-analogue inhibitor (TSA), which binds to the active site of the enzyme. Compound **2** is an α-aminoisobutyric acid (Aib)-containing helical peptide inhibitor (HPI), directed to a substrate-binding exosite. Compound **3** is a HPI-TSA conjugate, with HPI and TSA moieties connected through a ω-aminononoyl linker.

Advances in cryo-electron microscopy (cryo-EM) paved the way to the first detailed structures of the γ-secretase complex.^27^ Subsequently, structures of γ-secretase bound to APP- and Notch-based substrates were elucidated;^28, 29^ however, in both cases the enzyme was catalytically inactive, and substrate was covalently linked through artificial disulfide bonds. Most recently, structures of active enzyme bound to several inhibitors and to a modulator were also solved.^30^ Despite these advances, structural understanding of the multiple steps in substrate recognition and processive proteolysis in the lipid bilayer remains largely unclear. Our goal was to engineer a new class of inhibitors based on an entire substrate TMD, to trap the active enzyme in a conformation poised for TMD hydrolysis for study by cryo-EM. We recently reported the discovery of the first such TMD mimetic structural probe,^31^ HPI-TSA conjugate **3** (Figs. 1C and 2A), designed to trap the tricarboxypeptidase transition state of the enzyme (Fig. 2B vs. 2C). We demonstrated that this new probe is a stoichiometric inhibitor of γ-secretase (*K*_i_ of 0.42 ± 0.12 nM), assumes a solution conformation similar to enzyme-bound substrate (i.e., is preorganized for binding to γ-secretase), and interacts simultaneously with the active site and with a substrate-docking exosite.^31^ Encouraged by preliminary cryo-EM images showing stoichiometric inhibitor **3** bound to γ-secretase in a manner similar to substrate (Y. Shi, R. Zhou, and G. Yang, unpublished results), we expanded our work to design and develop variations on structural probe **3** that should allow visualization of the transition states not only for carboxypeptidase cleavage, but also for ε proteolysis, TMD helix unwinding, and lateral gating of substrate (Figs. 2D-F).

**Figure 2.**
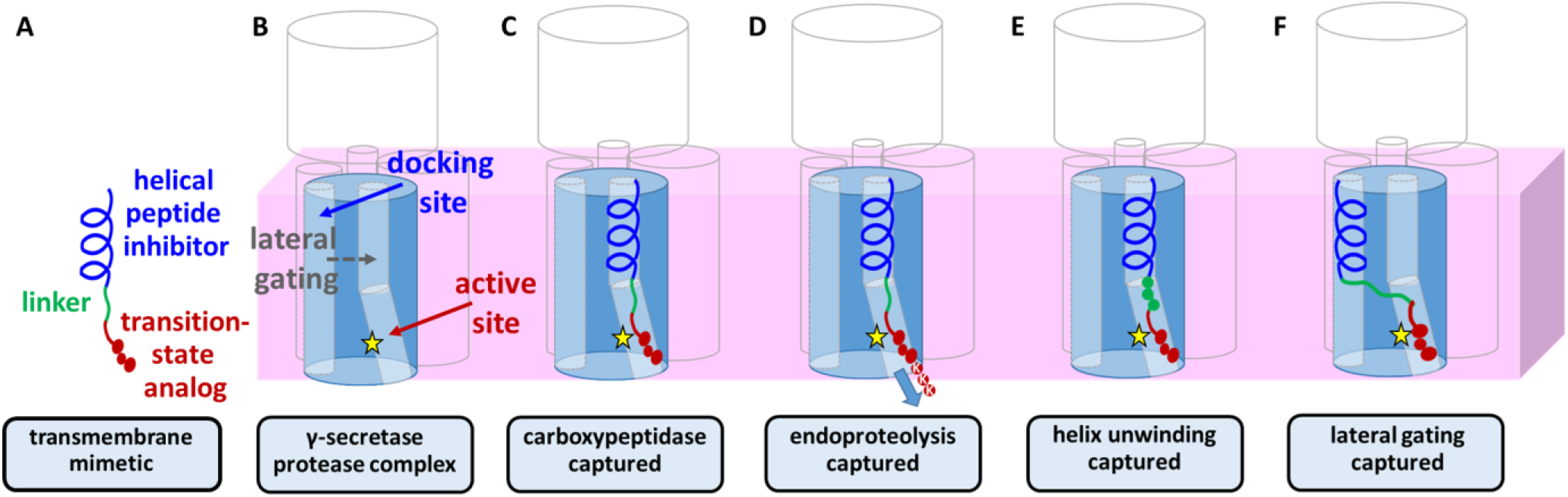
Design of substrate transmembrane mimetic inhibitors of γ-secretase substrate. **A**). Recently reported stoichiometric γ-secretase inhibitor containing an L-helical peptide inhibitor (HPI, blue) that engages the substrate docking exosite connected through a linker (green) to a transition-state analogue inhibitor (TSA, red) targeted to the active site. **B**) The active γ-secretase complex is composed of presenilin (blue cylinder) and three other membrane proteins (outlined). Transmembrane substrate docking is followed by lateral gating, with helix unwinding, into the internal active site with catalytic aspartates (yellow star). **C**) In the design of prototype HPI-TSA conjugate **3**, the TSA moiety has only three P’ residues (P1’, P2’, P3’) and should thereby trap the protease at the transition state of carboxypeptidase trimming. **D-F**). In this study, we developed variants of HPI-TSA **3** that should allow visualization of transition states for endoproteolysis, TMD helix unwinding, and lateral gating of substrate. **D**) Extending the C-terminus into P4’, P5’, P6’ with lysines (K) should trap the transition state of endoproteolysis. **E**) Replacement of the ten-atom linker with three amino acids may provide insight into the nature of helix unwinding, as this region connects the helical N-terminus to the extended C-terminus. **F**) To trap the gating process, the L-HPI moiety was replaced with a potent D-HPI. The opposite helical handedness of the D-HPI may prevent conformational changes required for lateral entry into presenilin.

For the prototype structural probe HPI-TSA conjugate **3**, the TSA component has only three C-terminal amino acid residues, i.e. P1’, P2’, P3’, capable of binding to three substrate pockets in the active site (Fig. 2C) and should thereby trap the carboxypeptidase trimming process.^26^ We therefore sought to extend the C-terminus of **3** with lysine residues P4’, P5’, P6’ in APP, which should trap γ-secretase at the transition state of ε proteolysis (Fig. 2D). In the γ-secretase structure bound to APP substrate,^28^ the P4’, P5’, and P6’ lysine residues are seen in an extended conformation in interacting with presenilin. In addition, we replaced the alkyl linker of the HPI-TSA conjugate **3** with natural amino acids (Fig. 2E). This region in substrate TMD is partially unwound in the bound substrate structures^28, 29^ and located at the juncture of the helical and extended regions. Therefore, this region is apparently critical to unwinding the C-terminal region of the substrate TMD and setting up the transition state for proteolysis of the TMD. The optimal 10-atom alkyl spacer in conjugate **3** is close in length to 3 amino acids. Thus, we replaced this linker with the corresponding residues in the APP TMD. We also planned to replace the linker with glycines, which are helix-destabilizing and lead to dramatic increases in endoproteolytic processing by γ-secretase.^32^ Finally, the pathway for lateral gating of substrate TMD into the active site interior of presenilin is unclear. To develop a probe to trap the lateral gating process, we swapped the L-HPI **2** of HPI-TSA conjugate **3** with a potent D-HPI (Fig. 2F).^23^ While the left-handed helicity of the D-HPI apparently presents an array of hydrophobic side chains for initial interaction with γ-secretase that is similar to that presented by L-HPIs, helical handedness may be critical for conformational changes during lateral gating. We therefore considered that D-HPIs may dock with γ-secretase but not be drawn inside presenilin,^25^ allowing identification of the site of lateral gating.

Generating cryo-EM structures of active γ-secretase in different states of substrate recognition and processing would greatly inform drug discovery for Alzheimer’s disease. In particular, the new structures should reveal mechanisms of processive proteolysis of long Aß peptides to secreted shorter forms by γ-secretase; this trimming process is deficient with familial Alzheimer’s disease (FAD) mutations found in APP and presenilins.^33–35^ Understanding of specific structural and functional changes caused by FAD mutations may suggest strategies to identify small drug-like molecules that bind allosterically and restore normal function, as both chemical tools and therapeutic prototypes.

## Results and Discussion

### Chemistry

For the preparation of target HPI-TSA conjugates, a divergent synthetic strategy was applied, which involved the construction of tripeptidomimetic building block **11** in a multi-step procedure (Scheme 1), followed by solid-phase peptide synthesis (SPPS) using this building block and Fmoc-protected amino acids (Scheme 2). In most cases, N-terminal *t*-butoxycarbonyl (Boc) was desired in the final product, as with prototype **3**, requiring solution-phase attachment of N-Boc-valine as the final step according to our previously described procedure^31^ with some modifications. Building block **11** was synthesized from the commercially available oxirane **4** as previously reported (Scheme 1). Oxirane **4** was opened under basic condition to hydroxyethylamine derivative **5** followed by reaction with isocyanate of L-leucine methyl ester to obtain hydroxylethylurea **6,** à la Nowick *et al*.^36^ The Boc protecting group of **6** was then swapped with Fmoc group resulting in intermediate **8**, followed by orthogonal TBDMS protection of the sterically hindered hydroxyl group (**9**). Hydrolysis of the methyl ester functionality of **9** resulted in carboxylic acid **10**, and final reinstallment of Fmoc provided **11** in an overall yield of 60%. HPI-TSAs **20-39** were synthesized using building block **11** by solid-phase coupling on Rink amide resin according to Scheme 2. All natural amino acids as well as building block **11** were sequentially added, and the peptidomimetics were released from the resin using a cleavage cocktail. For final peptides containing an N-terminal acetyl group, the peptides were capped after the last coupling step and then cleaved from the resin (Note: peptides **20, 21, 30** and **31** were synthesized as peptide **16** of Scheme 2). The O-TBDMS protecting group in building block **11** was removed during cleavage of peptides from the Rink amide resin, as the cleavage cocktail contains TFA. For the synthesis of N-Boc products, terminal Boc-Val-OH was finally added to the penultimate peptidomimetics by coupling with HATU in DIPEA in solution phase (Note: peptides **22-29** and **33-39** were synthesized as peptide **18** of Scheme 2). All final peptides were purified by preparative HPLC using a C8 column. The purity of all tested peptides were >95%.

**Scheme 1.**
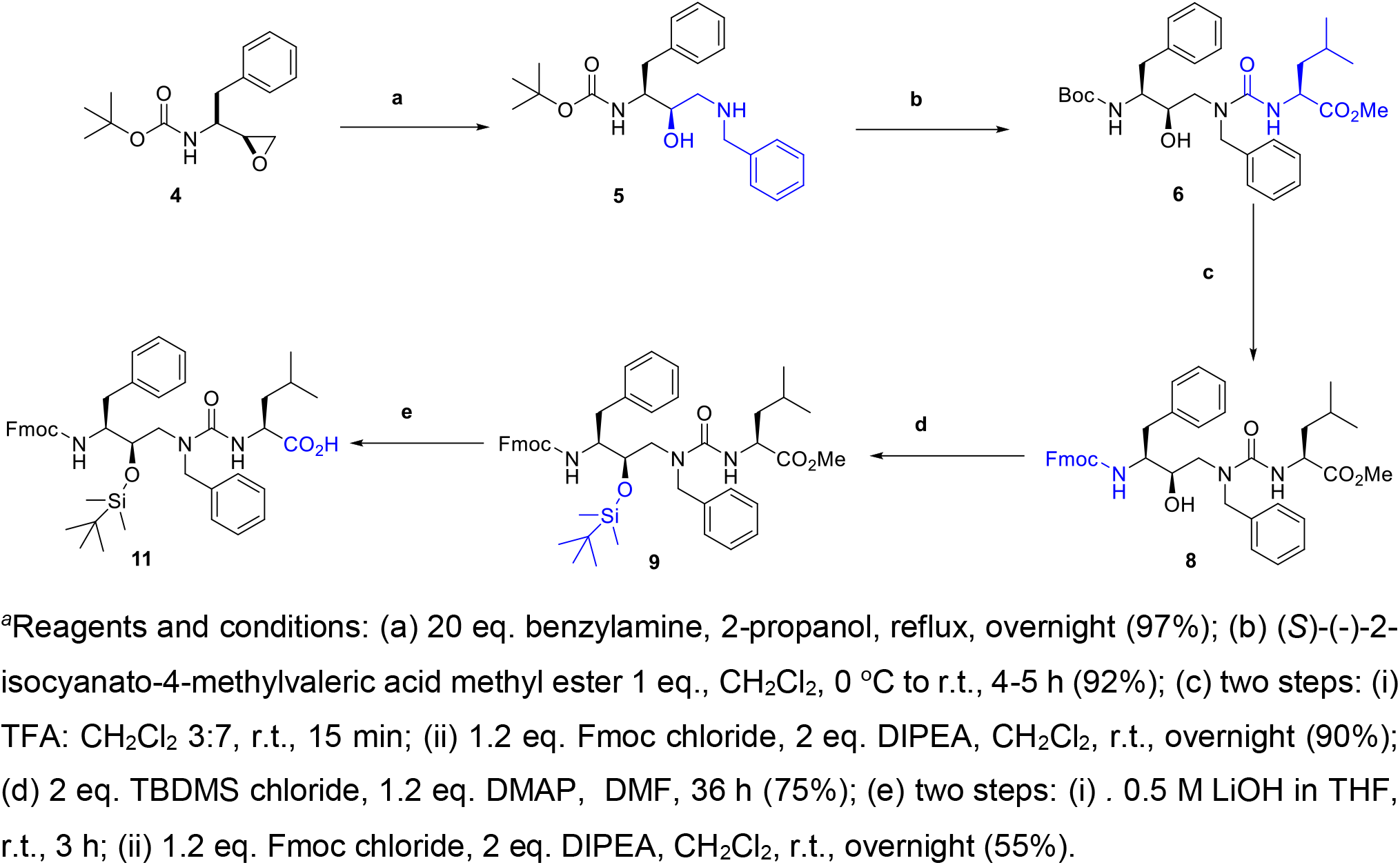
Synthesis of tripeptidomimetic building block **11**^*a*^

**Scheme 2.**
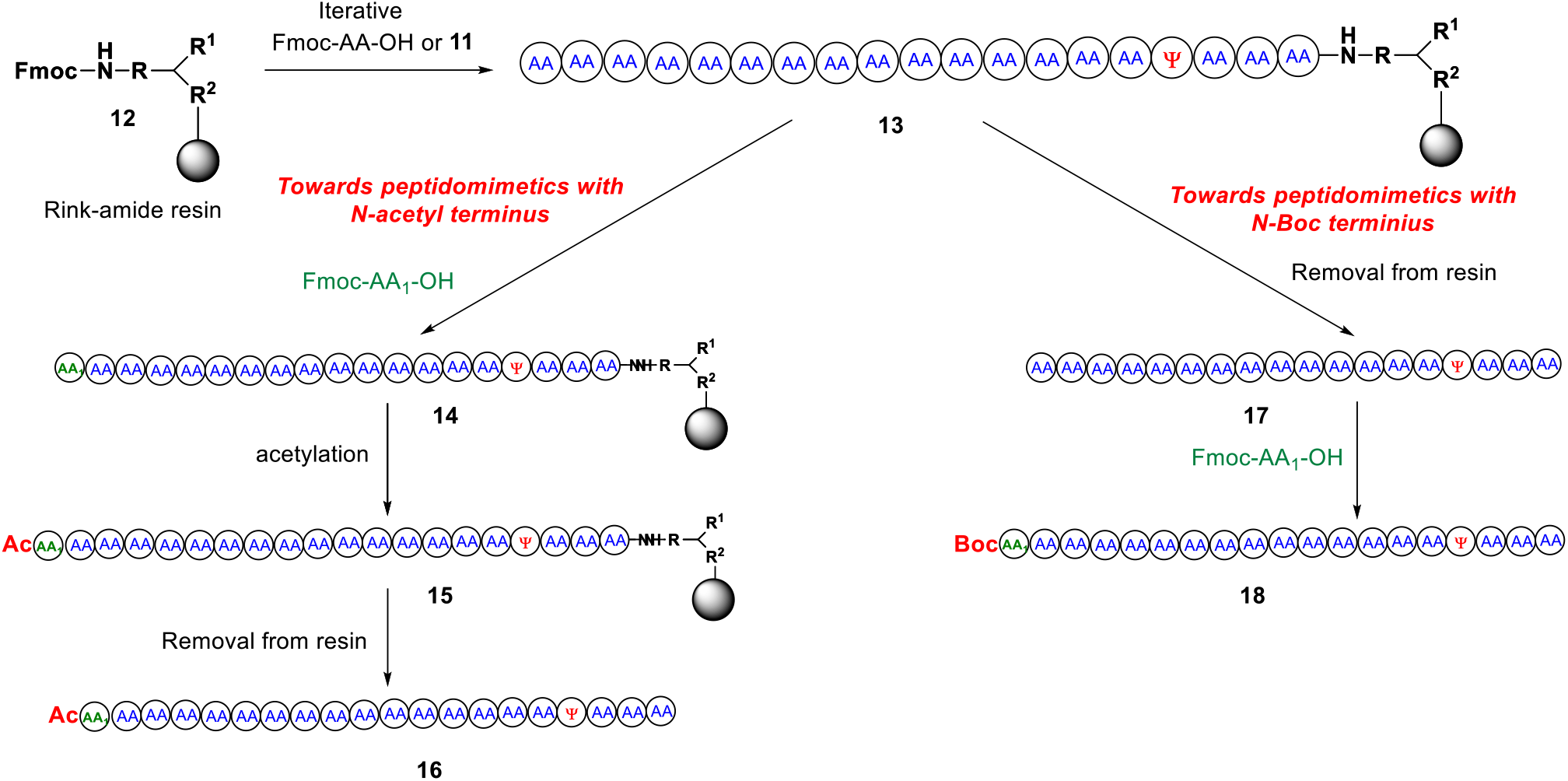
Strategy for solid-phase synthesis of TMD mimetic HPI-TSI conjugates.

### Pharmacological Evaluation

The inhibitory potencies of these peptidomimetics toward γ-secretase were determined using a highly sensitive sandwich ELISA assay for major proteolytic product Aβ40 formed from purified protease complex^37, 38^ and APP-based recombinant substrate C100Flag^39^ as previously reported.^*31*^ (Note: As expected from their mechanisms, TSAs and HPIs inhibit Aβ40 and Aβ42 production with similar potencies.^23, 32, 40^) The four-component γ-secretase complex was expressed from tetracistronic vector pMLINK in suspended HEK (human embryonic kidney) 293 cells.^37, 38^ 1 nM of purified γ-secretase in a standard detergent-lipid assay buffer was incubated with 500 nM C100Flag substrate and varying concentrations of inhibitor for 2 h at 37 °C. Results are summarized in Table 1 and 2, and selected concentration-inhibition curves are shown in Figure 3. In addition, we determined the inhibition constants (*K*_i_) of selected potent peptidomimetic inhibitors as above, but with varying concentrations of C100Flag at set inhibitor concentrations. The inhibition data, shown as double-reciprocal plots in Figure 4, were fit to a noncompetitive inhibition equation for *K*_i_ calculation.

**Table 1.**
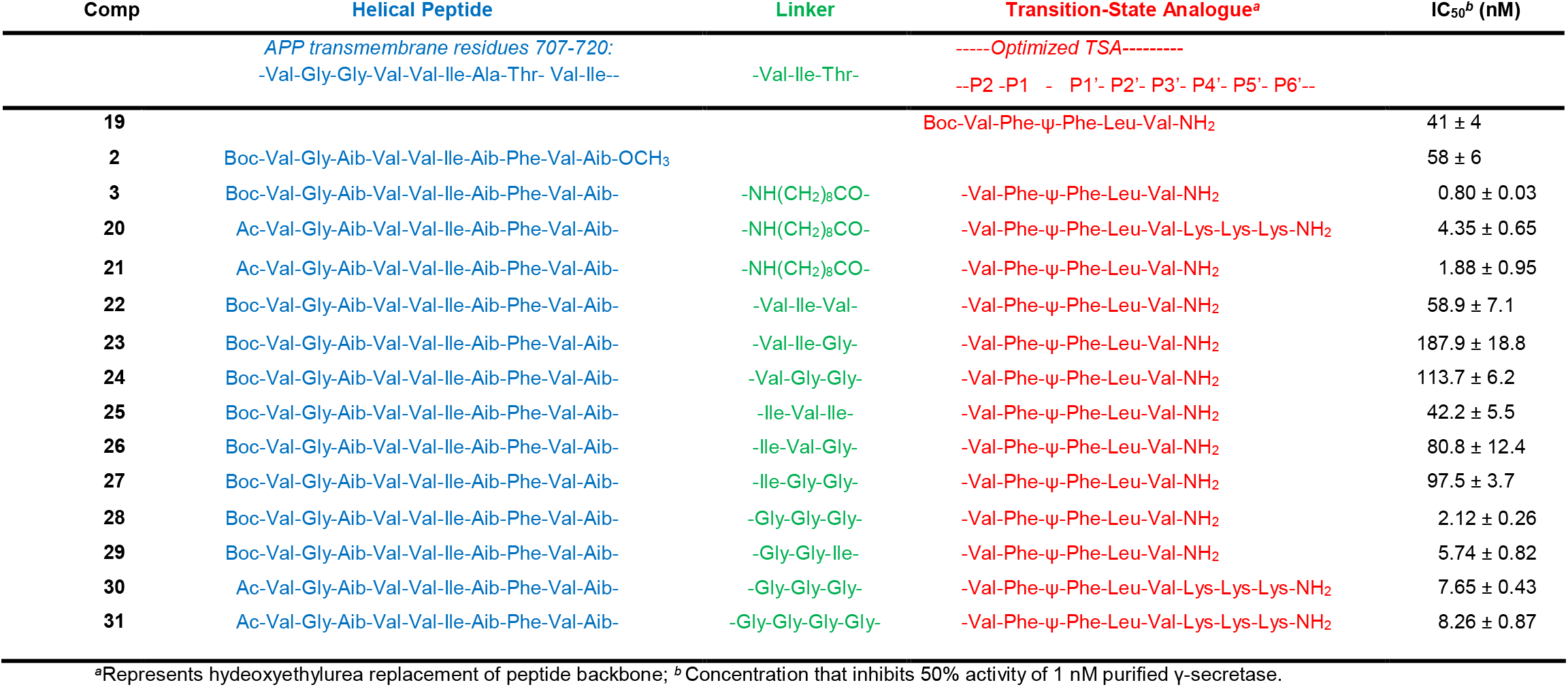
γ-Secretase inhibition by HPI-TSA conjugates designed to trap endopeptidase and helix unwinding stages of substrate recognition.

**Table 2.**
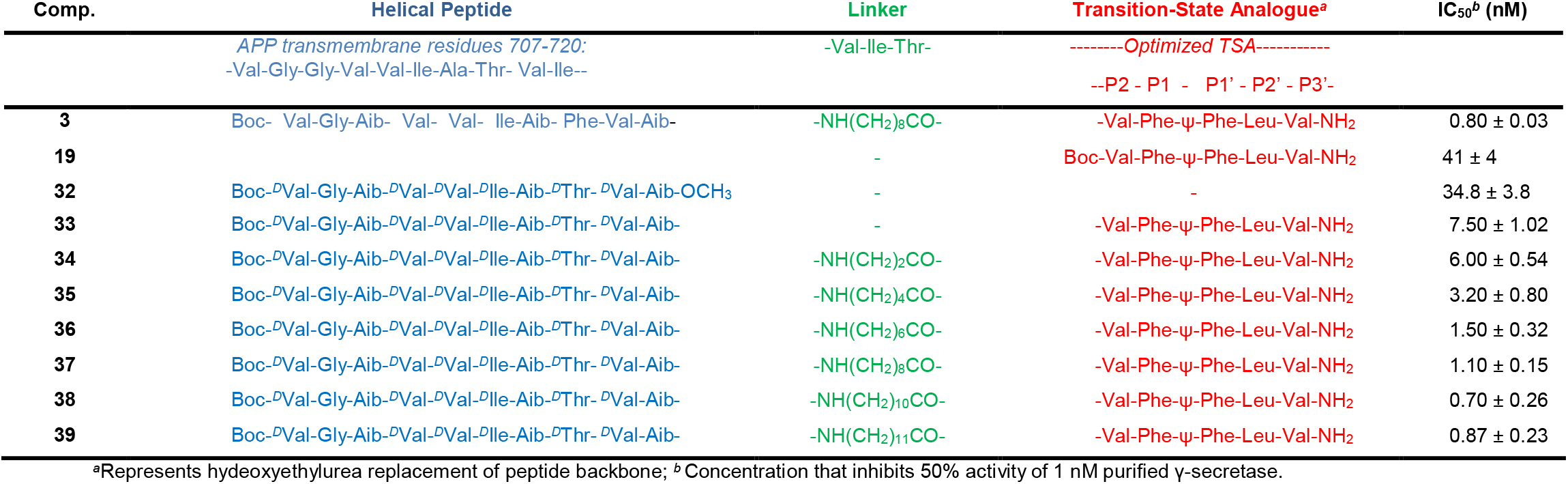
γ-Secretase inhibition by HPI-TSA conjugates designed to trap the lateral gating stage of substrate recognition.

**Figure 3.**
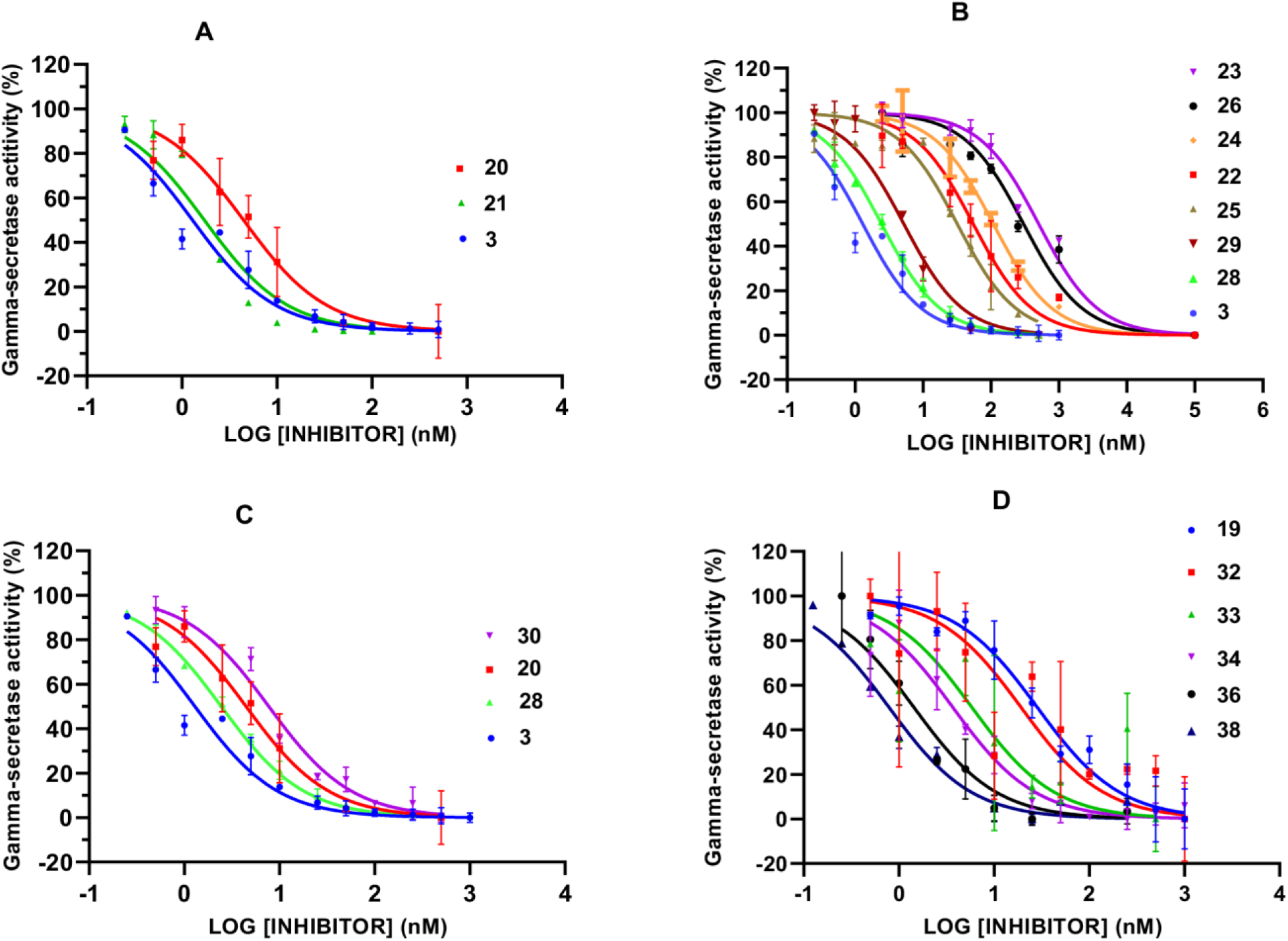
Concentration-dependent inhibition curves of selected HPI-TSA conjugates toward γ-secretase. **A**) Compound **20**, designed to trap the transition state of endoproteolysis (i.e., C-terminally extended with Lys-Lys-Lys), and comparable compounds without this extension. **B**) Compounds designed to capture helix unwinding (i.e., alkyl linker replaced by amino acids). **C**) Compound **30**, designed to capture both endoproteolysis and helix unwinding and comparable compounds. **D**) Compounds designed to capture lateral gating (i.e., compounds containing a D-HPI). All compounds were tested using 1 nM of purified enzyme and 0.5 μM of recombinant APP-based substrate C100Flag. IC_50_ values are shown in Tables 1 and 2.

**Figure 4.**
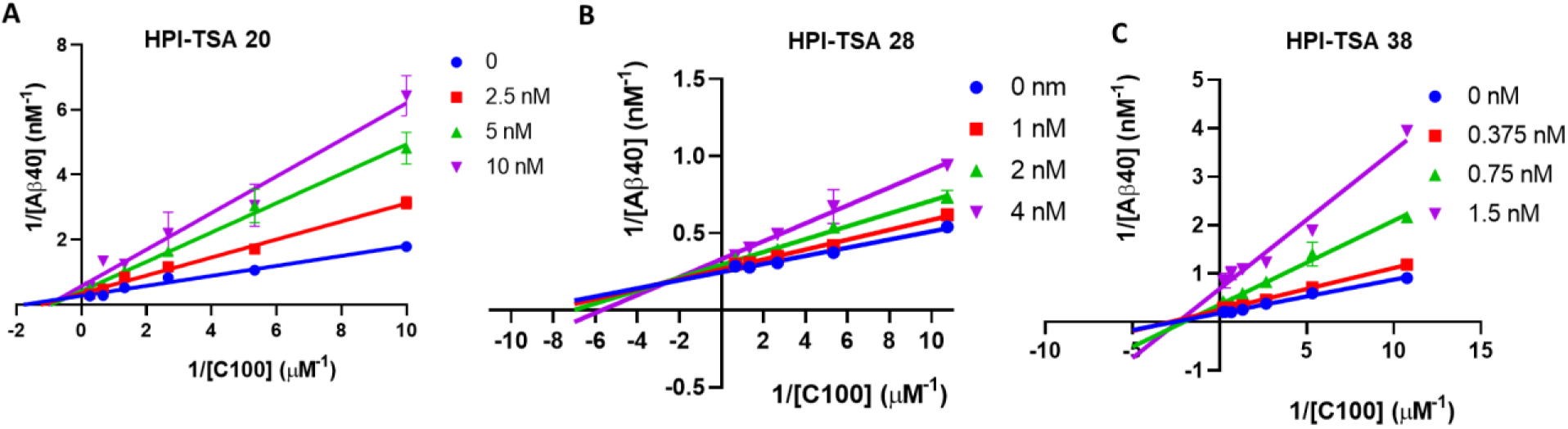
Inhibition constant (*K*_i_) determination experiments of top inhibitors of γ-secretase designed for each aspect of substrate recognition. **A**) Compound **20**, designed to trap the endoproteolysis transition state (*K*_i_ = 2.64 ± 0.18 nM). **B**) Compound **28**, designed to capture helix unwinding (*K*_i_ = 1.19 ± 0.10 nM). **C**) Compound **38**, designed to capture lateral gating (*K*_i_ = 0.65 ± 0.02 nM). A noncompetitive inhibition profile was seen for each compound.

### Structure-Activity Relationships

TSA inhibitor **19**, an optimized hydroxyethylurea pentapeptide analog spanning residues P2 through P3’,^20^ showed an IC_50_ value of 41 nM, while HPI **2**, a decapeptide containing helix-inducing α-aminoisobutyric acid (Aib) residues,^23^ showed comparable inhibitory activity at 58 nM (Table 1). The C-terminus of HPI **2** was coupled to the N-terminus of TSA **19** through a variable ω-aminoalkanoyl linker/spacer in generating full-TMD HPI-TSA conjugates. We previously reported that HPI-TSA **3**, with a 10-atom linker, is a virtual stoichiometric inhibitor of γ-secretase, with an IC_50_ of 0.8 nM in the presence of 1 nM enzyme and a *K*_i_ 0.42 ± 0.12 nM.^31^ Each component is critical for potent inhibitory activity: the helicity of the N-terminal HPI, the length of the linker, and the presence of transition-state-mimicking functionality of the C-terminal peptide analog. TMD mimetic **3** and related compounds containing a TSA moiety with only three P’ residues (P1’, P2’, P3’), should trap the protease at the transition state of carboxypeptidase trimming. Extending the C-terminus of the TMD mimics into P4’, P5’, P6’ should trap the transition state of endoproteolysis. The cryo-EM structure of enzyme bound to APP substrate shows juxtamembrane Lys-Lys-Lys residues P4’, P5’, and P6’ in an extended conformation and interacting with presenilin-1 (PSEN1).^28^ Extending conjugate **3** in this way led to **20** with an IC_50_ of 4.3 nM (Table 1). Although we had previously shown > twofold higher inhibitory potency with an N-terminal Boc versus acetyl group (cf. **3** and **21**, Table 1),^31^ conjugate **20** has acetyl at the N-terminus due to the need for Boc protection of the lysine side chains for SPPS. Extending the C-terminus of acetyl-capped conjugate **21** with three lysines as in conjugate **20**, led to a further > twofold reduction in potency (Table 1), suggesting that enhanced interactions with PSEN1 through the triple lysine extension may be counterbalanced by disrupted enzyme-inhibitor interactions elsewhere. Alternatively, the extended conformation of the juxtamembrane lysines of APP substrate and their interactions with PSEN1 observed in the reported cryo-EM structure might be less prominent at ambient temperature.^41^ In any event, **20** retained low-nanomolar potency (IC_50_ of 4.3 nM) and represents a new structural tool for trapping γ-secretase at the transition state of endoproteolysis. Inhibitor **20** showed ~9-fold and ~13-fold improved potency as compared to TSA **19** and HPI **2**, respectively, and is our optimized probe to trap endoproteolysis transition state. Along with capturing the transition state for proteolysis, this compound may lead to additional β-strand interactions with PSEN1, similar to that seen with γ-secretase bound to APP substrate.^28^ Recent structure elucidation of the protease bound to two non-transition inhibitors and former clinical candidates semagacestat and avagacestat revealed interaction with this same β-strand-forming region of PSEN1.^30^ As avagacestat has shown some selectivity for inhibiting γ-secretase cleavage of APP vis-à-vis Notch, a better understanding of the nature of this particular substrate-enzyme interaction may facilitate structure-based design of agents with improved selectivity for APP.

Next, we replaced the alkyl linker of the HPI-TSA conjugate **3** with the corresponding amino acid residues in the APP TMD. This region in substrate TMD is partially unwound in the bound substrate structures and located at the juncture of the helical and extended regions. Including this region in structural probes could therefore provide insight into the nature of helical substrate unwinding and setting up the transition state for proteolysis. The 10-atom ω-aminononanoyl spacer of **3** is close in length to the 9-atom backbone of a tripeptide. Thus, the linker was replaced with the corresponding three residues in the APP TMD: Val-Ile-Val (relative to Aβ48-type ε cleavage) and Ile-Val-Ile (relative to Aβ49-type ε cleavage). In addition, one or more residues of these tripeptide linkers was replaced with helix-destabilizing glycines, which can substantially increase endoproteolytic processing by γ-secretase when installed into APP substrate.^32^ Linker replacement of **3** with Val-Ile-Val (**22**) resulted in dramatic 74-fold reduction in potency (IC_50_ 59 nM) compared with **3** (IC_50_ 0.8 nM). Similarly, Ile-Val-Ile linker replacement (**25**) resulted in substantial 53-fold reduction in potency (IC_50_ of 42 nM). However, linker replacement with Ile-Val-Ile, with APP TMD residues relative to the Aβ49-generating ε cleavage site, was more potent than replacement with Val-Ile-Val, with residues relative to the Aβ48-generating ε site. This might reflect the preference for the Aβ49-generating ε cleavage site by the wild-type protease complex.^42^

One or two glycine replacements were made into conjugate **22** linked to Val-Ile-Val (**23** and **24**) or conjugate **25** linked to Ile-Val-Ile (**26** and **27**), and these replacements all resulted in further two-to-threefold loss of potency. However, alkyl linker replacement in **3** with Gly-Gly-Gly (**28**) provided the most potent tripeptide-linked HPI-TSA conjugate, with almost complete retention of potency (IC_50_ of **28** = 2.1 nM vs. IC_50_ of **3** = 0.8 nM). Low nanomolar potency by helix-destabilizing glycine linker replacements in **28** provides further support for a potential role of the corresponding region of APP substrate in TMD unwinding. The triple glycine replacement appears to be critical: replacement of the first or first and second Gly of **28** with Val and/or Ile (as in **23, 24, 26**, and **27**) led to substantial loss of inhibitory potency, and replacement of the third glycine with Ile (**29**) resulted in 2.7-fold reduced potency (IC_50_ = 5.7 nM). Since linker replacement with glycines resulted in potent HPI-TSA conjugates, we synthesized 30 with both C-terminal triple lysine extension and triple glycine linker replacement, which should serve as a probe for endoproteolysis as well as helix unwinding. Endoprotease probe **30** provided an IC_50_ of 7.6 nM, showing a similar ~twofold reduction in potency due to triple glycine replacement of the alkyl linker compared with **20** as seen between carboxypeptidase probes **3** vs. **28**. For conjugate **31**, in which the linker region with triple glycine was substituted with four glycine, the potency was statistically unchanged (IC_50_ of 7.7 nM for **30** vs. IC_50_ of 8.3 nM for **31**).

For probing the lateral gating pathway for substrate entry into the interior of γ-secretase, a 10-residue D-HPI **32** (Table 2) was coupled to TSA **19** through ω-aminoalkanoyl linkers of varying lengths. We previously showed that Aib-containing helical D-peptides such as **32** are also potent inhibitors of γ-secretase.^23, 24^ Originally designed as controls for the helical L-peptide inhibitors, some D-HPIs were even more potent than the most potent L-HPI. Helical disruption through inversion of two internal stereocenters substantially reduces inhibitory potency.^23^ Although adopting left-handed helicity, D-HPIs presumably present an array of hydrophobic side chains for initial interaction with γ-secretase that is similar to that presented by L-HPIs. However, the opposite helical handedness may prevent lateral gating, as entry of TMD substrate into the internal active site of PSEN1 involves helix unwinding.^28, 29^ Thus, D-HPIs may dock to the outside of the γ-secretase complex but not be drawn inside PSEN1, thereby allowing identification of the site of lateral gating. Like L-HPI **2**, D-HPI **32** is ten residues in length, with Aib residues in corresponding positions, and this compound inhibited purified γ-secretase under our assay conditions with an IC_50_ of 35 nM (Table 2). The new conjugate **33** with D-HPI **32** directly coupled with TSA **19**, displayed an IC_50_ of 7.5 nM, substantially more potent than either **19** or **32** alone.

As with the L-HPI series of conjugates,^31^ we synthesized a series of D-HPI-TSA conjugates with ω-aminoalkanoyl linkers of varying lengths. Similar to what was observed with the L-HPI-TSA series, inhibitory activity improved with increasing linker length and the conjugates **34-39** containing 4-, 6-, 8-, 10-, 12-, and 13-atom spacers displayed IC_50_ values of 6.0, 3.2, 1.5, 0.70 and 0.87 respectively. In this series, the 12-atom spacer (**38**) was optimal, showing stoichiometric inhibition (IC_50_ of 0.70 nM with 1 nM enzyme), as compared to a 10-atom spacer for L-peptidomimetics. The increased optimal length compared with L-HPI-TSAs may be due to the inability of the D-HPI moiety to enter into the interior of PSEN1. On the other hand, in the D-HPI-TSA series inhibitor potency is less dependent on the linker length compared with what we previously observed in the L-HPI-TSA series. Direct coupling of D-HPI to TSA (i.e., with no linker) led to substantially increased potency (Table 2, cf. TSA **19** and D-HPI **32** versus D-HPI-TSA **33**), while direct coupling of L-HPI to TSA did not increase potency at all.^31^ Instead of binding to an external docking exosite, the D-HPI may be bound to the same internal site as L-HPI but in a different orientation, one that allows the TSA component to bind to the active site when no linker is present.

Experiments to determine enzyme inhibition constants (*K*_i_ values) were carried out for the most potent HPI-TSA conjugates in each new class (**20, 28** and **38**)(Figure 4). *K*_i_ values for **20, 28** and **38** were 2.64, 1.19 and 0.65 nM, respectively, and all three conjugates exhibited non-competitive inhibition, as seen with prototype HPI-TSA **3** ^31^ as well as HPI **2** and TSA **1** on their own.^43^

### Inhibitor Competition Experiments

To verify that conjugates **28** and **38**—with linker and HPI replacements, respectively—occupy both the substrate-binding exosite as well as the active site, we performed competition experiments with biotinylated photoaffinity probes for γ-secretase.^25, 44^ Specifically, we used active-site-directed photoprobe TSA-Bpa-Bt and exosite-directed photoprobe HPI-Bpa-Bt (Fig. 5), in which each probe contains a photoactivatable 4-benzoyl-L-phenylalanine for covalent crosslinking and a linker-biotin moiety for isolation of photo-crosslinked proteins. Incubation of each probe with membranes isolated from HEK 293 cells overexpressing γ-secretase in the presence and absence of competitor was followed by irradiation at 350 nM. Labelled proteins were pulled down with streptavidin beads, and PSEN1 NTF subunit was detected by western blot. As previously demonstrated,^25^ TSA **1** but not HPI **2** inhibited labeling of PSEN1 NTF by TSA-Bpa-Bt, whereas HPI **2** but not TSA **1** inhibited labeling by HPI-Bpa-Bt. In this way, the distinct binding sites for HPIs and TSAs on PSEN1 was confirmed (Fig. 5). HPI-TSA conjugates **28** and **38** decreased labelling of PS1 NTF by both photoprobes, indicating that these conjugates occupy both the active site and exosite on γ-secretase.

**Figure 5.**
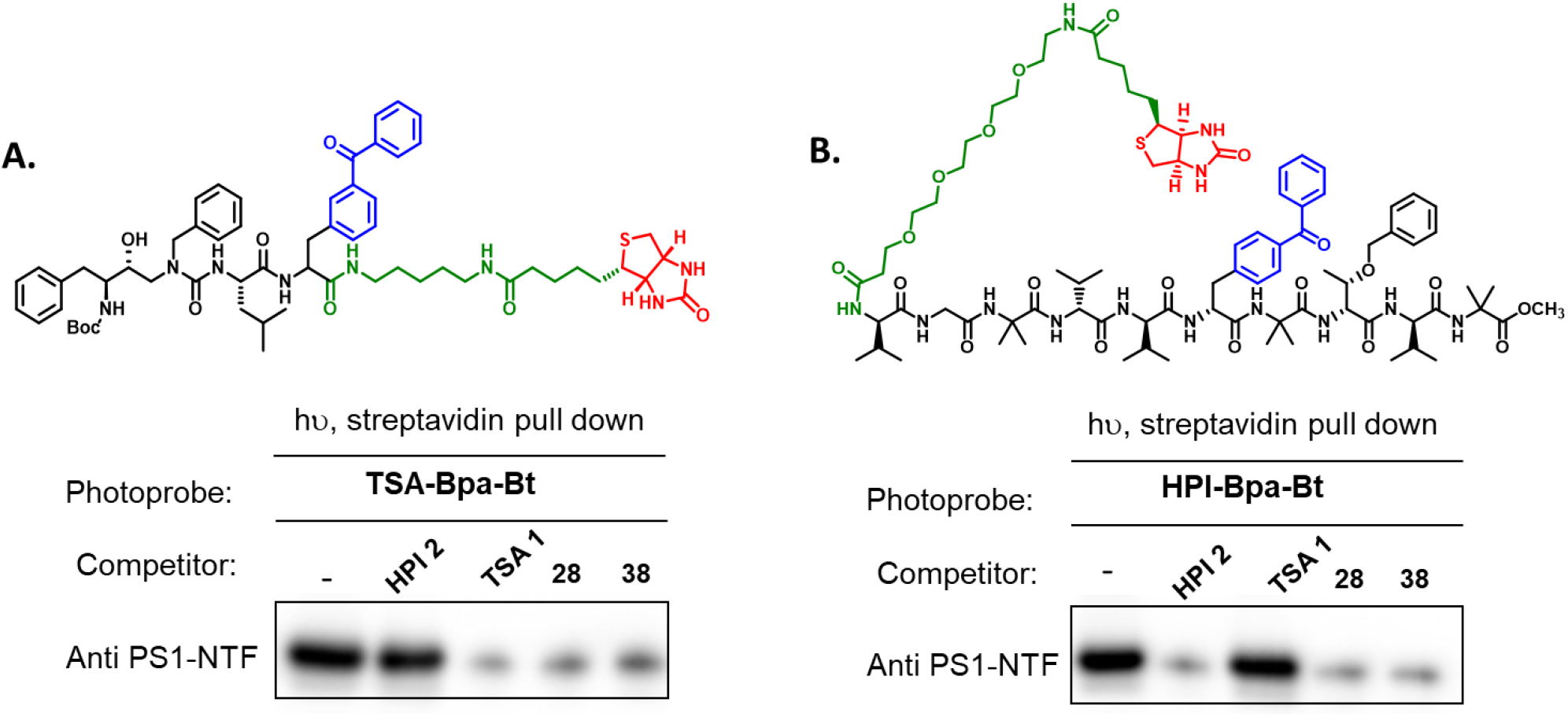
Competition of HPI-TSAs conjugates with photoaffinity probes for γ-secretase. Photoprobes TSA-Bpa-Bt (left) and HPI-Bpa-Bt (right) covalently label presenilin-1 (PS1) N-terminal fragment (NTF) at the active site and docking site, respectively, as shown through streptavidin pulldown and Western blotting. (**A**) TSA (**1**) but not HPI (**2**) decreased labelling of PSEN1 NTF by the TSA photoprobe. HPI-TSA conjugates **22** and **38** decreased labelling by this TSA photoprobe. (**B**) HPI (**2**) but not TSA (**1**) decreased labelling PS1 NTF by the HPI photoprobe. HPI-TSA conjugates **28** and **38** decreased labelling by this HPI photoprobe as well.

To verify again that the conjugate **38**—with replacement of L-HPI with D-HPI—occupy both the substrate-binding exosite as well as the active site, we performed enzyme inhibitor cross-competition experiments as previously reported.^31, 43^ Specifically, we used active-site-directed inhibitor TSA **1** and exosite-directed inhibitor HPI **2** as competitors of D-HPI-TSA **38**. D-HPI-TSA **38** could compete with both TSA **1** and HPI **2** for binding, as indicated by the parallel lines in Figure 6. In contrast, inhibitors **1** and **2**, which bind to distinct sites, do not compete with each other in such an experiment, as lines converge near the x-axis.^31^ These results, like those from the photoaffinity crosslinking experiments, suggest that **38** occupies both exosite and active site on the enzyme. Nevertheless, because direct coupling of D-HPI to TSA (i.e., with no linker) leads to substantially increased potency, it remains unclear whether the occupied exosite is an external docking site for lateral gating or an internal site contiguous with the active site.

**Figure 6.**
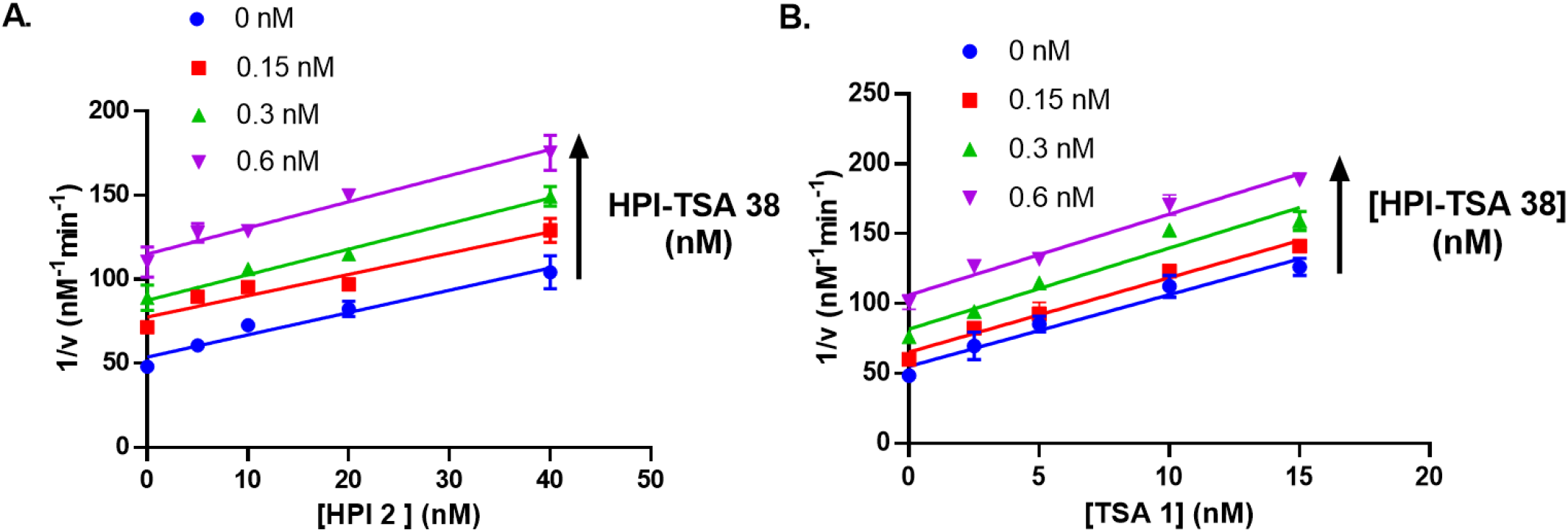
Cross-competition kinetic experiments between two inhibitors. (**A**) HPI **2** and HPI-TSA **38** at [**38**] = 0, 0.15, 0.30, and 0.60 nM. (**B**) TSA **1** and HPI-TSA **38** at [**38**] = 0, 0.15, 0.30, and 0.60 nM. Parallel lines indicate cross-competition, and HPI-TSA **38** competes with both TSA **1** and HPI **2**.

## Conclusions

We have developed hybrid peptidomimetic inhibitors of the γ-secretase complex, mimicking the entire substrate TMD and occupying both the protease active site and exosite. These TMD mimics are designed to trap the active enzyme in a conformation poised for intramembrane proteolysis. The prototype conjugate **3** is a stoichiometric inhibitor of γ-secretase designed to trap the protease at the transition state of carboxypeptidase trimming^31^ (i.e., poised to trim off a tripeptide from the C-terminus of intermediate substrate) for structure elucidation by cryo-EM, as the TSA component has only 3 P’ residues. Here we have developed next-generation structural probes for γ-secretase, which are designed to trap the enzyme in different stages of substrate recognition and proteolytic processing. In addition to the carboxypeptidase-trapping probe **3**, related probes have been developed to trap the endoprotease transition state, substrate TMD helix unwinding and substrate lateral gating. For the design of endopeptidase probe, the C-terminus of TMD mimetic **3** was extended with three amino acids from P4’ to P6’, three lysines that represent the cytosolic juxtamembrane region of APP substrate. Probes for helix unwinding have the ten-atom alkyl linker region of **3** replaced with three amino acids, with helix-disrupting glycines providing the most potent inhibition. Lateral gating probes have the L-HPI moiety of **3** replaced with D-HPI, which may prevent helix unwinding and entry into the interior of the protease. These conjugates have been produced via solid- and solution-phase peptide synthesis involving natural amino acids and a tri-peptidomimetic building block accessed in a multi-step protocol in a moderate yields. These highly hydrophobic TMD-mimetic compounds were purified to >95%, and inhibitory potencies were determined using purified enzyme expressed from a tetracistronic construct in suspension HEK 293 cells. At least one compound in each new series (C-terminal extension, linker replacement, and D-HPI replacement) retained low- or subnanomolar inhibitory activities toward γ-secretase, comparable to that of HPI-TSA prototype conjugate **3**. These potent peptidomimetics exhibit non-competitive inhibition profiles and occupy both exosite and active site of the enzyme, as demonstrated by the photo-affinity labelling experiments and enzyme-inhibitor cross competition assays.

We expect that each of these potent peptidomimetics will be conducive to co-structure elucidation with γ-secretase by cryo-EM and that each optimized conjugate will trap the protease complex in a different conformational state. Preliminary cryo-EM experiments in collaboration with the laboratory of Yigong Shi of Tsinghua University show each type of HPI-TSA conjugate bound to the γ-secretase complex in a manner generally similar to that seen with APP substrate crosslinked to inactive enzyme. A high-resolution structure of one compound (**28**) shows the hydroxyl group of the TSA moiety coordinated with the two catalytic aspartates on PSEN1, trapping the enzyme at the transition state, poised as it would be for intramembrane proteolysis of TMD substrate (Y. Shi, R. Zhou, and G. Yang, unpublished results). Further refinement of the structures of each probe type bound to γ-secretase is in progress and will be reported in due course. These structures should provide information on substrate recognition, entry and processing as well as facilitate drug design for Alzheimer therapeutics (e.g., to identify APP substrate-selective inhibitors or modulators). Understanding of structural mechanisms of substrate interaction and processing by γ-secretase may also help elucidate how Alzheimer-causing mutations in APP and PSEN1 lead to deficient carboxypeptidase trimming of long Aβ intermediates.^33–35^ More broadly, the approach of developing substrate-based inhibitors as structural probes may have wider applicability in trapping active conformations of other large enzyme complexes (e.g., the spliceosome) for cryo-EM analysis.

## Experimental Section

### General

Chemicals used for synthesis and biochemical experiments were commercially obtained from various vendors (Fischer, Acros, TCI America, Sigma-Aldrich, Chem-Impex International). The purity of all starting materials and solvents used in the synthesis was > 95% and used without further purification or drying. Chemical reactions toward the synthesis of tripeptidomimetic building block **11** were monitored via thin layer chromatography (TLC) using aluminum sheets with silica gel 60 F_254_ (Merck). Silica gel 0.060-0.200 mm, pore diameter ca. 6 nm was employed for column chromatography. LC/ESI-MS (HPLC analysis coupled to electrospray ionization mass spectrometry) was used to determine the purity of the final compounds, which was found to be > 95% in all cases. For LC/ESI-MS measurement, samples were prepared by dissolving 1 mg/mL of compound in H_2_O/MeOH (1:1) containing 2 mM ammonium acetate, 10 μL of which was injected for HPLC analysis, eluting with a gradient of water/methanol (containing 2 mM ammonium acetate) from 90:10 to 0:100 for 15 min at a flow rate of 250 μL/min. LC-MS analysis was performed using a Waters Analytical System-Acquity HPLC with an APCI mass spectrometer; UV absorption was detected using an Acquity diode array detector. ^1^H and ^13^C NMR spectra were performed on a Bruker Avance 500 and 400 MHz spectrometer and were recorded at ambient temperature using either DMSO-*d*_*6*_, or CDCl_3_ as solvent. High-resolution mass spectroscopy (HRMS) analysis was recorded on a LCT Premier mass spectrometer (Micromass Ltd., Manchester, UK), a quadrupole and time-of-flight tandem mass analyzer with an electrospray ion source. Solid phase synthesis of peptides were carried out using standard chemistry with Fmoc-protected amino acids using an Aapptec Focus XC peptide synthesizer. Lyophilization was carried out using a Labconco FreeZone 4.5 Liter Benchtop Freeze Dry Systems. For peptide purification, a preparative HPLC system (2545 Quaternary Gradient Module) from Waters was used with XBridge Peptide BEH C18, 300Å column. Melting points were measured on Mel-Temp Digital apparatus and are reported without correction.

Altogether, 20 different peptidomimetics derivatives (HPI-TSA conjugates) were synthesized and investigated in this study, and all are new compounds not previously described in literature. The procedure for synthesis of tripeptidomimetic building block **11** is described below. Two different analytical methods were used to control the purity of peptides as described in Table 3. The purity of all final HPI-TSA conjugates were determined to be >95%.

**Table 3.** Measured mass, HPLC peak retention time and LC-MS purity of tested peptides

| Compound | Mass calc. | Mass measured (ESI) | Mass measured (TOF) | Retention time (Method A)<sup>a</sup> | Retention time (Method B)<sup>b</sup> | LC-MS % Purity |
| --- | --- | --- | --- | --- | --- | --- |
|  |  | M+Na<sup>+</sup> | M+H<sup>+</sup> | min | min |  |
| **20** | 2174.4323 | 2197.4341 | 2175.4401 | 10.33 | 10.35 | 99 |
| **21** | 1790.1474 | 1813.1342 | 1791.1542 | 7.86 | 8.55 | 95 |
| **22** | 2004.2791 | 2027.2632 | 2005.2868 | 10.98 | 10.99 | 96 |
| **23** | 1962.2322 | 1985.2225 | 1963.2391 | 8.40 | 9.87 | 97 |
| **24** | 1906.1696 | 1929.1540 | 1907.1725 | 6.28 | 6.46 | 96 |
| **25** | 2018.2948 | 2041.2835 | 2019.3027 | 10.99 | 6.48 | 96 |
| **26** | 1962.2322 | 1985.2178 | 1963.2386 | 6.99 | 7.19 | 98 |
| **27** | 1920.1802 | 1943.2187 | 1921.1803 | 5.19 | 4.78 | 96 |
| **28** | 1864.1226 | 1887.1153 | 1865.1284 | 8.03 | 8.10 | 96 |
| **29** | 1920.1852 | 1943.1650 | 1921.1939 | 8.55 | 8.54 | 98 |
| **30** | 2190.3657 | 2213.3653 | 2191.3842 | 7.31 | 7.14 | 95 |
| **31** | 2247.3871 | 2270.3864 | 2248.3576 | 11.06 | 11.04 | 97 |
| **33** | 1737.0845 | 1760.0735 | 1738.1045 | 10.49 | 11.58 | 99 |
| **34** | 1808.1216 | 1831.1128 | 1809.1274 | 9.52 | 9.81 | 97 |
| **35** | 1836.1529 | 1859.1460 | 1837.1654 | 9.30 | 9.58 | 99 |
| **36** | 1849.1733 | 1872.1361 | 1850.1381 | 8.50 | 8.40 | 98 |
| **37** | 1892.2155 | 1915.2350 | 1893.2325 | 8.78 | 8.89 | 96 |
| **38** | 1920.2468 | 1943.2636 | 1921.2845 | 8.24 | 8.67 | 98 |
| **39** | 1934.2624 | 1957.2225 | 1935.2407 | 8.35 | 8.85 | 99 |

<sup>a</sup>gradient of water/methanol: 2-propanol (1: 1) (containing 0.02% formic acid) from 60:40 to 0:100
<sup>b</sup>gradient of water/acetonitrile (containing 0.05% trifluoroacetic acid) from 50:40 to 0:10

### Solid- and solution-phase synthesis of peptides and peptidomimetics inhibitors

TSAs **1** and **19** and HPIs **2** and **32** were synthesized as previously described.^22–24, 31^ Biotinylated peptidomimetic inhibitors TSA-Bpa-Bt and HPI-Bpa-Bt used for photoaffinity labeling experiments were synthesized as previously described.^25^ All HPI-TSA conjugates used in this study were synthesized in an automated solid-phase synthesizer (Focus XC, Aapptec LLC) on Rink amide resin using Fmoc-chemistry. N-acetylated HPI-TSAs (**20, 21, 30 and 31**) were iteratively double-coupled to full length using building block **5** and Fmoc-protected amino acids and finally acetylated before cleaving from the resin (Scheme 2). N-Boc HPI-TSAs were iteratively double-coupled to the penultimate length, then cleaved from the resin. These peptidomimetic intermediates were purified by HPLC and further coupled in solution phase with N-Boc-L-valine (Scheme 2). Note that the *O*-TBDMS protecting group in building block **11** was removed during cleavage of peptides from the Rink amide resin, as the cleavage cocktail contains TFA.

### Procedure for the synthesis of tripeptidomimicking building block 11

#### *tert*-Butyl ((2*S*,3*R*)-4-(benzylamino)-3-hydroxy-1-phenylbutan-2-yl)carbamate (5)

To a 5.0 M solution of oxirane **4** (1 eq.) in 2-propanol (5 mL) was added 20 eq. of benzylamine, and the reaction was refluxed under dry N_2_ for 20 h. The reaction mixture was allowed to cool to room temperature, diluted with ethyl acetate and washed consecutively with water, aqueous 1 N HCl and saturated NaHCO_3_ solution. The organic phase was dried over anhydrous MgSO_4_, concentrated, and pure **2** was obtained by precipitation with hexane, filtration and drying.

#### (*S*)-(-)-2-isocyanato-4-methylvaleric acid methyl ester

The isocyanates of *L*-leucine was obtained by stirring the HCl salt of the corresponding amino ester (20 mmol) in a mixture of CH_2_Cl_2_ (10 mL) and saturated NaHCO_3_ solution (10 mL) for 20 min in a three-necked flask followed by addition of triphosgene (20 mmol) in a single portion. After being stirred for 1 h, the reaction mixture was poured into a beaker containing ice and stirred for 20 min. The mixture was then extracted with 3 × 50 mL CH_2_Cl_2_. The combined organic fraction was dried over MgSO_4_, filtered, and evaporated to a colorless oil. Then the colourless oil was purified was purified using flash chromatography (Methanol: CH_2_Cl_2_ 1:50) and a trituration with ethyl acetate and hexane provided a white solid.

#### Methyl (benzyl((2*R*,3*S*)-3-((tert-butoxycarbonyl)amino)-2-hydroxy-4-phenylbutyl)carbam-oyl)-*L*-leucinate (6)

To a solution of hydroxyethylamine **5** (2.0 g, 5.40 mmol) in CH_2_Cl_2_ (3 mL) was added (*S*)-(-)-2-isocyanato-4-methylvaleric acid methyl ester (1 eq., 935 mg, 5.40 mmol) in CH_2_Cl_2_ (3 mL) at 0 °C for 30 min. After being stirred at room temperature for 6 h, the reaction mixture was concentrated, and the hydroxyethylurea **6** was purified using flash chromatography (Methanol:CH_2_Cl_2_ 1:49) as a white solid (2.7 g, 93% yield). ^1^H NMR (400 MHz, CDCl_3_) *δ* 7.28 – 7.10 (m, 10H), 5.41 (s, 1H), 4.75 (s, 1H), 4.51 (dd, *J* = 12.4 Hz, 2H), 4.36 (m, 2H), 3.68 (m, 1H), 3.65 (s, 3H), 3.64 (m, 1H), 3.52 (m, 1H), 3.15 (m, 1H), 2.88 (m, 2H), 1.49 (m, 2H), 1.39 (m, 1H), 1.21 (s, 9H), 0.82 (m, 6H). ^13^C NMR (125 MHz, CDCl_3_) *δ* 175.13, 159.37, 155.82, 137.28, 137.06, 129.48, 128.89, 128.42, 127.64, 127.11, 126.39, 73.64, 54.55, 53.82, 52.66, 52.24, 42.09, 35.87, 28.24, 24.87, 22.89, 21.81. LC-MS (*m*/*z*): negative mode 540 [M-H]^−^, positive mode 542 [M+H]^+^. Purity by HPLC-UV (214 nm)-ESI-MS: 98.60%. mp 175-177 °C.

#### Methyl (((2*R*,3*S*)-3-amino-2-hydroxy-4-phenylbutyl)(benzyl)carbamoyl)-*L*-leucinate (7)

To a solution of hydroxyethylurea **6** (2.0 g) in CH_2_Cl_2_ (7 mL) was added trifluoroacetic acid (3 mL). After being stirred at r.t. for 2 h, the reaction mixture was quenched by adjusting to pH 7 with saturated aq. sodium bicarbonate. Extraction with CH_2_Cl_2_ (2 × 50 mL) was followed by washing with water and additional extraction with CH_2_Cl_2_ (2 × 50 mL). All organic fractions were pooled, dried over anhydrous MgSO_4_ and concentrated to 1.58 g (99%) of pure product. ^1^H NMR (400 MHz, CDCl_3_) *δ* 7.38 –7.17 (m, 10H), 4.75 (d, 1H), 4.44 (m, 2H), 3.72 (d, 1H), 3.70 (s, 3H), 3.67 (m, 2H), 3.34 (m, 1H), 3.05 (m, 1H), 2.95 (m, 1H), 2.51 (m, 1H), 1.66 – 1.42 (m, 3H), 0.92 (d, 3H), 0.91 (d, 3H). ^13^C NMR (125 MHz, CDCl_3_) *δ* 175.27, 159.94, 138.60, 137.67, 135.35, 128.82, 128.63, 127.50, 127.31, 126.50, 74.44, 55.35, 52.60, 52.12, 51.69, 50.91, 41.29, 39.04, 24.94, 22.91, 21.85. LC-MS (*m*/*z*): negative mode 440 [M-H]^−^, positive mode 442 [M+H]^+^. Purity by HPLC-UV (214 nm)-ESI-MS: 99.00%. mp 158–160 °C.

#### Methyl(((2*R*,3*S*)-3-((((9H-fluoren-9-yl)methoxy)carbonyl)amino)-2-hydroxy-4-phenylbutyl)-(benzyl)carbamoyl)-*L*-leucinate (8)

To a solution of hydroxyethylurea **7** (1.0 g, 2.27 mmol) in CH_2_Cl_2_ (5 mL) under dry N_2_, was added 1.2 eq. of Fmoc chloride and 2 eq. of DIPEA. The reaction mixture was stirred overnight at r.t. and concentrated *in vacuo*. Silica gel chromatography (0 to 5% MeOH in CH_2_Cl_2_) provided 1.5 g (91%) of **8** as white powders. ^1^H NMR (400 MHz, CDCl_3_) δ 7.78 (d, *J* = 7.6 Hz, 2H), 7.50 (dd, *J* = 7.5 Hz, 2H), 7.45 – 7.38 (m, 2H), 7.34 –7.20 (m, 10H), 7.16 (d, *J* = 7.4 Hz, 2H), 4.79 (d, *J* = 9.0 Hz, 2H), 4.55 (m, 3H), 4.43 (m, 1H), 4.27 (dd, *J* = 10.7 Hz, 1H), 4.13 (m, 1H), 3.84 (m, 1H), 3.75 (s, 3H), 3.66 (m, 1H), 3.22 (d, *J* = 14.9 Hz, 1H), 2.98 –2.84 (m, 2H), 1.56 (m,2H), 1.45 (m, 1H), 0.92 (d, 3H), 0.90 (d, 3H). ^13^C NMR (125 MHz, CDCl3) *δ* 175.07, 159.89, 156.14, 143.89, 141.34, 137.43, 129.50, 128.95, 128.50, 127.70, 127.04, 126.53, 124.98, 119.96, 73.30, 66.52, 54.98, 52.55, 52.29, 53.98, 51.97, 47.23, 41.32, 35.48, 31.60, 24.87, 22.89, 21.79. LC-MS (*m*/*z*): negative mode 662 [M-H]^−^, positive mode 664 [M+H]^+^. Purity by HPLC-UV (214 nm)-ESI-MS: 99.60%. mp 183-185 °C.

#### Methyl(((2R,3S)-3-((((9H-fluoren-9-yl)methoxy)carbonyl)amino)-2-((tert-butyldimethylsilyl)-oxy)-4-phenylbutyl)(benzyl)carbamoyl)-*L*-leucinate (9)

To a stirred solution of Fmoc-protected hydroxyethylurea **8** (1.0 g, 1.50 mmol) in DMF (3 mL) under dry N_2_ at r.t. was added 2 eq. of TBDMS chloride and 1.2 eq. DMAP. After 36 h, the reaction mixture was quenched with water, extracted with 3 × 100 mL of ethyl acetate, dried under anhydrous MgSO_4_ and concentrated *in vacuo*. The resultant oil was purified by silica gel chromatography (0 to 2 % MeOH in CH_2_Cl_2_) to obtain 880 mg of **9** (75% yield) as white powders. ^1^H NMR (400 MHz, CDCl_3_) *δ* 7.73 (d, *J* = 7.7 Hz, 2H), 7.45 (dd, *J* = 7.6 Hz, 2H), 7.36 (m, 2H), 7.30 – 7.20 (m, 10H), 7.16 (d, *J* = 7.4 Hz, 2H), 4.71 (m, 1H), 4.45 (m, 3H), 4.03 (m, 1H), 3.67 (m, 3H), 3.58 (s, 3H), 3.48 (m, 1H), 3.26 – 3.12 (m, 1H), 2.99 (m, 1H), 2.85 (m, 1H), 1.41(m, 1H), 1.23 (m, 2H), 0.89 (s, 9H), 0.86 (s, 3H), 0.83 (s, 3H), 0.03 (s, 3H), 0.00 (s, 3H). ^13^C NMR (125 MHz, CDCl_3_) *δ* 174.72, 159.89, 158.65, 141.34, 139.26, 135.99, 129.50, 129.16, 128.68, 127.38, 126.47, 119.96, 74.92, 66.52, 55.93, 55.31, 52.56, 52.27, 51.94, 49.99. 47.24, 41.47, 31.60, 25.94, 22.84, 21.96, 17.98, −4.25, −4.87. LC-MS (*m*/*z*): negative mode 777 [M-H]^−^, positive mode 779 [M+H]^+^. Purity by HPLC-UV (214 nm)-ESI-MS: 99%. mp 193-195 °C.

#### (((2*R*,3*S*)-3-amino-2-((tert-butyldimethylsilyl)oxy)-4-phenylbutyl)(benzyl)carbamoyl)-*L*-leucine (10)

To a solution of *N*-Fmoc-hydroxyethylurea **9** (1.0 g, 1.28 mmol) in THF (5 mL) was added 5 mL of 1.0 M aqueous LiOH and the reaction was stirred for 4 h. After addition of brine (20 mL), the aqueous solution was extracted with 3 × 100 mL of ethyl acetate, dried over anhydrous MgSO_4_ and concentrated to give 580 mg of **10** (yield 85%)as white solids. ^1^H NMR (400 MHz, CDCl3) *δ* 7.31 – 7.15 (m, 10H), 4.70 (d, *J* = 7.5 Hz, 1H), 4.45 – 4.34 (m, 1H), 4.09 (m, 1H), 3.70 (m, 1H), 3.50 (m, 2H), 3.18 (m, 1H), 2.99 (m, 1H), 2.32 (m, 1H), 1.84 (m, 1H), 1.70 (m, 1H), 0.94 (d, *J* = 6.6 Hz, 3H), 0.89 (m, 12H), 0.03 (s, 3H), 0.00 (s, 3H), 1.02 (s, 3H). ^13^C NMR (125 MHz, CDCl3) *δ* 174.19, 155.77, 139.26, 138.07, 129.15, 128.67, 128.56, 127.40, 127.13, 126.40, 74.59, 55.84, 52.55, 51.83, 48.98, 41.47, 34.67, 31.60, 25.90, 22.62, 14.12, −4.26, −4.88. LC-MS (*m*/*z*): negative mode 540 [M-H]^−^, positive mode 542 [M+H]^+^. Purity by HPLC-UV (214 nm)-ESI-MS: 99.00%.

#### (((2*R*,3*S*)-3-((((9H-fluoren-9-yl)methoxy)carbonyl)amino)-2-((tert-butyldimethylsilyl)oxy)-4-phenylbutyl)(benzyl)carbamoyl)-L-leucine (11)

To a solution of Hydroxyethylurea **10** (0.5 g, 0.92 mmol) in CH_2_Cl_2_ (5 mL) under dry N_2_ was added 1.2 eq. of Fmoc chloride and 2 eq. of DIPEA, and the reaction mixture was stirred overnight at room temperature. After being concentrated *in vacuo*, the mixture was eluted through silica gel using 0 to 5 % MeOH in CH_2_Cl_2_ to give 423 mg of titled compound (yield 60%) as white powders. ^1^H NMR (400 MHz, CDCl_3_) *δ* 7.74 (m, 2H), 7.46 (m, 2H), 7.39 – 7.27 (m, 4H), 7.28 – 7.19 (m, 10H), 5.07 (m, 1H), 4.52 (m, 2H), 4.39 (dd, *J* = 15.5 Hz, 2H), 4.28 (m, 3H), 4.08 (m, 1H), 3.85 (bs, 1H), 3.66 (m, 2H), 3.47 (m, 1H), 3.18 (m, 1H), 3.04 (m, 1H), 1.74 (m, 1H), 1.47 (m, 1H), 1.33 (m, 1H), 0.94 (m, 3H), 0.91 (s, 3H), 0.83 (s, 6H), 0.03 (s, 3H), 0.00 (s, 3H). ^13^C NMR (125 MHz, CDCl_3_) *δ* 174.74, 160.12, 155.21, 146.02, 139.30, 138.05, 128.67, 128.57, 127.81, 127.38, 127.13, 126.48, 123.96, 120.07, 74.94, 68.81, 55.92, 52.54, 51.84, 45.96, 41.47, 25.94, 24.95, 22.84, 17.88, −4.25, −4.87. LC-MS (*m*/*z*): negative mode 763 [M-H]^−^, positive mode 765 [M+H]^+^. Purity by HPLC-UV (214 nm)-ESI-MS: 97.20%. mp 186-188 °C.

### Protein overexpression and purification

C100-Flag construct in the pET22b vector was transformed and expressed into *E. coli* BL21 cells.^39^ *E. coli* BL21 cells were grown in LB media at 37 °C in an incubator shaker until OD_600_ reached 0.6. Cells were induced with 0.5 mM IPTG and was grown for 4 hours shaking. Cells were then pelleted by centrifugation and were resuspended in lysis buffer composed of 50 mM HEPES pH 8, 1% Triton X-100. Cells were lysed by French press three times, and lysate was centrifuged to remove the cell debris. The clear lysate was mixed with anti-FLAG M2-agarose beads (Sigma-Aldrich) for 16 hours at 4 °C. Beads were washed with lysis buffer three times before elution with 100 mM glycine pH 2.5, 0.25% NP-40 detergent. The eluate was neutralized with Tris HCl and stored at −80 °C.

The γ-secretase complex was expressed in suspension human embryonic kidney (HEK) 293 cells from a pMLINK tetracistronic vector (courtesy of Y. Shi, Tsinghua University, Beijing).^45^ HEK293 cells were cultured in Freestyle 293 media (Life Technologies), shaking at 125 rpm while incubating at 37 oC with 8% CO_2_ until a density 2 × 10^6^ cells/ml was reached. For transfection, media was replaced with fresh Freestyle 293 media. 150 μg pMLINK tetracistronic vector and 450 μg polyethylenimines of 25 kDa (PEI) was mixed in 5 mL of Freestyle media and incubated for 30 min at room temperature. The DNA/PEI mixture was added to the cell culture, which after 24 h was subcultured in 3 flasks and further grown for 36 h before harvesting. Cells were pelleted, resuspended in buffer consisting of 50 mM MES pH 6.0, 150 mM NaCl, 5 mM CaCl_2_ and 5 mM MgCl_2_, and lysed via French Press by passing twice through a French press. The lysate was pelleted by centrifugation at 3000 x g for 10 minutes. The supernatant was ultracentrifuged at 100,000 x g for 1 h to isolate the membrane pellet. The membrane pellet was washed with 0.1 M sodium bicarbonate pH 11.3 by successive passage through 18, 22, 25 and 27 gauge needles. The resulting solution was incubated on ice for 20 min, followed by ultracentrifugation at 100,000 x g for 1 h. The pellet was resuspended in 50 mM HEPES pH 7, 150 mM NaCl, 1% CHAPSO and incubated in ice for 1 h, followed by ultracentrifugation at 1000,000 X g. The supernatant was mixed with anti-FLAG M2-agarose beads (Sigma-Aldrich) and TBS with 0.1% digitonin and incubated for 16 h at 4 °C. Beads were washed three times with TBS/0.1% digitonin mixture before elution of the γ-secretase complex with buffer consisting of 0.2 mg/ml FLAG peptide in TBS/ 0.1% digitonin. The resulting eluate was stored at −80 °C.

### γ-Secretase activity assays

For IC_50_ determination, 1 nM purified γ-secretase in standard assay buffer^*5*^ (50 mM HEPES, pH 7.0, 150 mM NaCl, 0.025% DOPC (18:1 (Δ9-Cis) phosphatidylcholine, Avanti Polar Lipids), 0.1% DOPE (18:1 (Δ9-Cis) phosphatidylethanolamine, Avanti Polar Lipids), and 0.25% CHAPSO (Sigma-Aldrich) was incubated at 37 °C for 30 min before addition of varying concentrations of inhibitor in DMSO (2% final DMSO concentration) and 0.5 μM C100Flag substrate. The enzyme reaction mixture was incubated for 2 h at 37 °C, and major proteolytic product Aβ40 was determined by a sensitive and specific sandwich ELISA (Thermo-Fisher). Sigmoidal curves were generated from inhibition data using GraphPad Prism 9.0. For *K*_i_ determination, enzyme reactions were run as above, but with varying concentrations of C100Flag (0-4.5 μM) and inhibitor. Data were fit to a noncompetitive inhibition equation also using GraphPad Prism 9.0.

Peptide IC_50_ inhibition was determined by using the following equation:

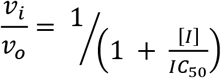

Where *v*_*i*_ and *v*_*o*_ are the initial velocity in the presence and absence, respectively, of inhibitor at concentration [*I*].

*K*_i_ values for noncompetitive inhibition were determined using the following equation:

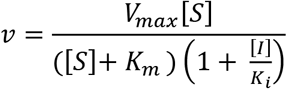

where, v is the initial rate, *S* is the substrate concentration, *K*_*i*_ is the dissociation constant for inhibitor.

### Photoaffinity labeling experiments

γ-secretase (2 nM) in standard assay buffer was incubated at 37 °C for 30 min, whereupon 1 μM of biotinylated photoaffinity probe **TSA-BPa-Bt** (IC_50_ 313 nM)^44^ or **HPI-Bpa-Bt** (IC_50_ 107 nM)^25^ was added in presence and absence of 10 μM inhibitor. After incubation for 1.5 h at 37 °C, enzyme reaction mixtures were irradiated at 350 nm for 30 min on ice (Rayonet Photochemical Reactor). Biotinylated photoprobe-bound γ-secretase was pulled down using streptavidin beads (Sigma-Aldrich), washed six times with standard assay buffer, eluted with loading buffer for SDS-PAGE, and heated to 70 °C for 10 min. Presenilin N-terminal fragment (NTF) was detected by western blot using anti-presenilin-1 monoclonal antibody (Millipore).

### Cross-competition kinetic experiments

For cross-competition studies between two inhibitors, enzyme reactions were run as activity assays, where varying concentration of two inhibitors were titrated against each-other, keeping the concentrations of C100 (0.50 μM) constant.^31, 43^ Reciprocal plots were generated, and data were fit using GraphPad Prism 9.0 into the following equation:

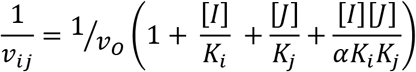

where, *v*_*ij*_ and *v*_0_ are the initial rate in the presence and absence, respectively, of inhibitors; *K*_*i*_ and *K*_*j*_ are the dissociation constants for inhibitors *I* and *J*, respectively; and α is the constant defining the interaction between the two inhibitors *I* and *J*.

## Supporting information

Supporting Information Available

## ASSOCIATED CONTENT

### Supporting Information Available

Detailed schematic method for the solid-phase synthesis, HRMS and LC-MS traces of final peptidomimetic conjugates, Michaelis-Menten plots of top inhibitors, and Molecular String Formulas (as a CSV file) are provided in supporting information. This material is available free of charge in the Internet at http://pubs.acs.org.

## Conflict of interest

The authors declare no conflicts of interest.

## ACKNOWLEDGMENTS

This work was supported by NIH grants GM122894 and AG66986 to M.S.W. We thank Dr. Benjamin Neuenswander (University of Kansas Specialized Chemistry Center) for LC-MS purity analysis.

## Abbreviations

Aβ: amyloid β-protein
AD: Alzheimer’s disease
Aib: α-aminoisobutyric acid
APP: amyloid precursor protein
CTF: C-terminal fragment
cryo-EM: cryo-electron microscopy
HPIs: helical peptide inhibitors
HPI-TSA: helical peptide inhibitor/transition-state analogue inhibitor conjugate
NTF: N-terminal fragment
PSEN1: presenilin-1
TSA: transition-state analogues inhibitor

## Table of Contents Graphic

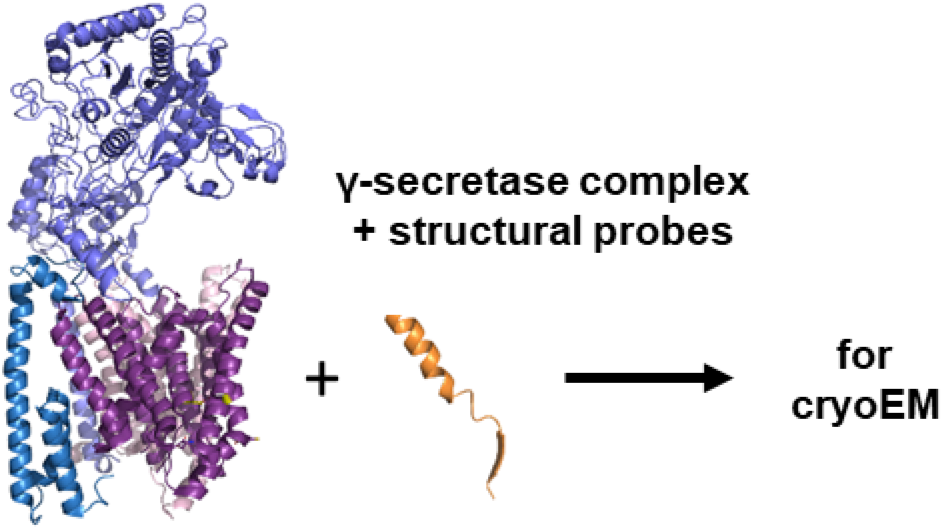

## Notes

### Competing Interest Statement

The authors have declared no competing interest.

