## Supporting Information Available for "Design of Transmembrane Mimetic Structural Probes to Trap Different Stages of γ-Secretase-Substrate Interaction"

### Table of Contents

Scheme S1: Solid-phase synthesis of *N*-acetylated, C-amide peptidomimetics S2
Scheme S2: Solid-phase synthesis of *N*-Boc, C-amide peptidomimetics S3
Scheme S3: Solid-phase synthesis of *N*-Boc, C-amide D-peptidomimetics S4
Figure S1: Michaelis-Menten plot of top inhibitors S5
Figure S2 to S20: HRMS traces of synthesized peptides S6
Figure S21 to S39: LC-MS traces of synthesized peptides S25

**Scheme S1.** Solid-phase synthesis of *N*-acetylated, C-amide L-peptidomimetics using Rink amide resin (for peptides **20, 21, 30, 31**).[a]

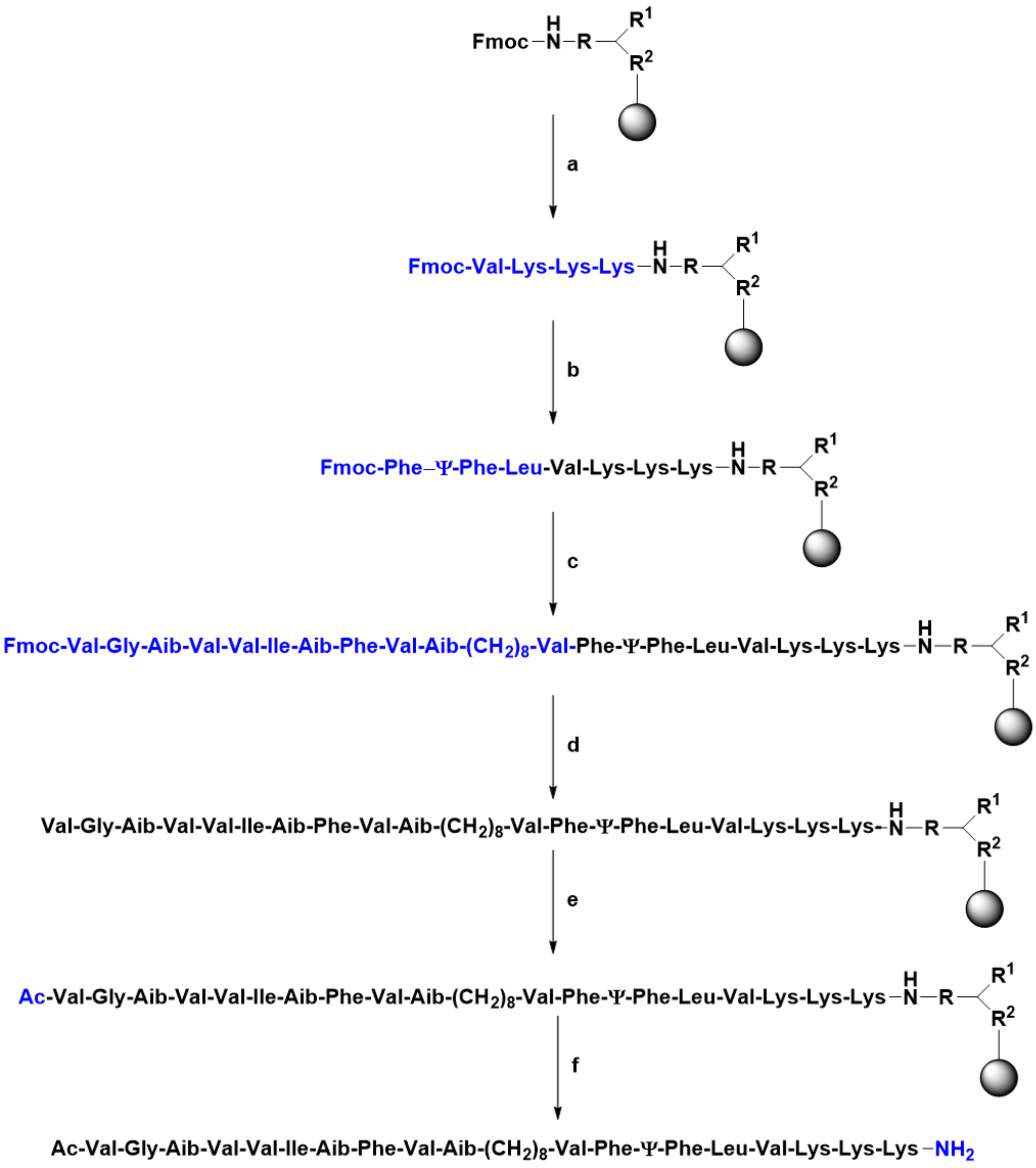


[a]Reagents and conditions: (a) Iteratively: i. 20% piperidine in DMF; ii. 0.2 M Fmoc-protected amino acid (3 x ε-*N*-Boc-Lys, then Val), 0.2 M DIC (*N,N′*-diisopropylcarbodiimide), and 0.2 M OXYMA (ethyl cyano(hydroxyimino)acetate) in DMF, 70 $^oC$, 8 min, double coupling; (b) i. 20% piperidine in DMF; (ii) 0.2 M **11**, 0.2 M DIC, and 0.2 M OXYMA in DMF, 70 $^oC$, 8 min, double coupling; (c) Iteratively: i. 20% piperidine in DMF; ii. 0.2 M Fmoc-protected L-amino acids (Val, Fmoc-NH($CH_2$)$_8$$CO_2$H for peptide **20** and **21** or 3 x Gly for peptide **30** or 4 x Gly for peptide **31**, Aib, Val, Phe, Aib, Ile, Val, Val, Aib, Gly, Val), 0.2 M DIC, and 0.2 M OXYMA in DMF, 70 $^oC$, 8 min, double coupling; (d) 20% piperidine in DMF; (e) $Ac_2O$, 7% DIPEA in DMF, 60 min (f) TFA:TIPS (triisopropylsilane): $H_2O$: DoDt (2,2′-(ethylenedioxy)diethanethiol):: 92.5:2.5:2.5:2.5, r.t., 2 h. (Note: for the synthesis of peptide **21** in step a, the amino acid residue coupled was only valine).

**Scheme S2.** Solid-phase synthesis of *N*-Boc, C-amide L-peptidomimetics using Rink amide resin (for peptide **22-29**).[a]

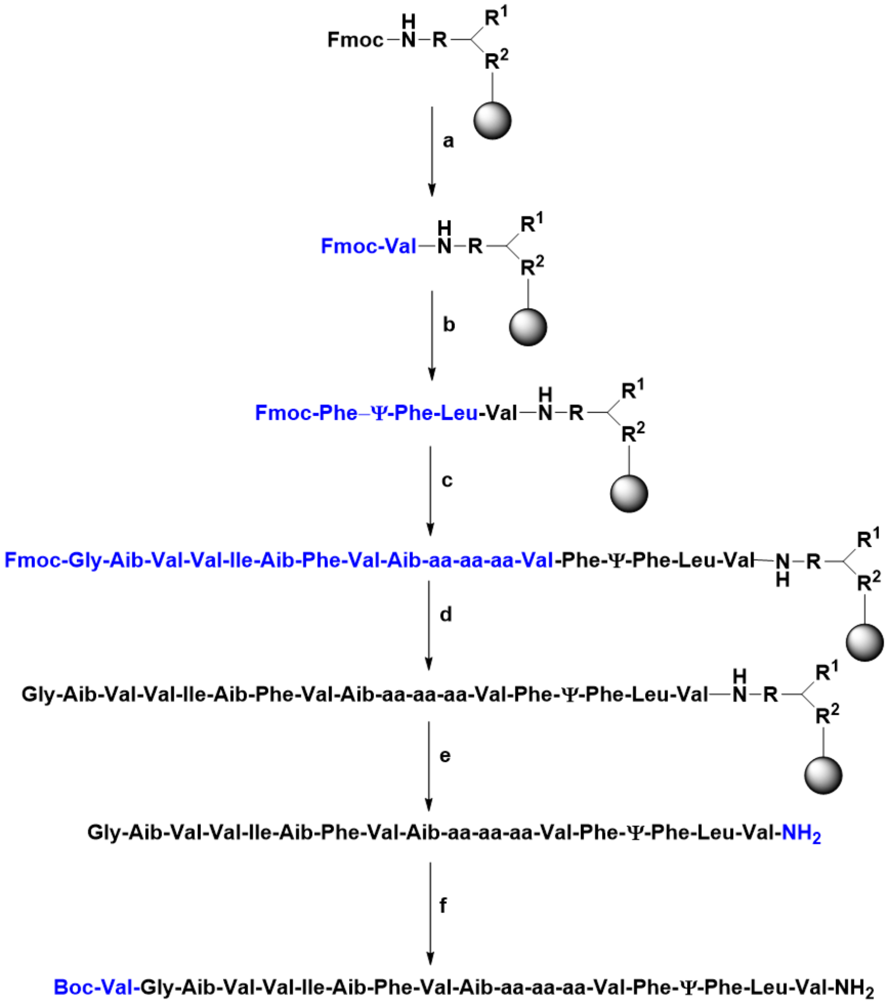


[a]Reagents and conditions: (a) (i) 20% piperidine in DMF; (ii) 0.2 M Fmoc-valine, 0.2 M DIC, and 0.2 M OXYMA in DMF, 70 °C, 8 min, double coupling; (b) (i) 20% piperidine in DMF; (ii) 0.2 M **11**, 0.2 M DIC, and 0.2 M OXYMA in DMF, 70 °C, 8 min, double coupling; (c) Iteratively: i. 20% piperidine in DMF; ii. 0.2 M Fmoc-amino acids (Val, 3 amino acids replacing 10-atom linker region, Aib, Val, Phe, Aib, Ile, Val, Val, Aib, Gly), 0.2 M DIC, and 0.2 M OXYMA in DMF, 70 °C, 8 min, double coupling (d) 20% piperidine in DMF (e) TFA:TIPS:$H_2O$:DoDt :: 92.5:2.5:2.5:2.5, rt, 2 h; (f) 1.0 eq. Boc-valine, 0.9 eq. HCTU, 2.0 eq. DIPEA, 3 mL DMF, rt, 24 h, yield 45-50%. (Note: for the synthesis of peptide **22, 23, 24, 25, 26, 27, 28, 29** three amino acids replacing linker region were VIV, VIG, VGG, IVI, IVG, IGG, GGG, and GGI, respectively. All the peptides were cleaved from resin at their penultimate length, and the terminal Boc-Val-OH was attached in solution phase using ~0.1 mmol penultimate peptide precursor).

**Scheme S3**. Solid-phase synthesis of *N*-Boc, C-amide D-peptidomimetics using Rink amide resin (for peptide **33-39**).[a]

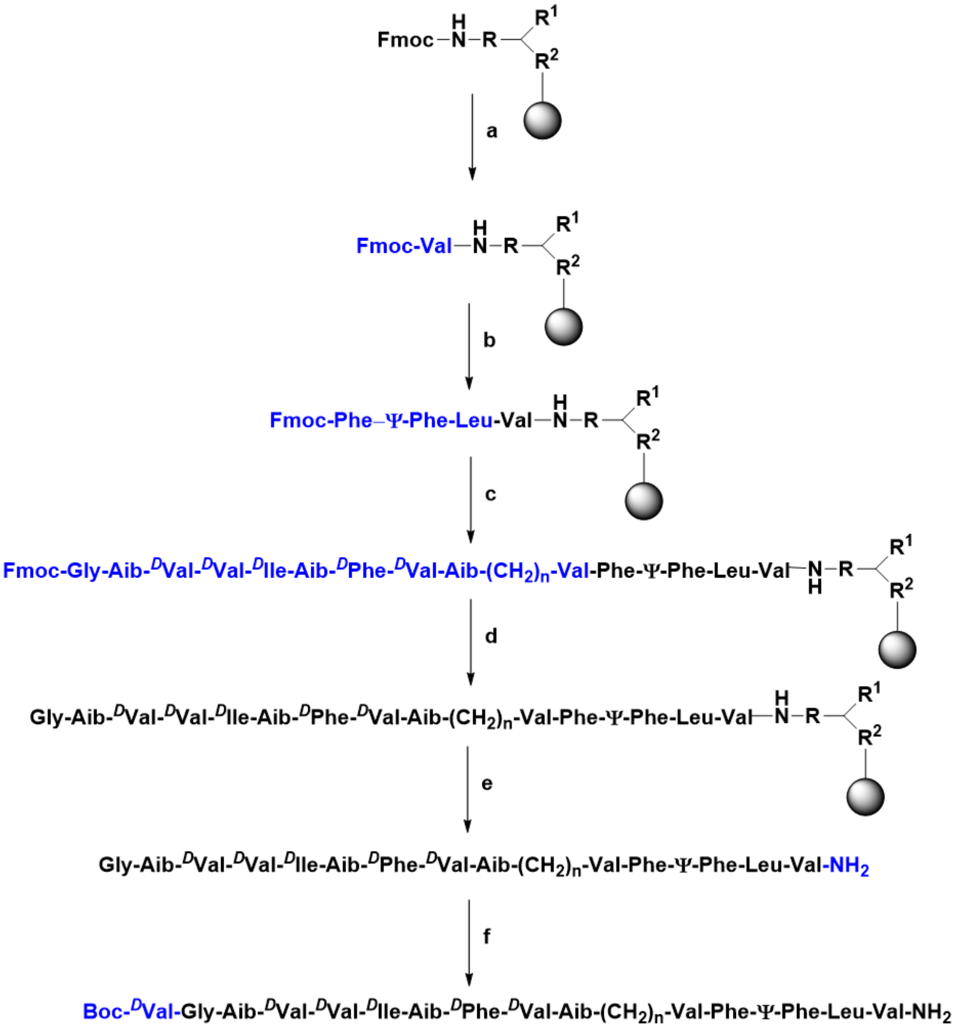


[a]Reagents and conditions: (a) (i) 20% piperidine in DMF; (ii) 0.2 M Fmoc-valine, 0.2 M DIC, and 0.2 M OXYMA in DMF, 70 °C, 8 min, double coupling; (b) Iteratively: (i) 20% piperidine in DMF; (ii) 0.2 M **11**, 0.2 M DIC, and 0.2 M OXYMA in DMF, 70 °C, 8 min, double coupling; (c) Iteratively: i. 20% piperidine in DMF; ii. 0.2 M Fmoc-amino acids (Val, Fmoc-NH$(CH_2)_nCO_2$H, Aib, $^D$Val, $^D$Phe, Aib, $^D$Ile, $^D$Val, $^D$Val, Aib, Gly), 0.2 M DIC, and 0.2 M OXYMA in DMF, 70 °C, 8 min, double coupling (d) 20% piperidine in DMF (e) TFA:TIPS:$H_2O$:DoDt :: 92.5:2.5:2.5:2.5, rt, 2h; (f) 1.0 eq. Boc-*D*-valine, 0.9 eq. HCTU, 2.0 eq. DIPEA, 3 mL DMF, rt, 24 h, yield 45-50%. (Note: All peptides were cleaved from the resin at their penultimate length, and the terminal Boc-$^D$Val-OH was attached in solution phase using ~0.1 mmol penultimate peptide precursor. Alkyl spacers n = 0, 2, 4, 8, 10, and 11 were used for HPI-TSAs **33**-**39**, respectively).

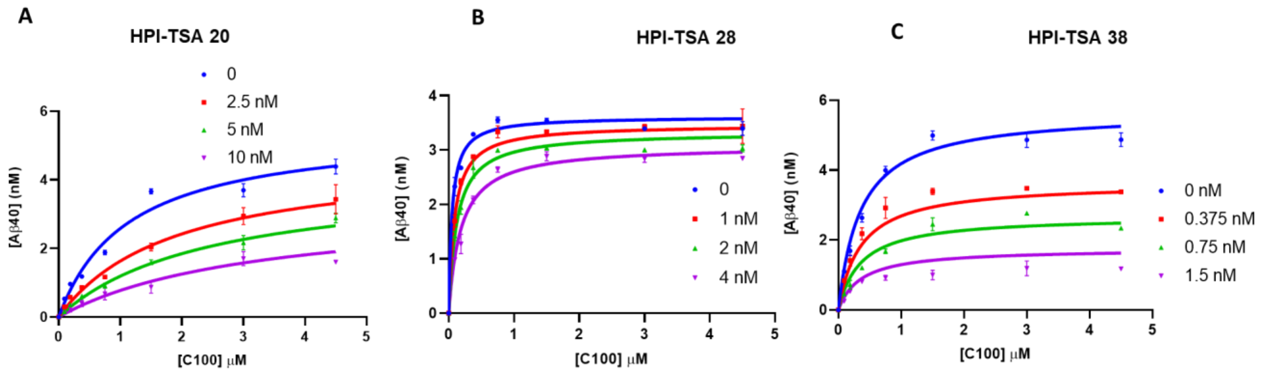


**Figure S1. Michaelis-Menten plot of top inhibitors of γ-secretase designed for each stage of substrate recognition. A**) Compound **20**, designed to trap the endoproteolysis transition state ($K_i$ = 2.64 ± 0.18 nM). **B**) Compound **28**, designed to capture helix unwinding ($K_i$ = 1.19 ± 0.10 nM). **C**) Compound **38**, designed to capture lateral gating ($K_i$ = 0.65 ± 0.02 nM).

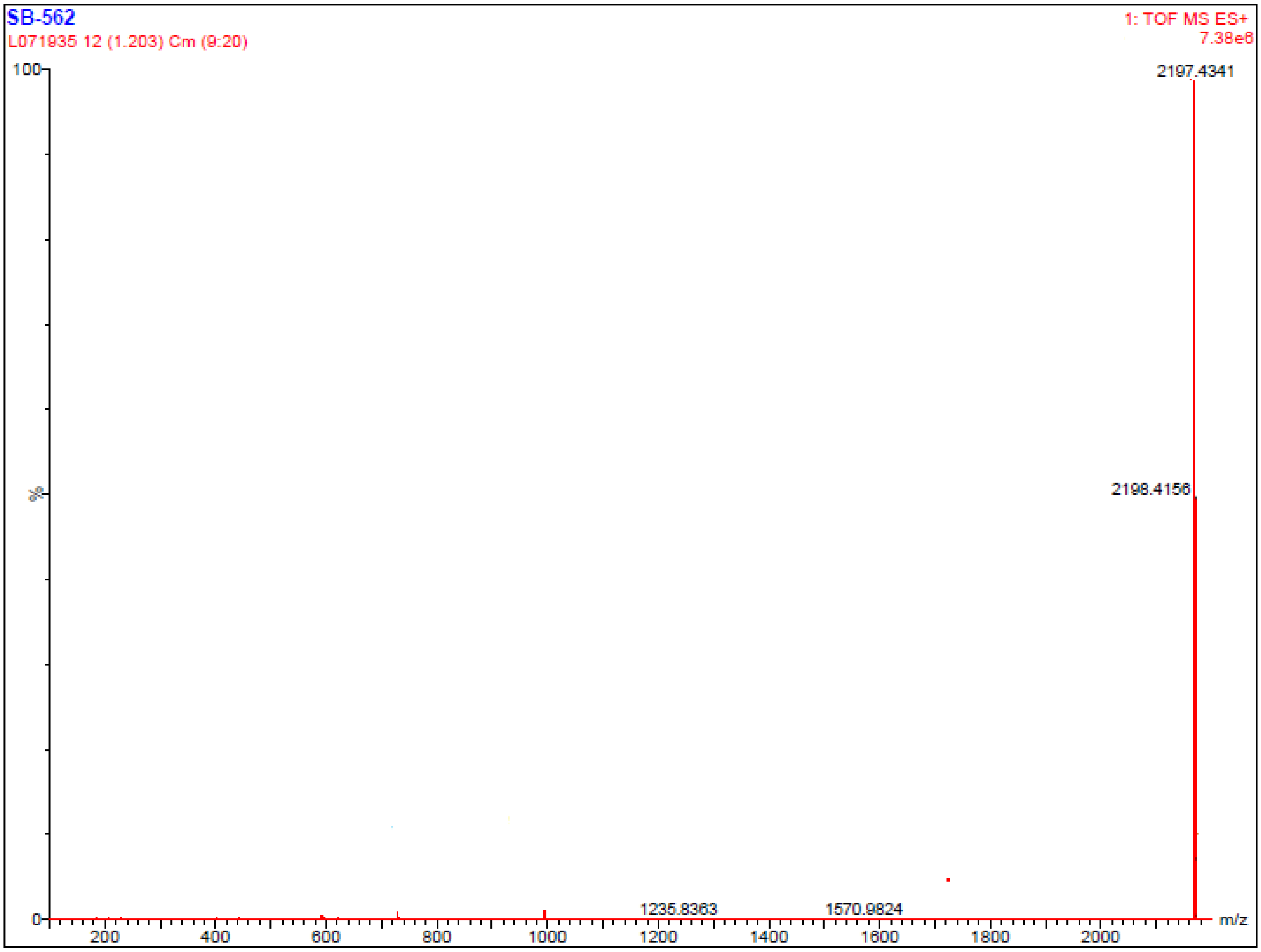


**Figure S2**: Section of HRMS (ESI): *m/z* [M + Na]$^+$ spectra for the synthesized peptide **20**. calcd for $C_{112}H_{187}N_{23}O_{20}Na$: 2197.4221; found: 2197.4341.

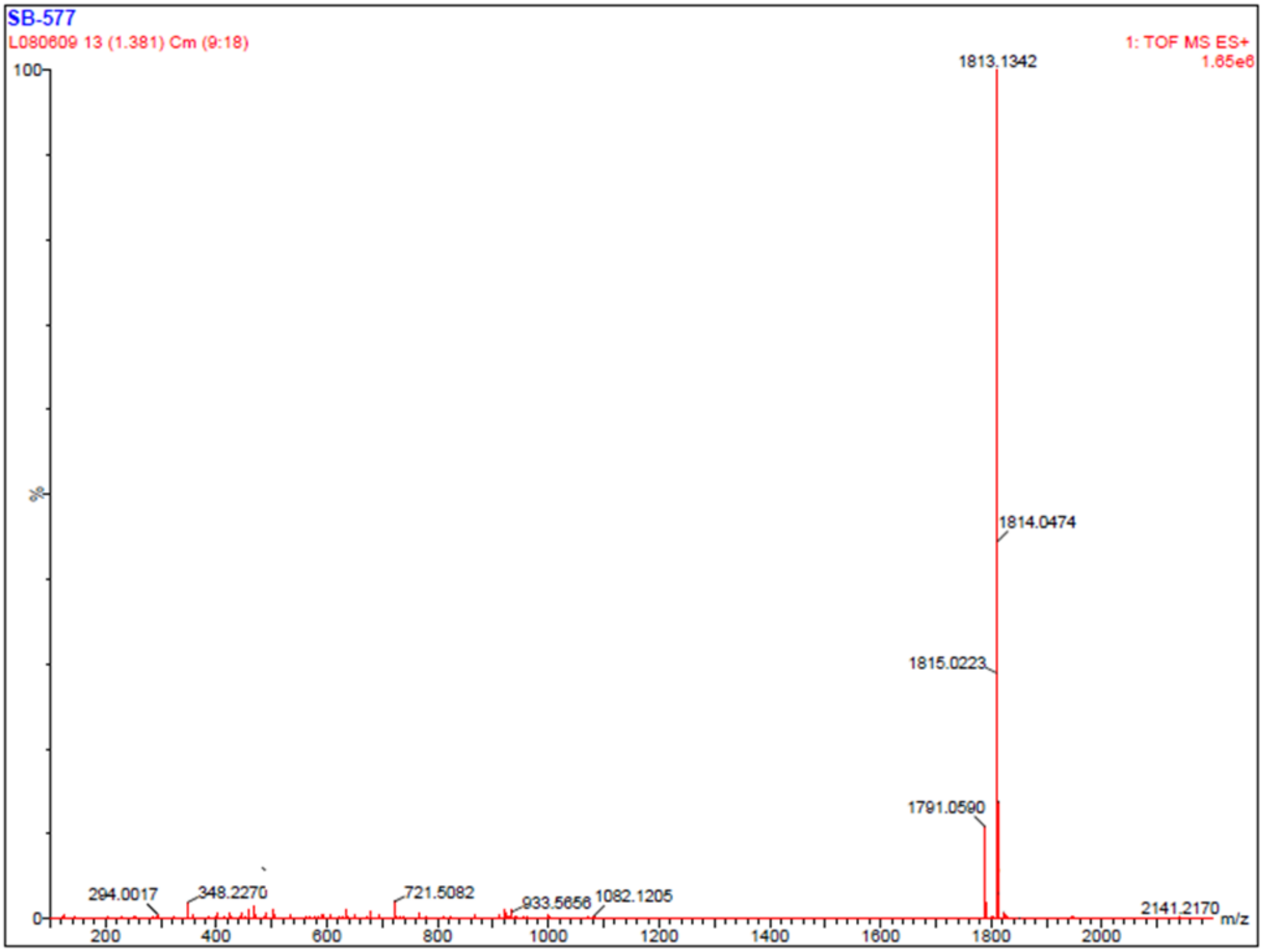


**Figure S3**: Section of HRMS (ESI): *m*/*z* $[M + Na]^+$ spectra for the synthesized peptide **21**. calcd for $C_{112}H_{187}N_{23}O_{20}Na$: 1813.1372; found: 1813.1342.

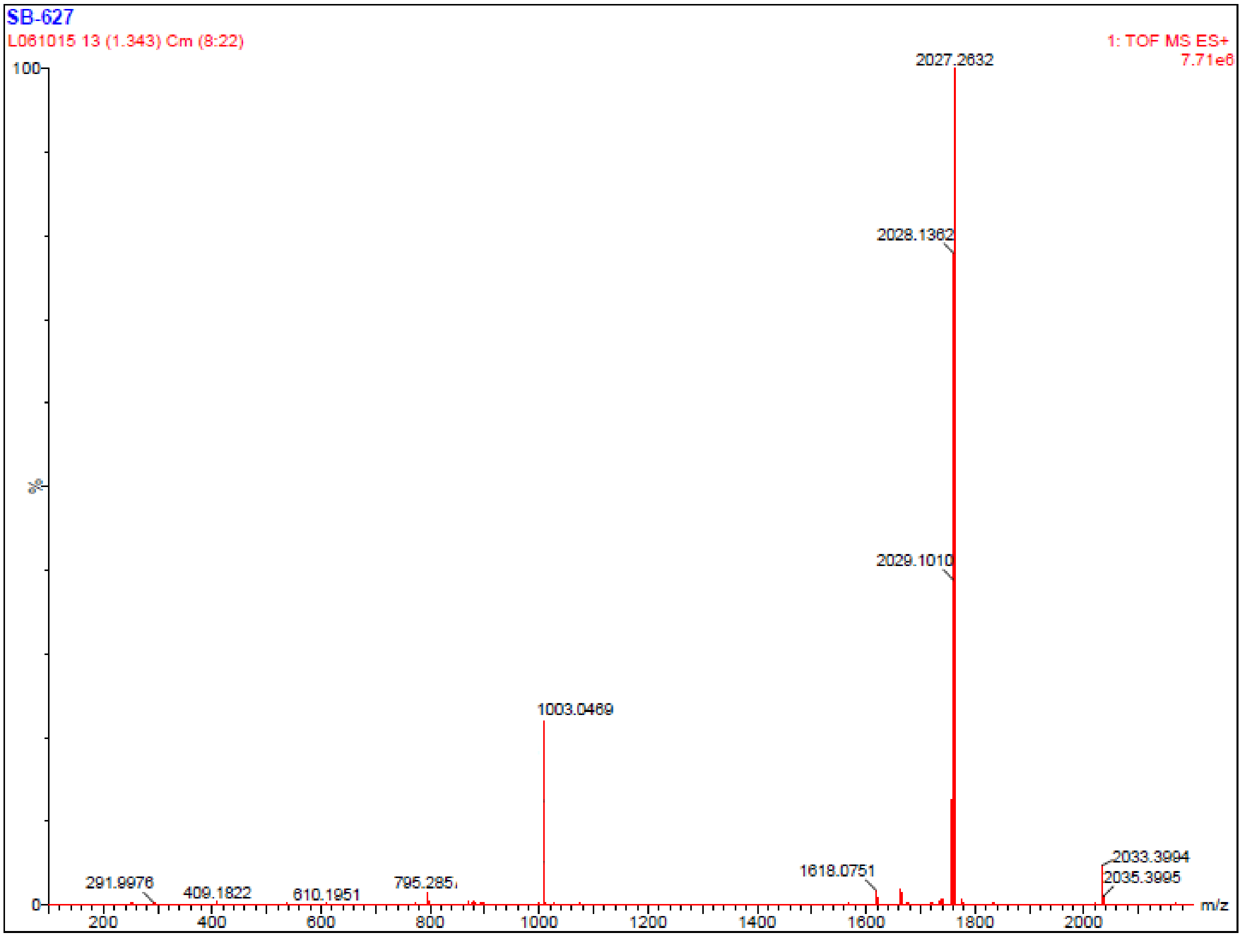


**Figure S4**: Section of HRMS (ESI): *m*/*z* [M + Na]$^+$ spectra for the synthesized peptide **22**. calcd for $C_{104}H_{169}N_{19}O_{20}Na$: 2027.2689; found: 2027.2632.

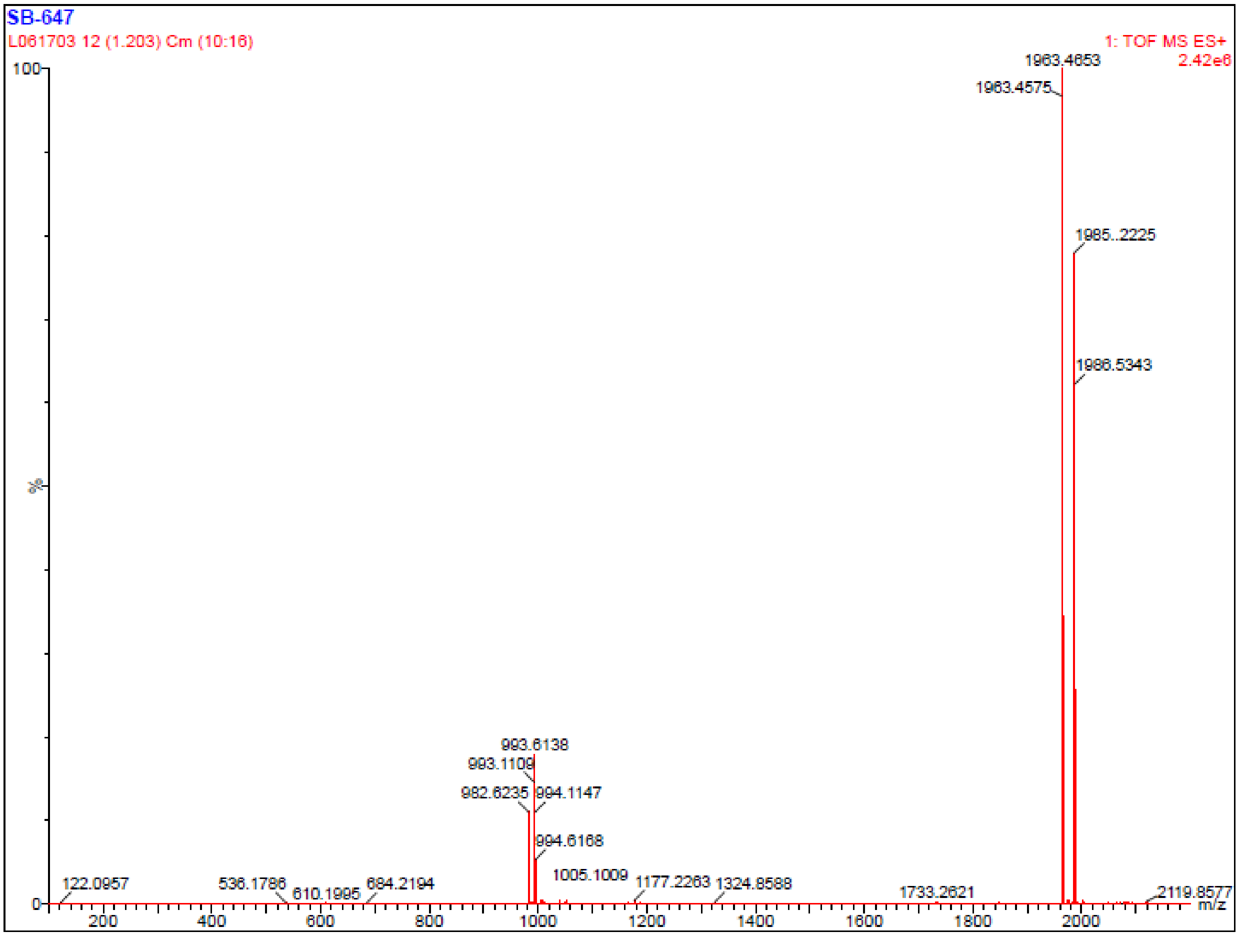


**Figure S5**: Section of HRMS (ESI): *m*/*z* $[M + Na]^+$ spectra for the synthesized peptide **23**. calcd for $C_{101}H_{163}N_{19}O_{20}Na$: 1985.2220; found: 1985.2225.

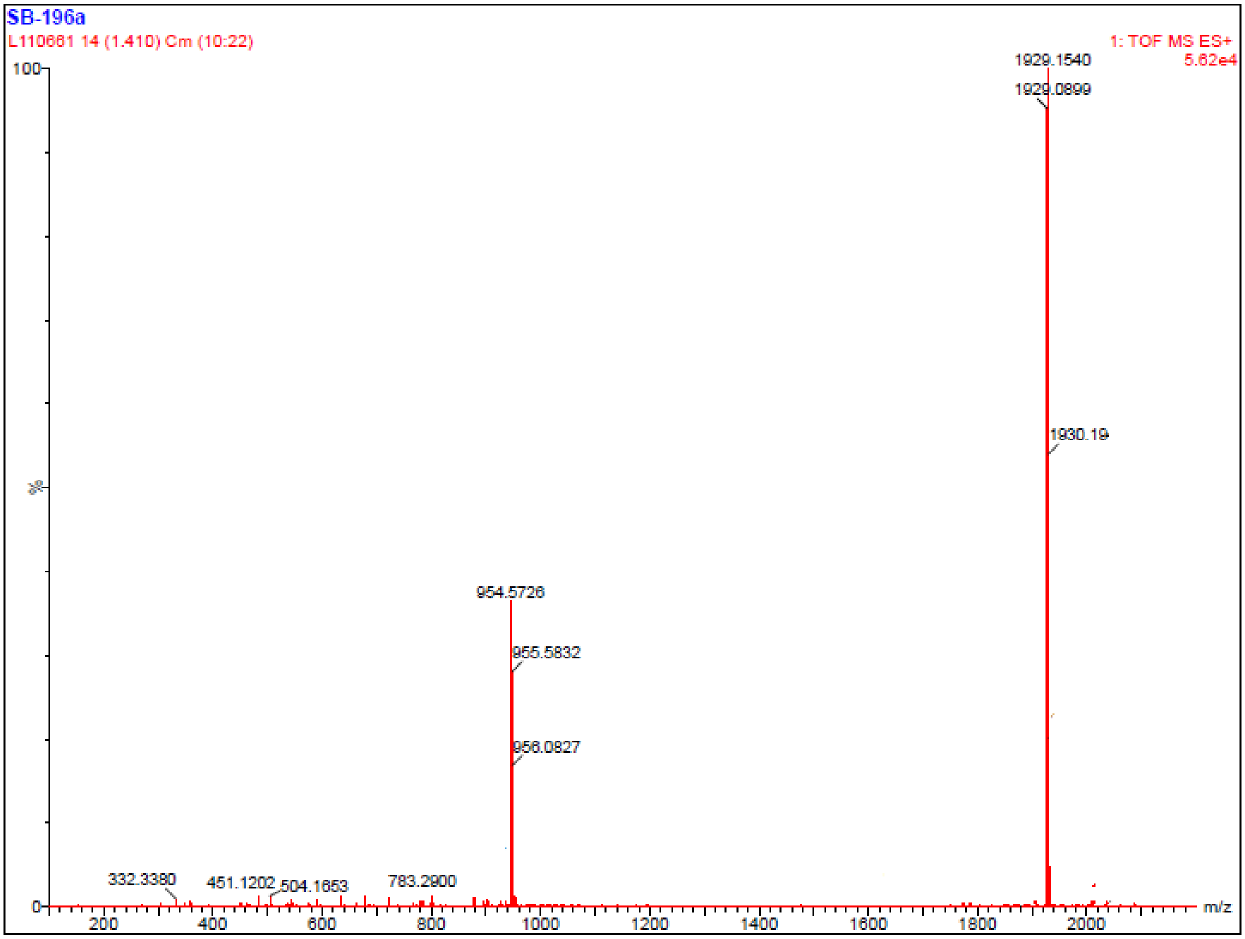


**Figure S6**: Section of HRMS (ESI): *m*/*z* $[M + Na]^+$ spectra for the synthesized peptide **24**. calcd for $C_{97}H_{155}N_{19}O_{20}Na$: 1929.1594; found: 1929.1540.

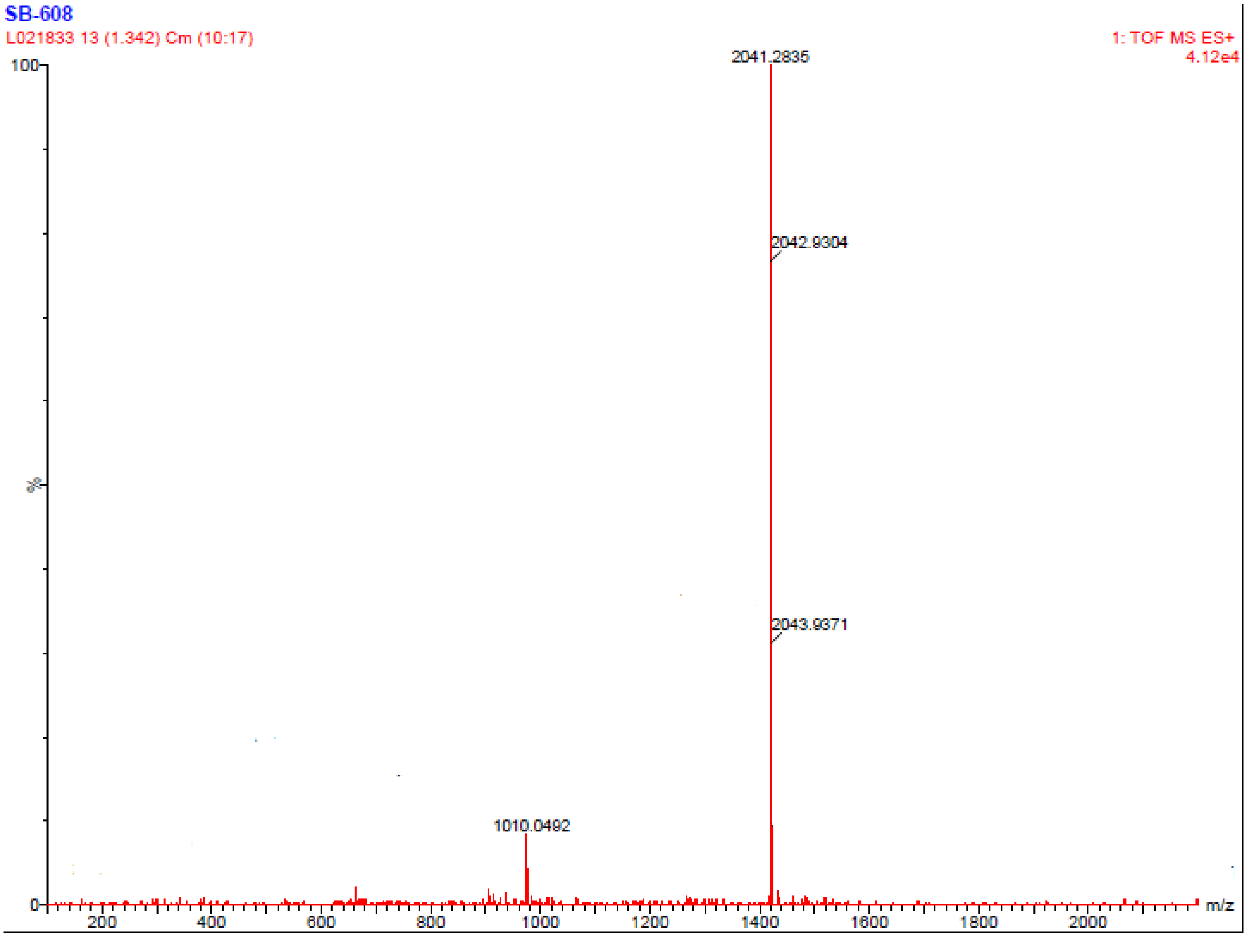


**Figure S7**: Section of HRMS (ESI): *m/z* $[M + Na]^+$ spectra for the synthesized peptide **25**. calcd for $C_{105}H_{171}N_{19}O_{20}Na$: 2041.2846; found: 2041.2835.

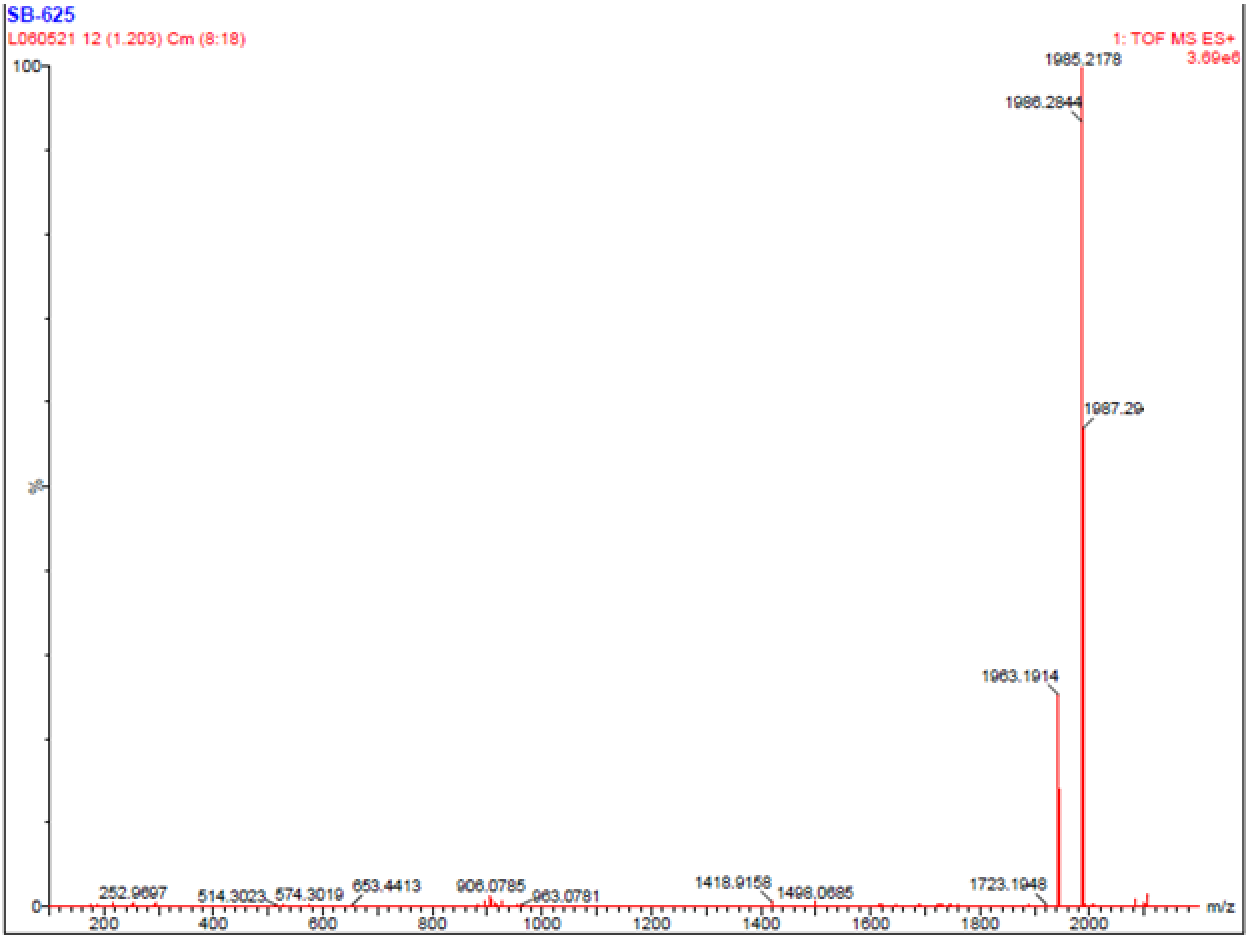


**Figure S8**: Section of HRMS (ESI): *m*/*z* $[M + Na]^+$ spectra for the synthesized peptide **26**. calcd for $C_{101}H_{163}N_{19}O_{20}Na$: 1985.2220; found: 1985.2178.

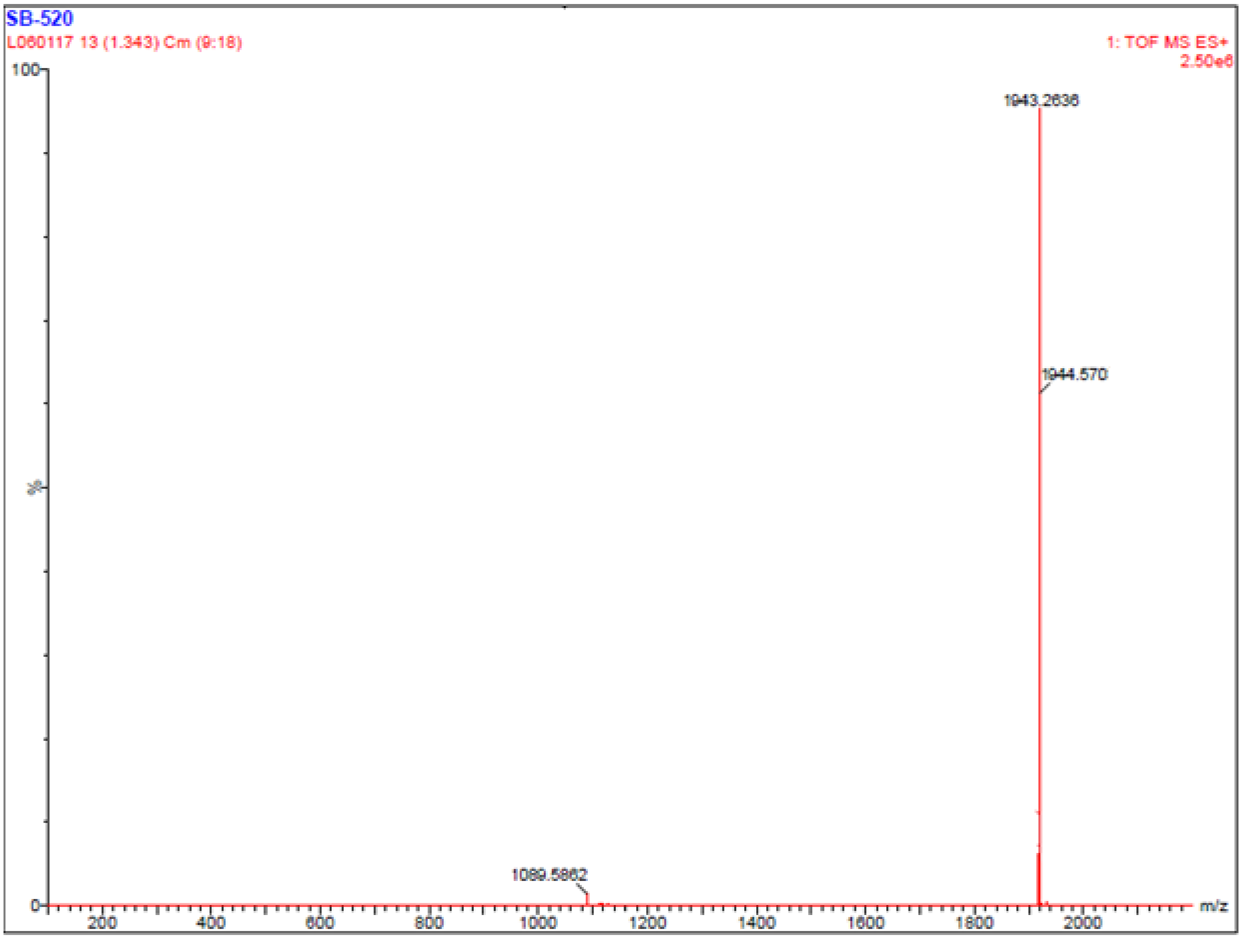


**Figure S9**: Section of HRMS (ESI): *m*/*z* $[M + Na]^+$ spectra for the synthesized peptide **27**. calcd for $C_{98}H_{157}N_{19}O_{20}Na$: 1943.1750; found: 1943.2187.

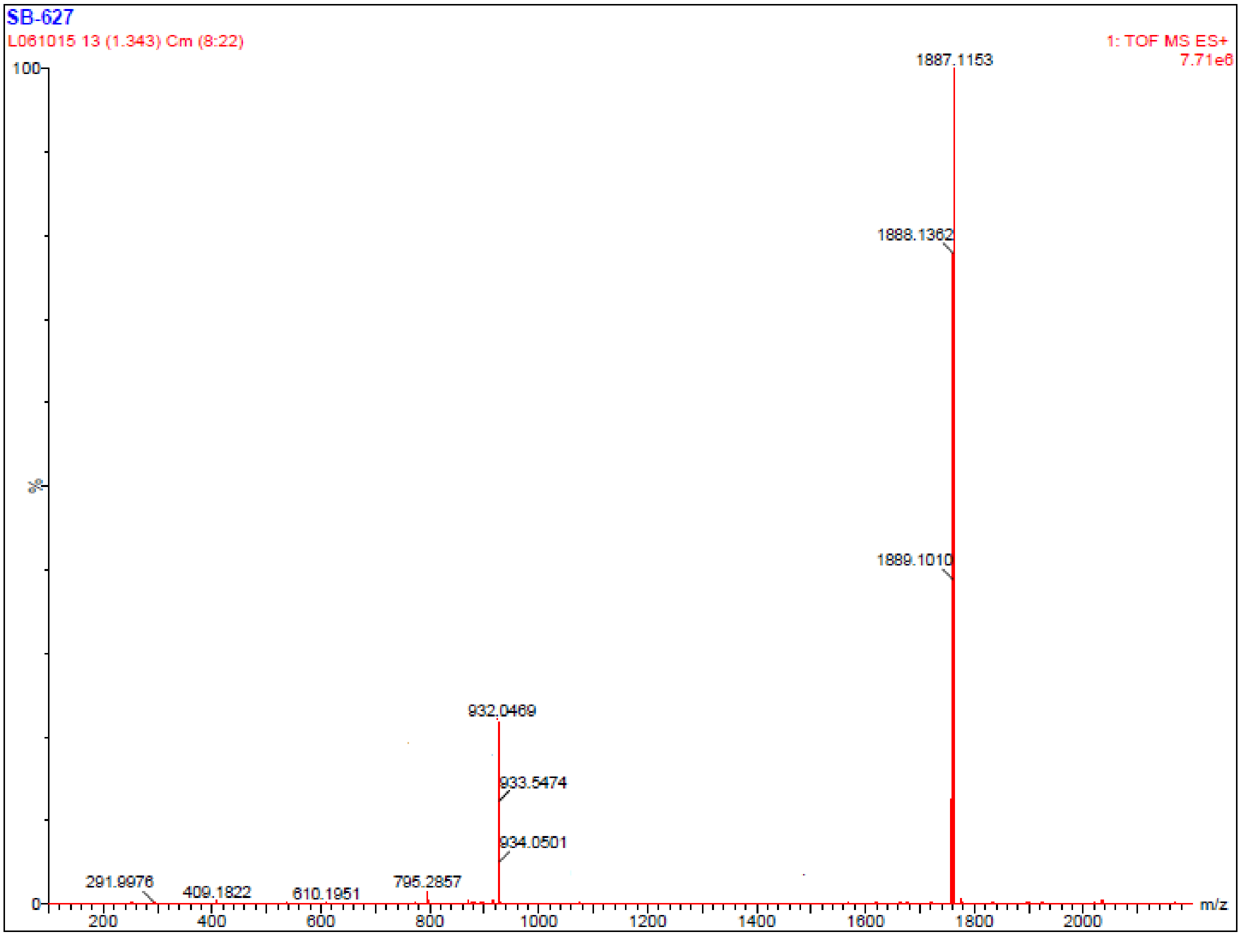


**Figure S10**: Section of HRMS (ESI): *m*/*z* $[M + Na]^+$ spectra for the synthesized peptide **28**. calcd for $C_{94}H_{149}N_{19}O_{20}Na$: 1887.1124; found: 1887.1153.

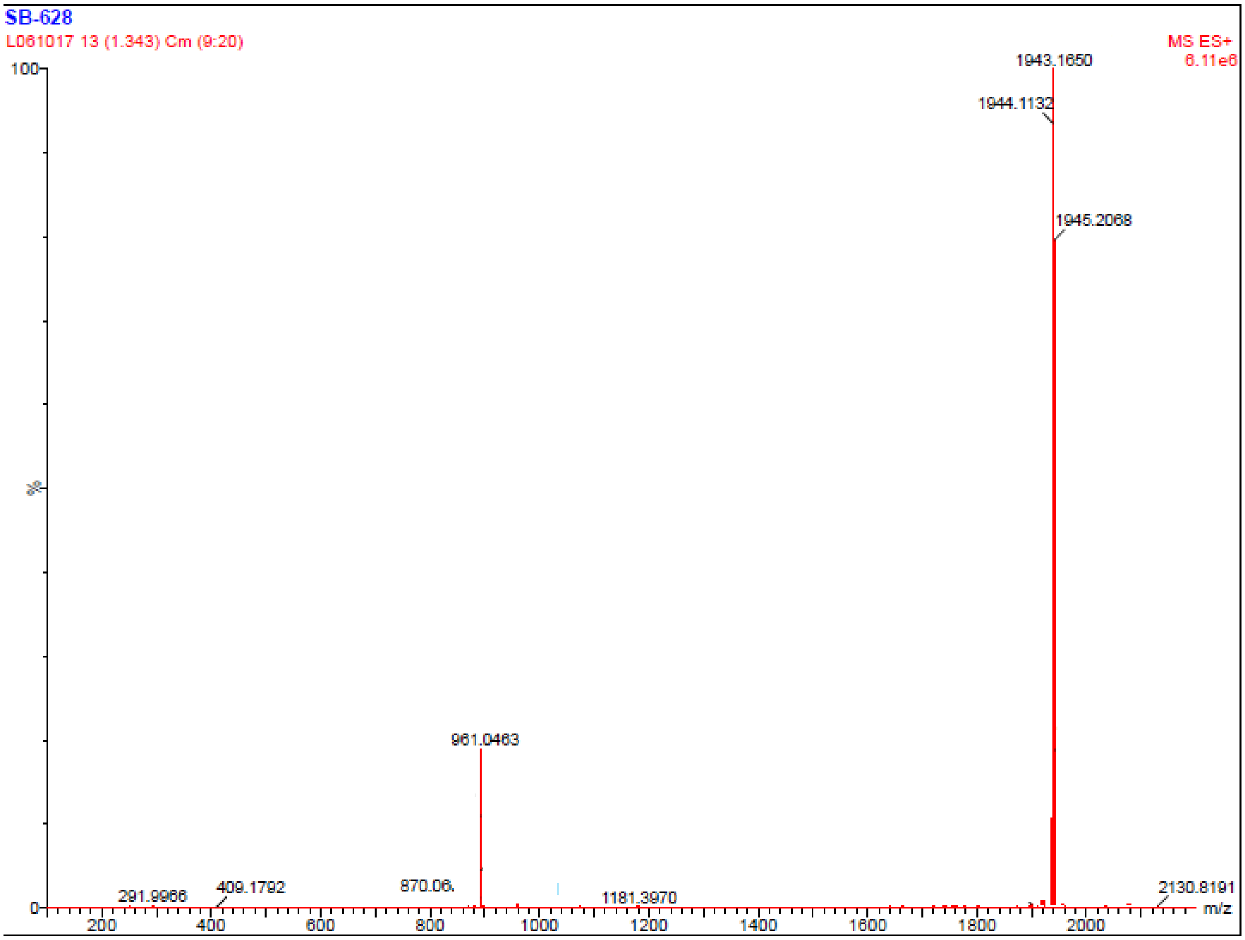


**Figure S11**: Section of HRMS (ESI): *m*/*z* [M + Na]$^+$ spectra for the synthesized peptide **29**. calcd for $C_{98}H_{157}N_{19}O_{20}Na$: 1943.1750; found: 1943.1650.

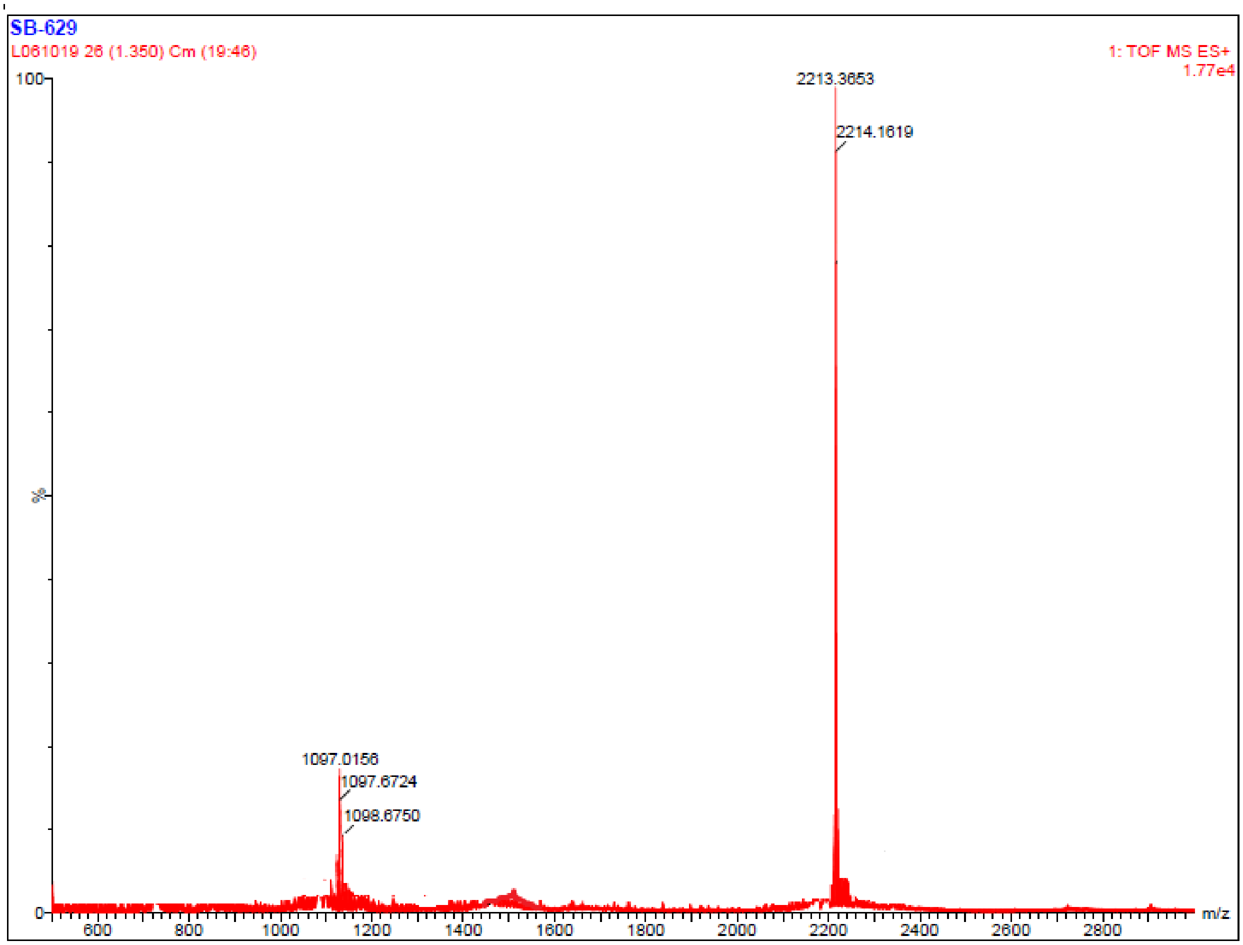


**Figure S12**: Section of HRMS (ESI): *m/z* $[M + Na]^+$ spectra for the synthesized peptide **30**. calcd for $C_{109}H_{179}N_{25}O_{22}Na$: 2213.3555; found: 2213.3653.

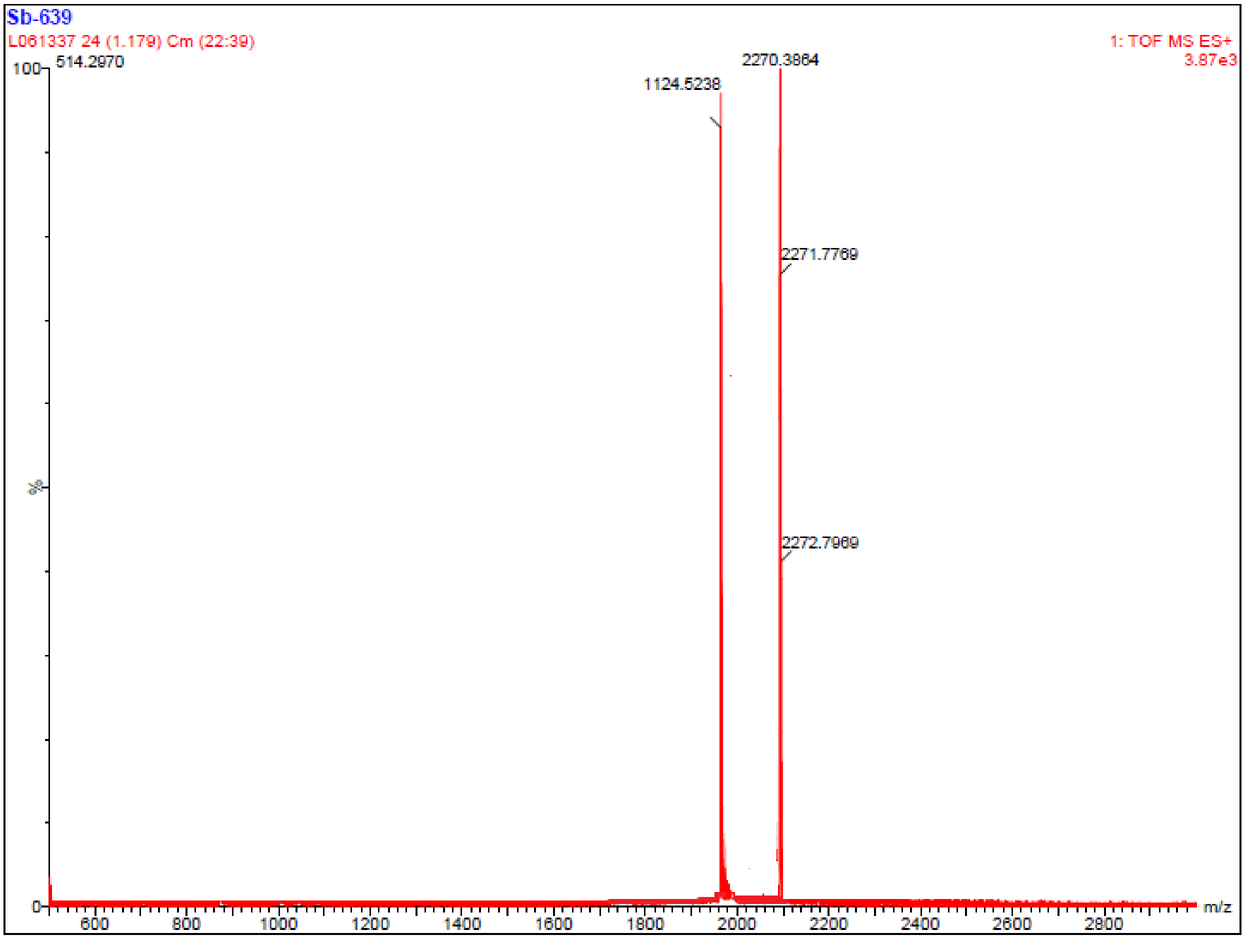


**Figure S13**: Section of HRMS (ESI): *m/z* [M + Na]$^+$ spectra for the synthesized peptide **31**. calcd for $C_{111}H_{182}N_{26}O_{23}Na$: 2270.3769; found: 2270.3864.

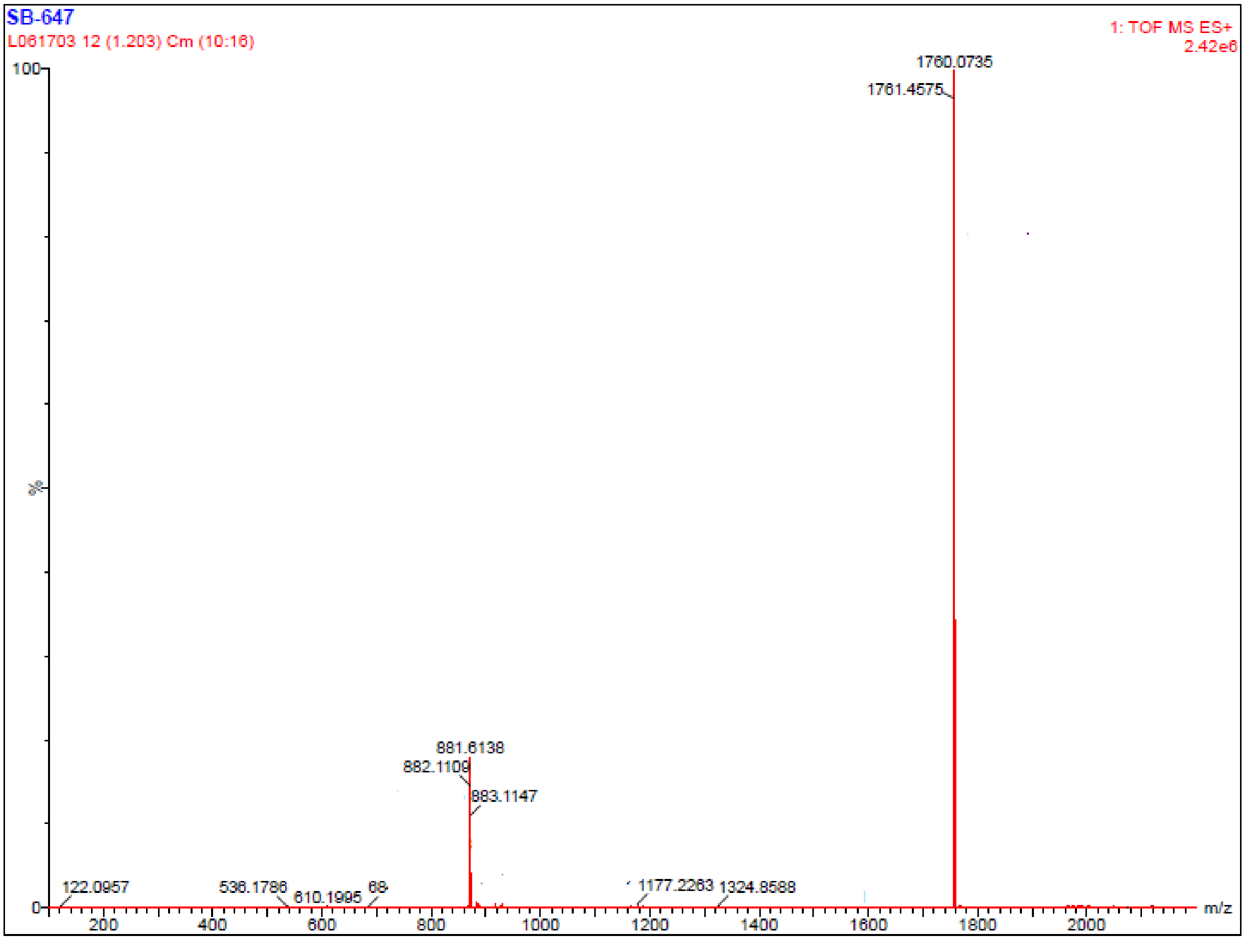


**Figure S14**: Section of HRMS (ESI): *m/z* $[M + Na]^+$ spectra for the synthesized peptide **33**. calcd for $C_{90}H_{144}N_{16}O_{18}Na$: 1760.0743; found: 1760.0735.

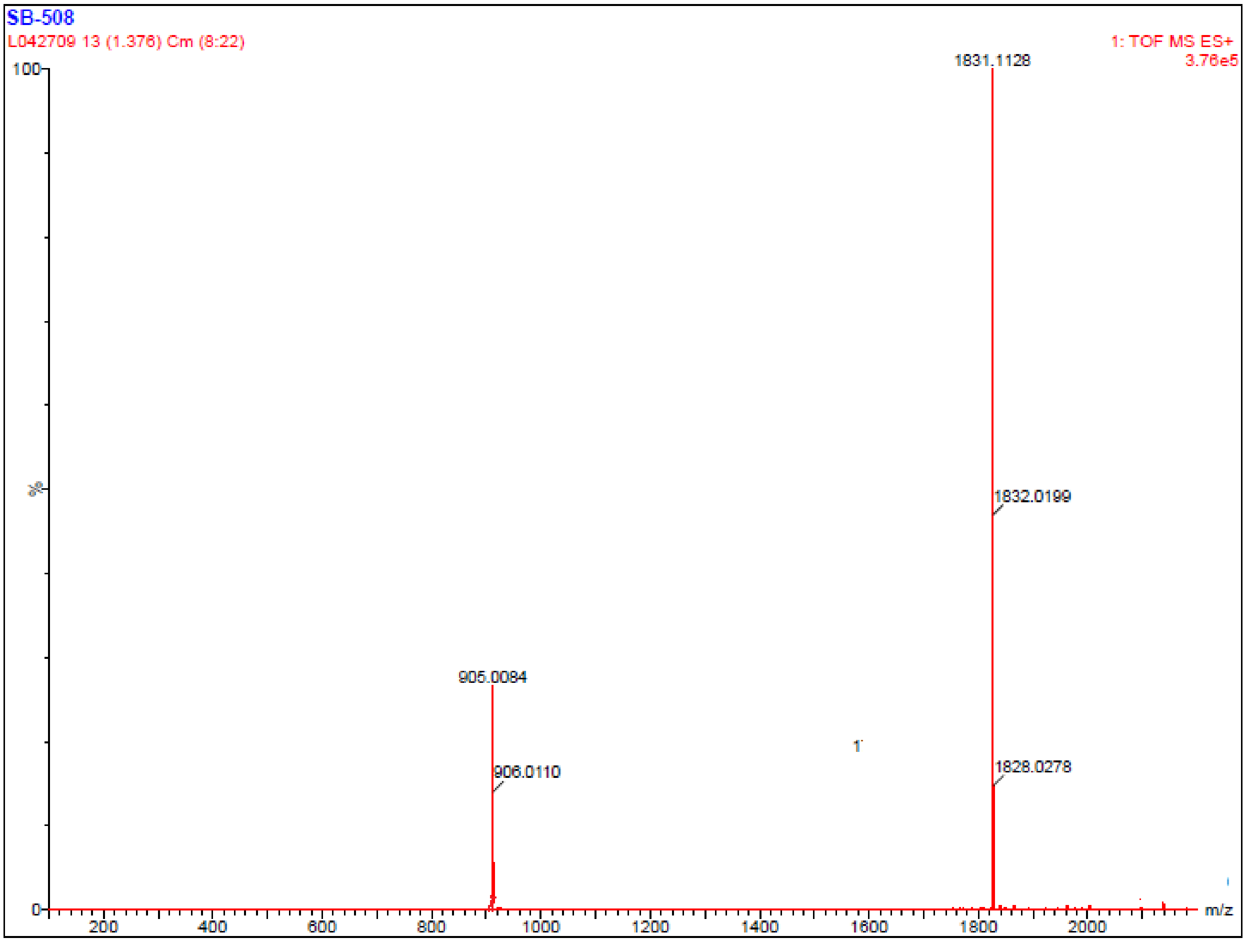


**Figure S15**: Section of HRMS (ESI): *m/z* $[M + Na]^+$ spectra for the synthesized peptide **34**. calcd for $C_{93}H_{149}N_{17}O_{19}Na$: 1831.1114; found: 1831.1128.

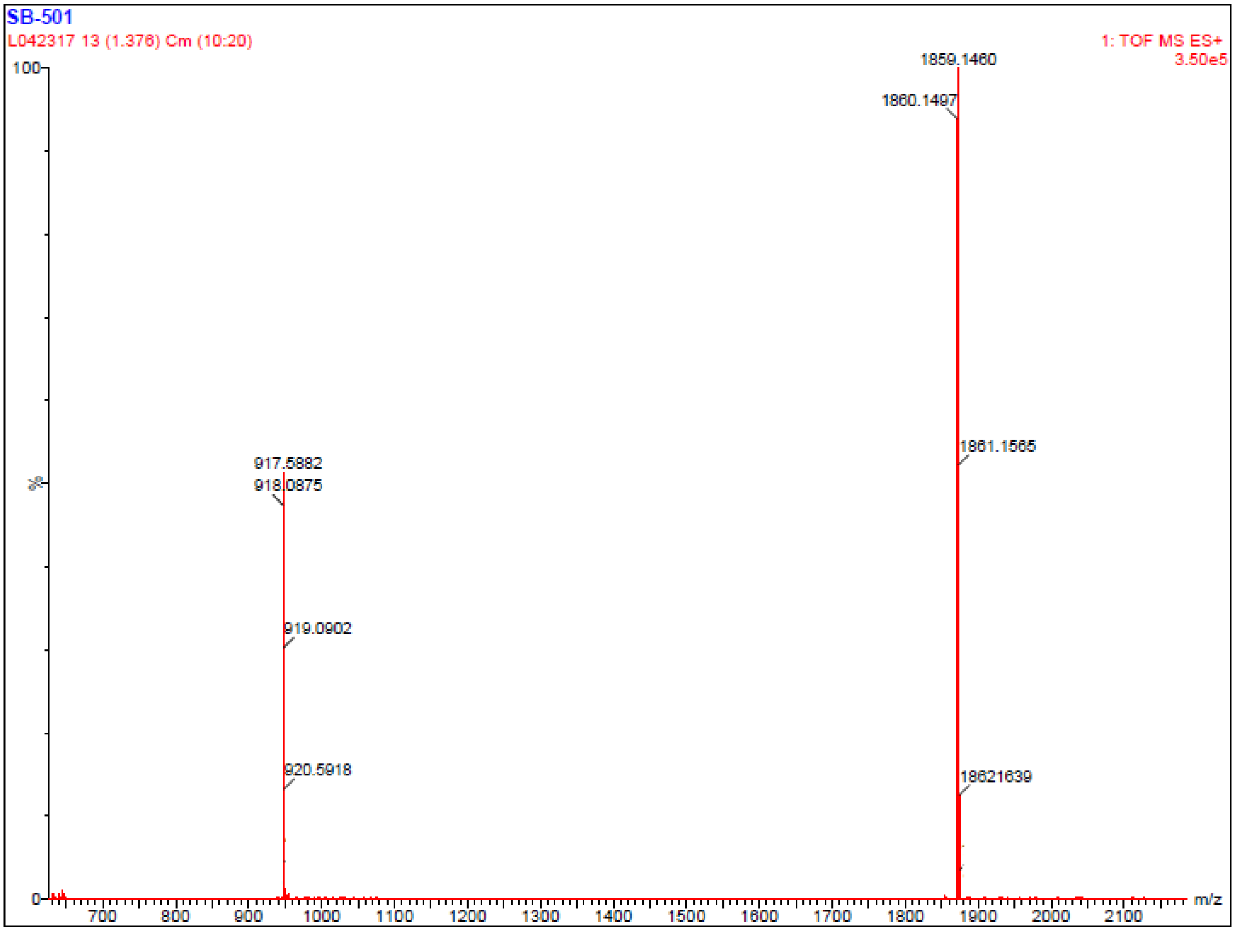


**Figure S16**: Section of HRMS (ESI): *m*/*z* [M + Na]$^+$ spectra for the synthesized peptide **35**. calcd for $C_{95}H_{153}N_{17}O_{19}Na$: 1859.1427; found: 1859.1460.

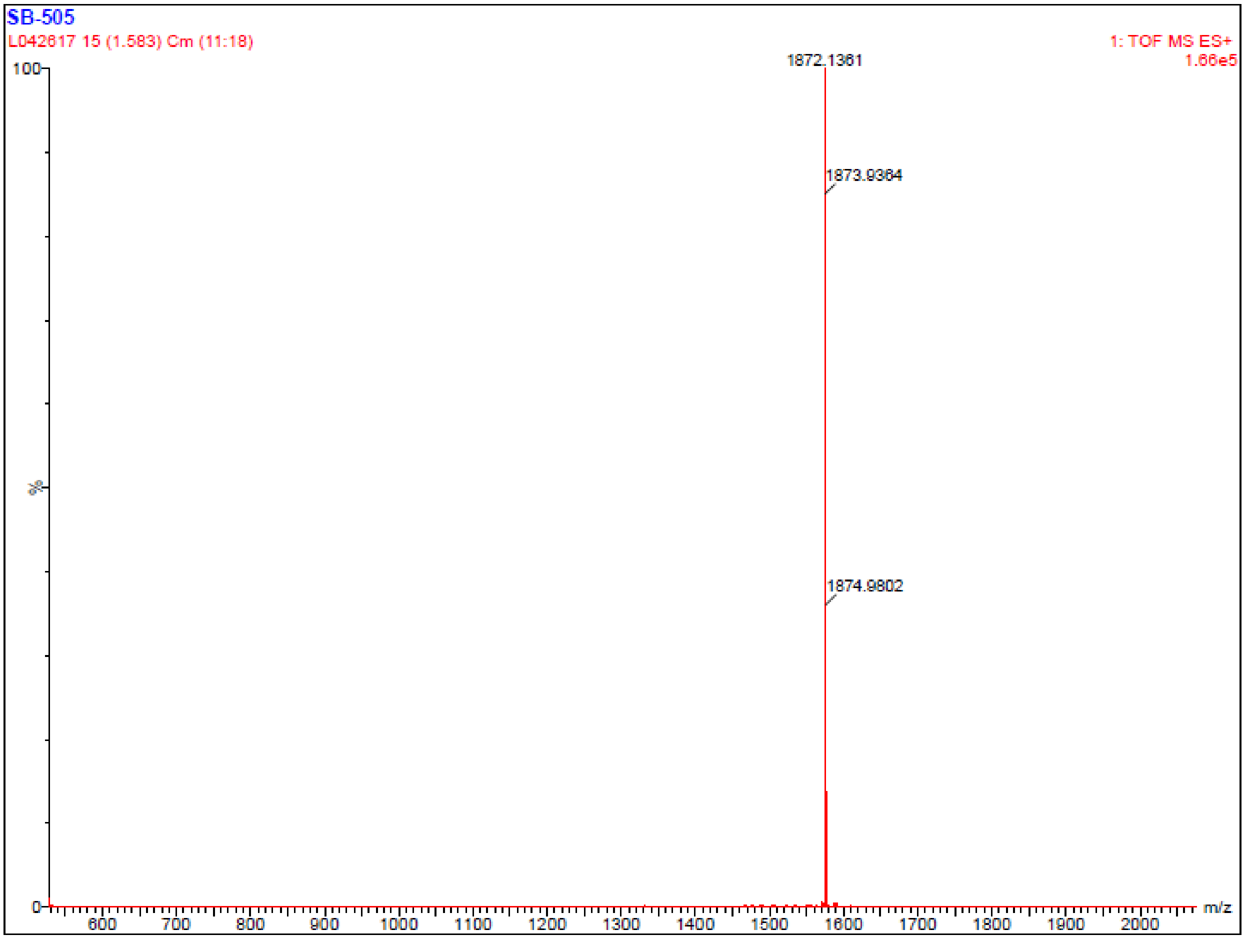


**Figure S17**: Section of HRMS (ESI): *m*/*z* [M + Na]$^+$ spectra for the synthesized peptide **36**. calcd for $C_{95}H_{153}N_{17}O_{19}Na$: 1872.1427; found: 1872.1361.

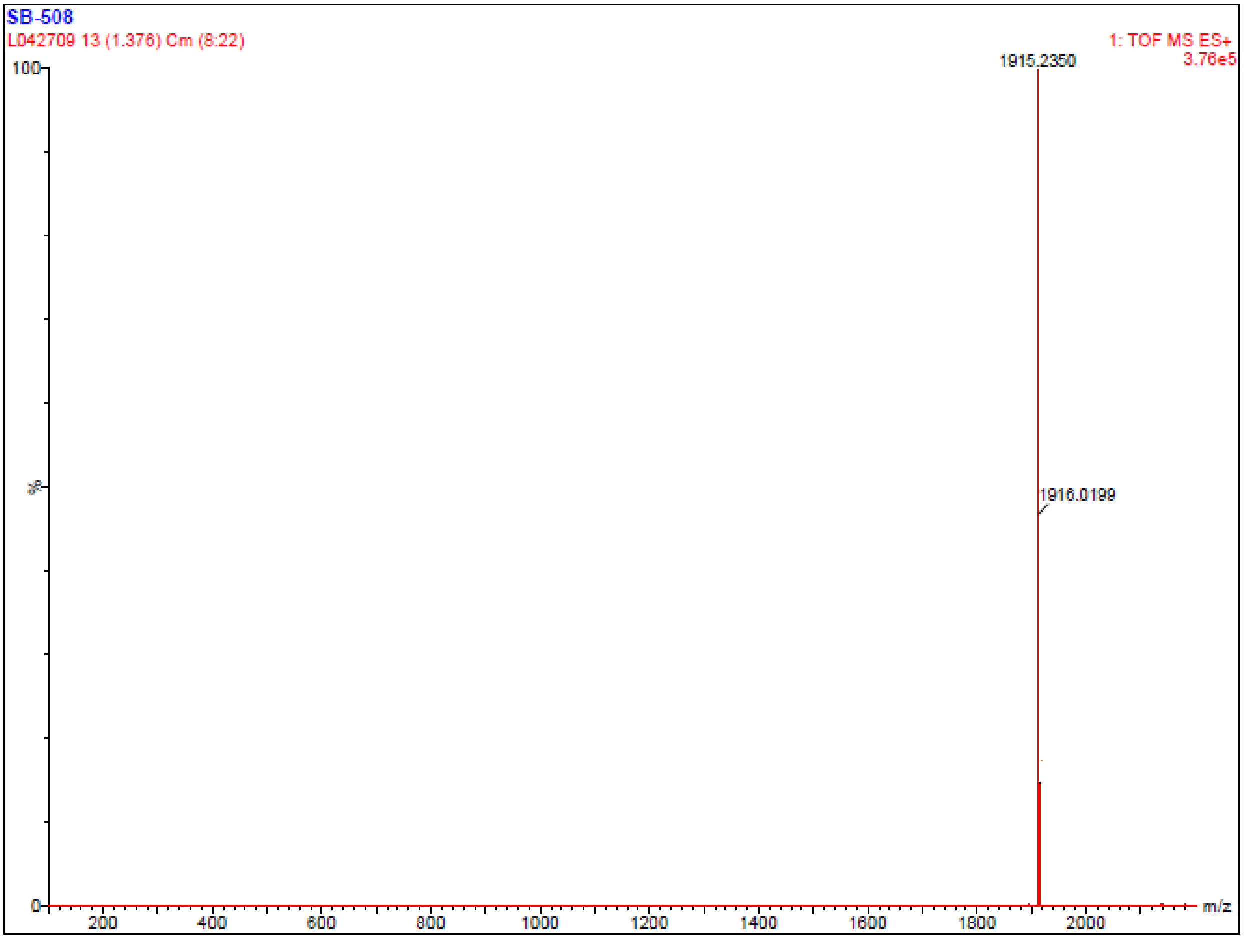


**Figure S18**: Section of HRMS (ESI): *m*/*z* $[M + Na]^+$ spectra for the synthesized peptide **37**. calcd for $C_{99}H_{161}N_{17}O_{19}Na$: 1915.2053; found: 1915.2350.

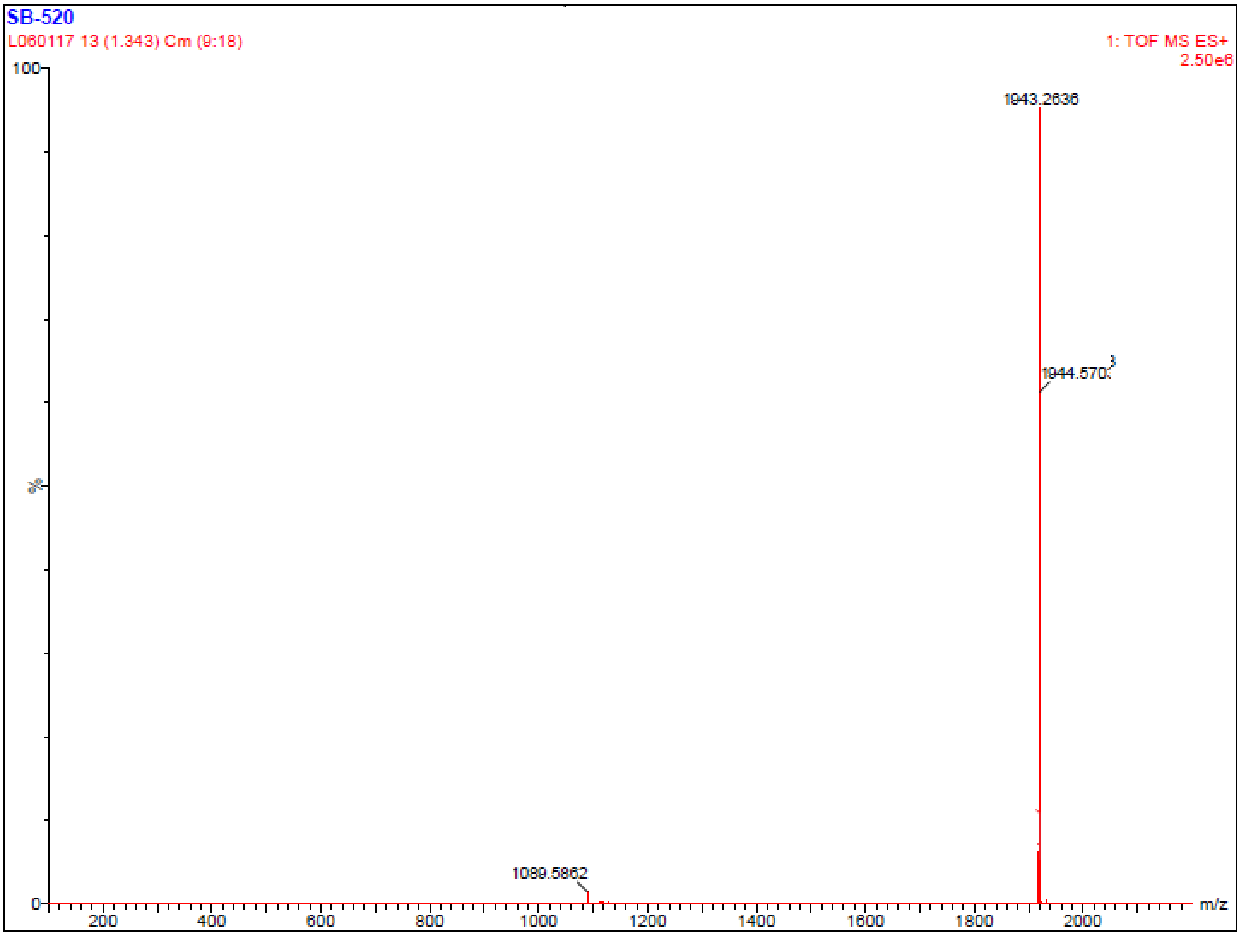


**Figure S19**: Section of HRMS (ESI): *m*/*z* $[M + Na]^+$ spectra for the synthesized peptide **38**. calcd for $C_{101}H_{165}N_{17}O_{19}Na$: 1943.2366; found: 1943.2636.

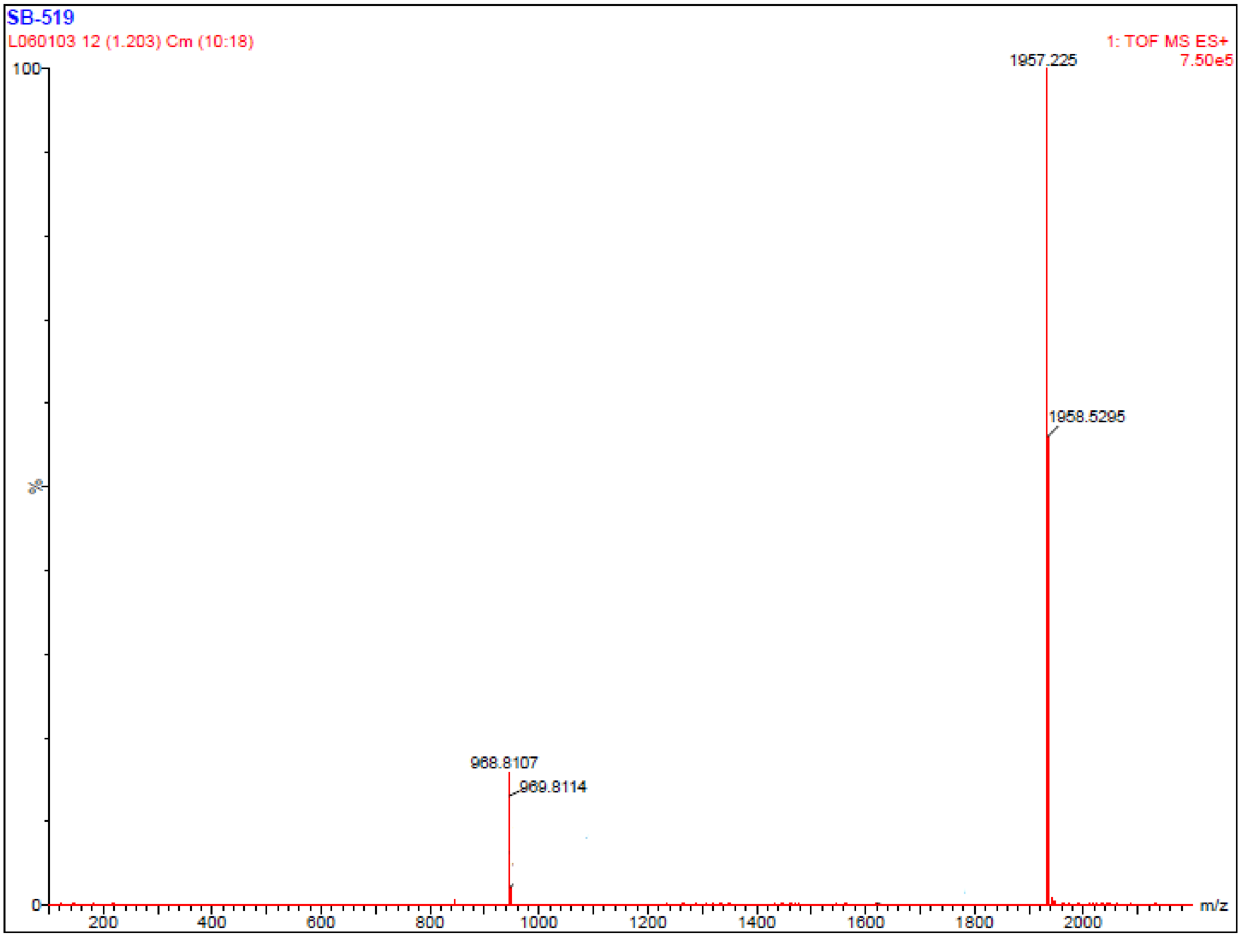


**Figure S20**: Section of HRMS (ESI): *m*/*z* [M + Na]$^+$ spectra for the synthesized peptide **39**. calcd for $C_{102}H_{163}N_{17}O_{19}Na$: 1957.2245; found: 1957.2225.

**Sample Summary:**

| ID | File | Vial | Found | Time | Area Abs | Peak | Calc Mass | Result Mass | Error mDa |
|---|---|---|---|---|---|---|---|---|---|
| njay7-171019-01 | B171019WT002 | 8:7 | YES | 10.33 | 13610 | 1 | 2175.4403<br>1088.2161 | 2175.4401<br>1088.2163 | 1.10<br>1.10 |

**ID Sanjay7-171019-01 File SB171019WT002 Date 23-Oct-2017 Time 14:17:38 Description MDF031177**

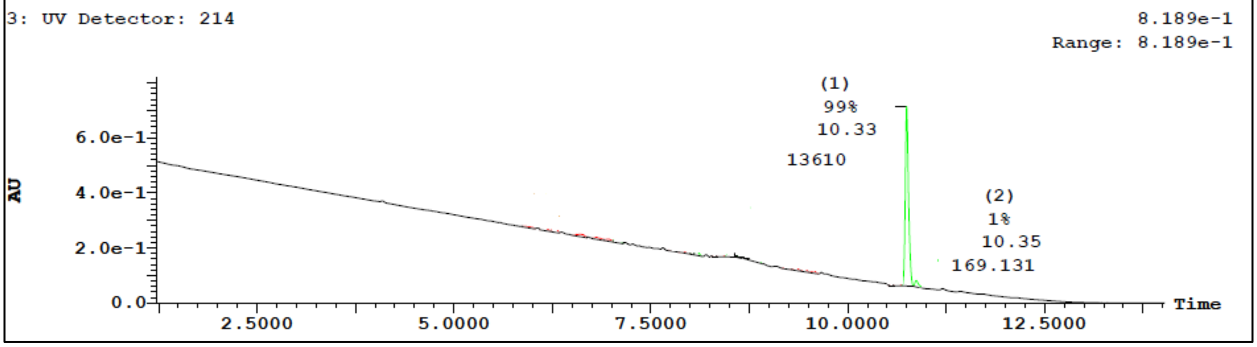


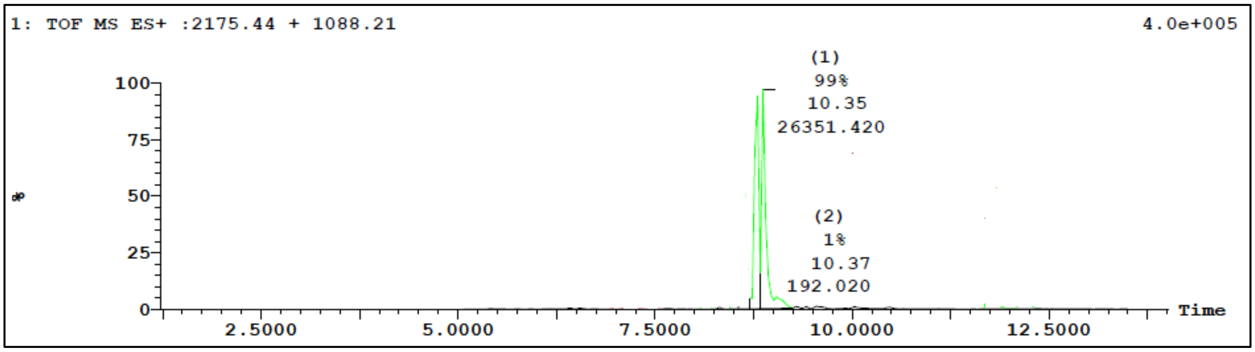


| Peak ID | Time | Compound |
|---|---|---|
| 1 | 10.33 | |

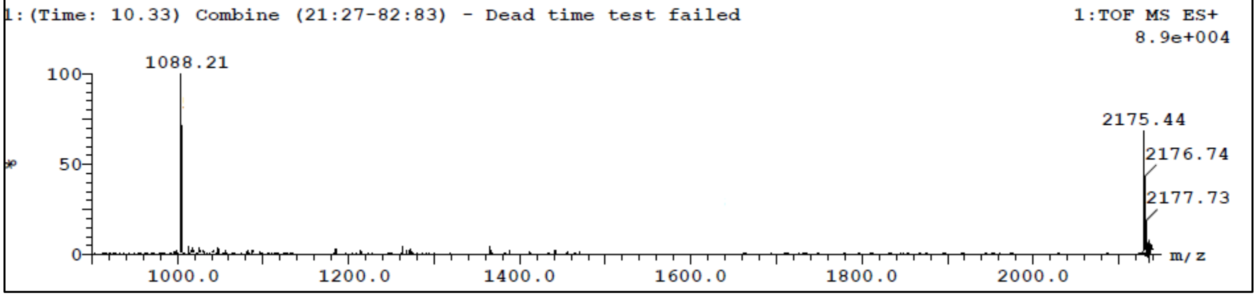


**Figure S21.** LC/TOF-ES-MS spectra of the synthesized peptide **20** is shown in positive mode and the peak at 10.33 min belongs to the peptide **20** (m/z = 2175). The purity of peptide **20** was also determined by HPLC-UV (214 nm)-ESI-MS and was found to be 99%.

Sample Summary:

| ID | File | Vial | Found | Time | Area Abs | Peak | Calc Mass | Result Mass | Error mDa |
|---|---|---|---|---|---|---|---|---|---|
| anjay-171030-01 | B171030WT012 | 8:7 | YES | 7.86 | 48091 | 1 | 1791.1858<br>896.0967 | 1791.1542<br>896.0968 | 2.20 |

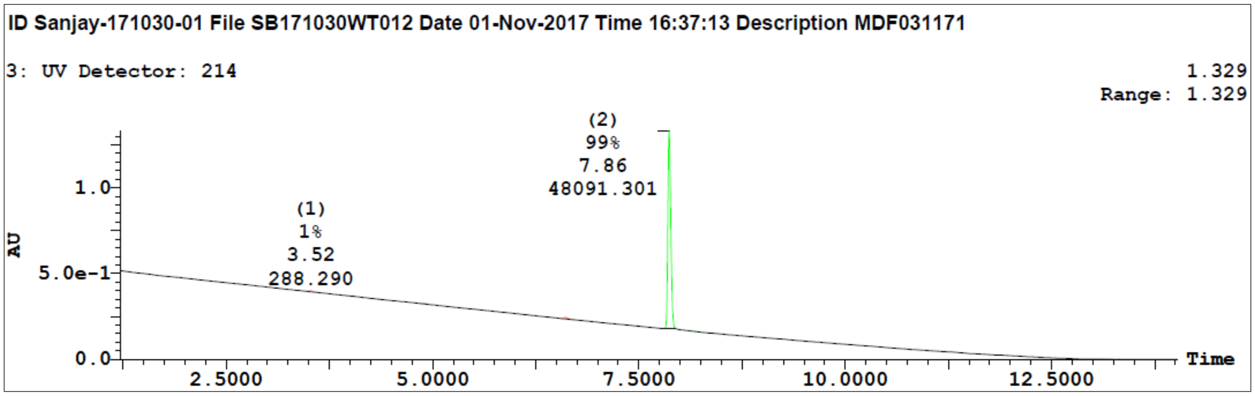


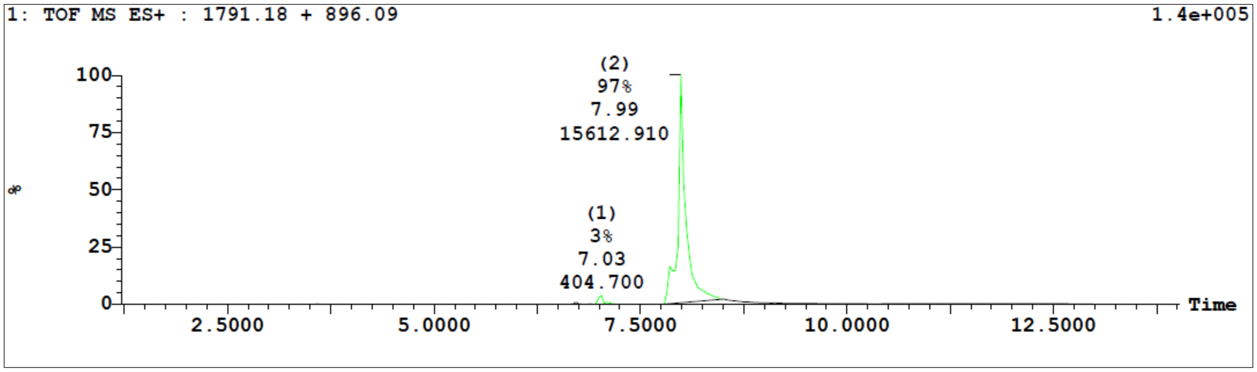


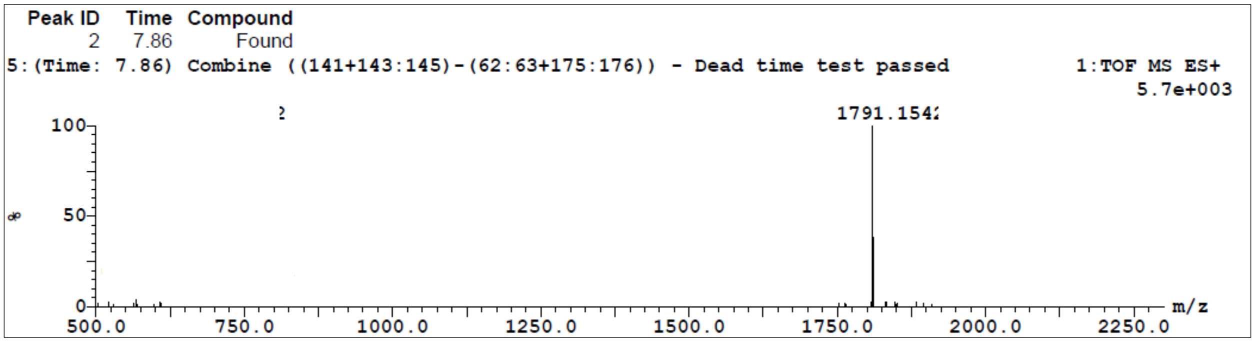


**Figure S22.** LC/TOF-ES-MS spectra of the synthesized peptide **21** is shown in positive mode and the peak at 7.86 min belongs to the peptide **21** (m/z = 1791). The purity of peptide **21** was also determined by HPLC-UV (214 nm)-ESI-MS and was found to be 96%.

**Sample Summary:**

| ID | File | Vial | Found | Time | Area Abs | Peak | Calc Mass | Result Mass | Error mDa |
|---|---|---|---|---|---|---|---|---|---|
| anjay-171030-03 | B171030WT014 | 8:9 | YES | 10.98 | 31537 | 3 | 2005.2871<br>1003.1475 | 2005.2868<br>1003.1474 | -0.90 |

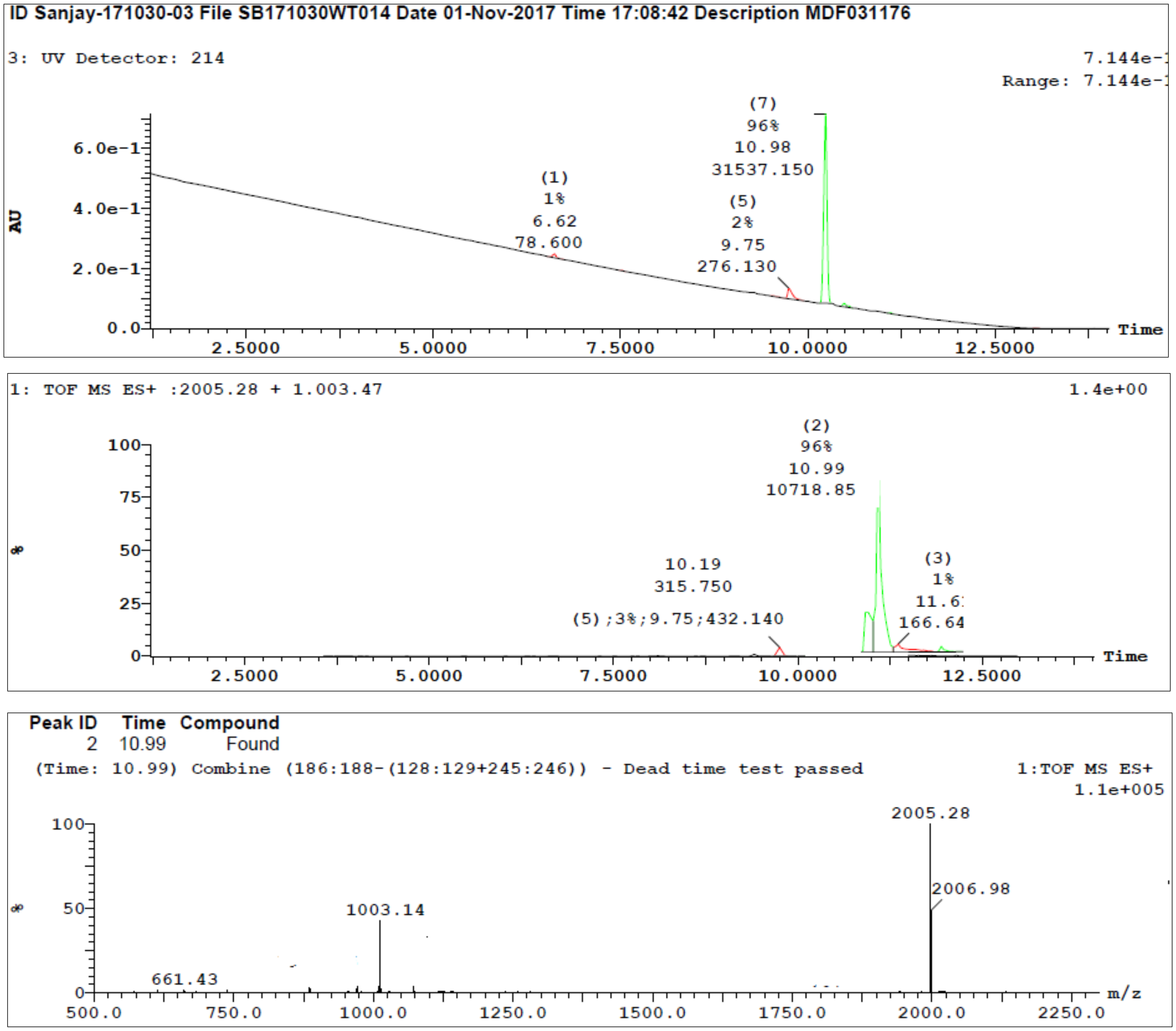


**Figure S23.** LC/TOF-ES-MS spectra of the synthesized peptide **22** is shown in positive mode and the peak at 10.33 min belongs to the peptide **22** (m/z = 2005). The purity of peptide **22** was also determined by HPLC-UV (214 nm)-ESI-MS and was found to be 96%.

**Sample Summary:**

| ID | File | Vial | Found | Time | Area Abs | Peak | Calc Mass | Result Mass | Error mDa |
|---|---|---|---|---|---|---|---|---|---|
| anjay-171107-01 | B171107WT002 | 8:1 | YES | 8.40 | 44190 | 3 | 1963.2402 | 1963.2391 | -3.20 |
| | | | | | | | 982.1241 | 982.0285 | 2.30 |
| | | | | | | | 565.0199 | 0.0000 | |

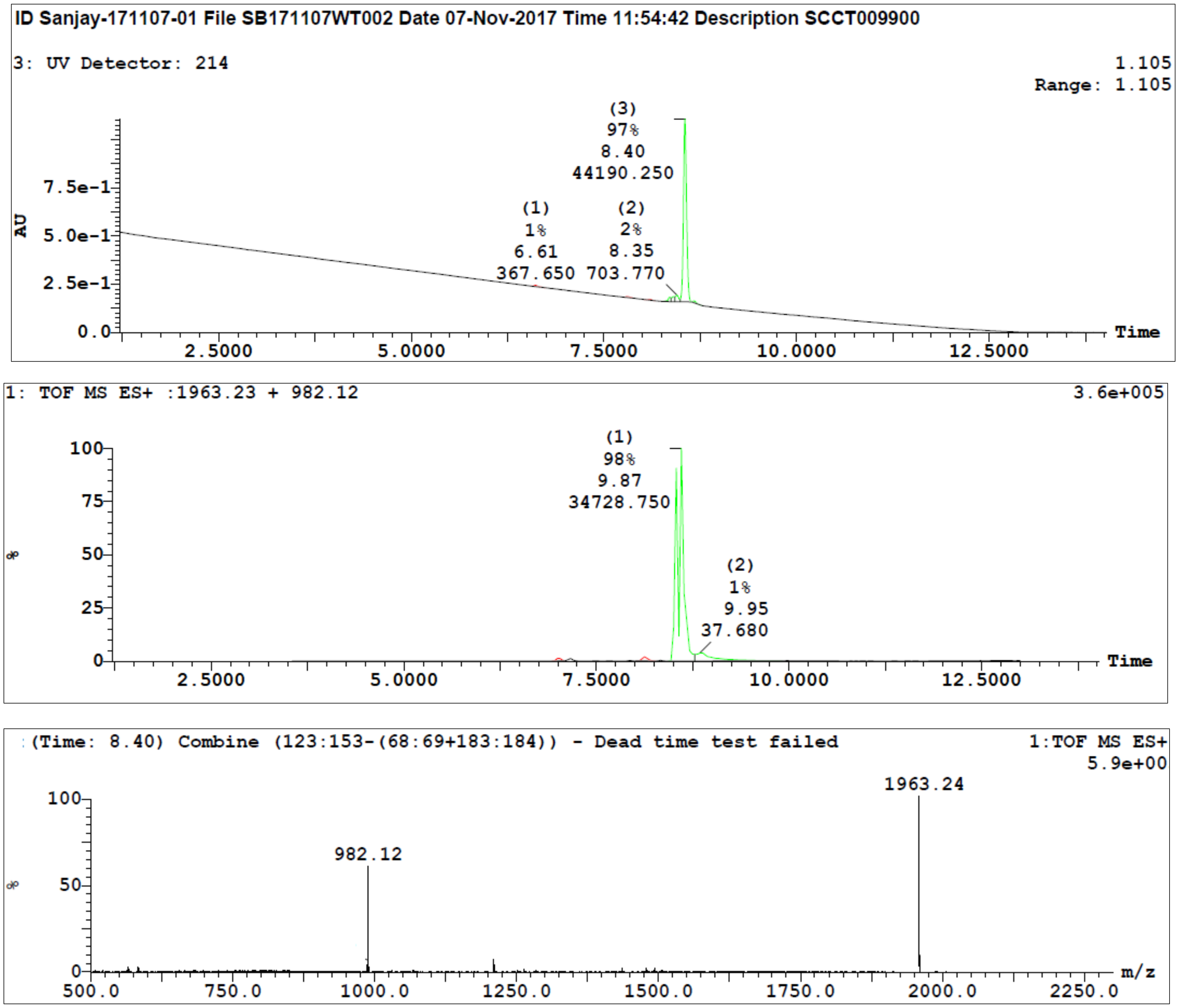


**Figure S24.** LC/TOF-ES-MS spectra of the synthesized peptide **23** is shown in positive mode and the peak at 10.33 min belongs to the peptide **23** (m/z = 1963). The purity of peptide **23** was also determined by HPLC-UV (214 nm)-ESI-MS and was found to be 97%.

**Sample Summary:**

| ID | File | Vial | Found | Time | Area Abs | Peak | Calc Mass | Result Mass | Error mDa |
|---|---|---|---|---|---|---|---|---|---|
| anjay-180207-01 | B180207WT001 | 8:1 | YES | 6.28 | 20418 | 3 | 1907.1776<br>954.0928 | 1907.1725<br>954.0926 | -2.20 |

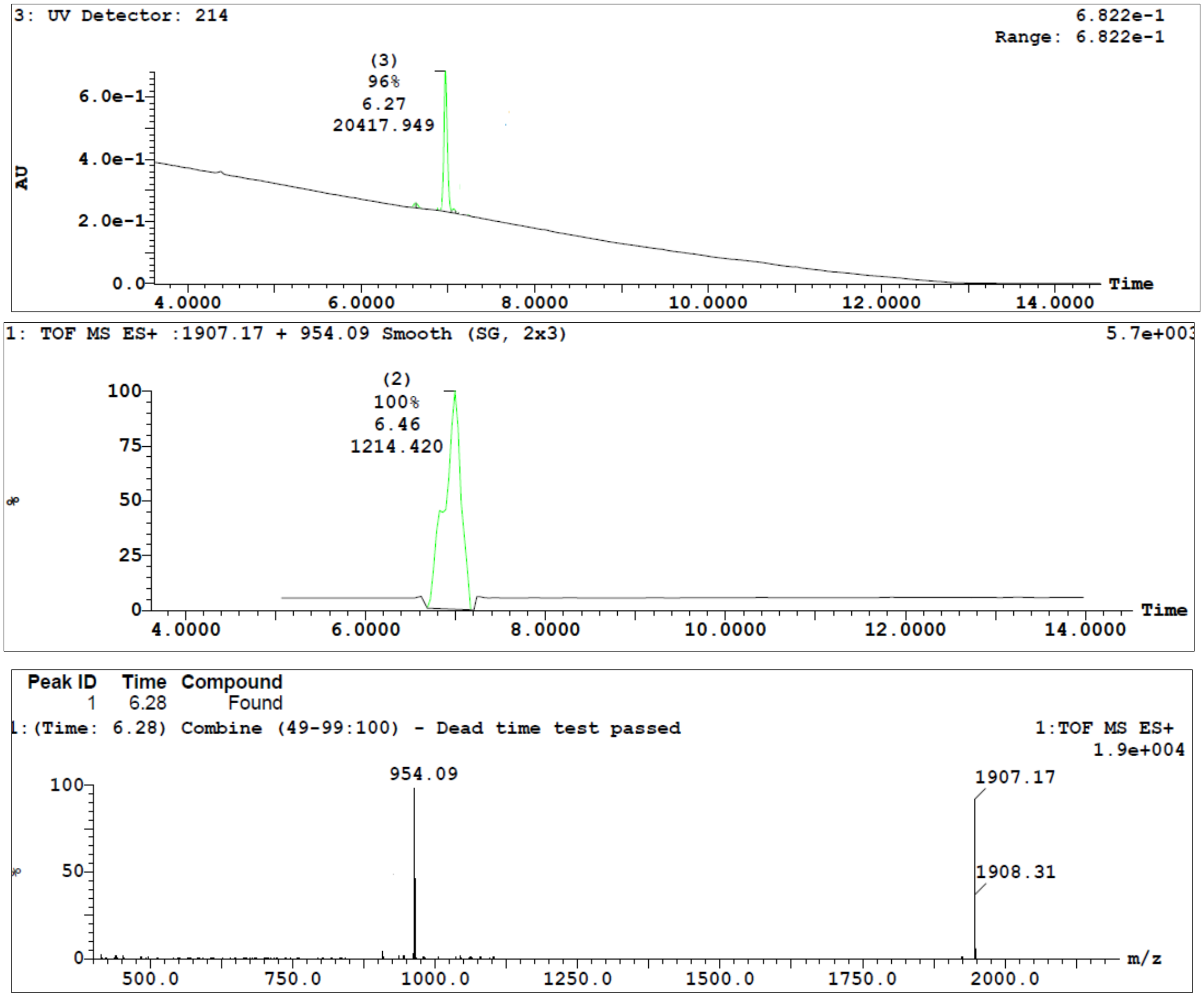


**Figure S25.** LC/TOF-ES-MS spectra of the synthesized peptide **24** is shown in positive mode and the peak at 6.28 min belongs to the peptide **24** (m/z = 1907). The purity of peptide **24** was also determined by HPLC-UV (214 nm)-ESI-MS and was found to be 96%.

Sample Summary:

| ID | File | Vial | Found | Time | Area Abs | Peak | Calc Mass | Result Mass | Error mDa |
|---|---|---|---|---|---|---|---|---|---|
| anjay-180116-04 | B180117WT005 | 8:4 | YES | 6.48 | 43744 | 3 | 2019.3028 | 2019.3027 | 0.00 |

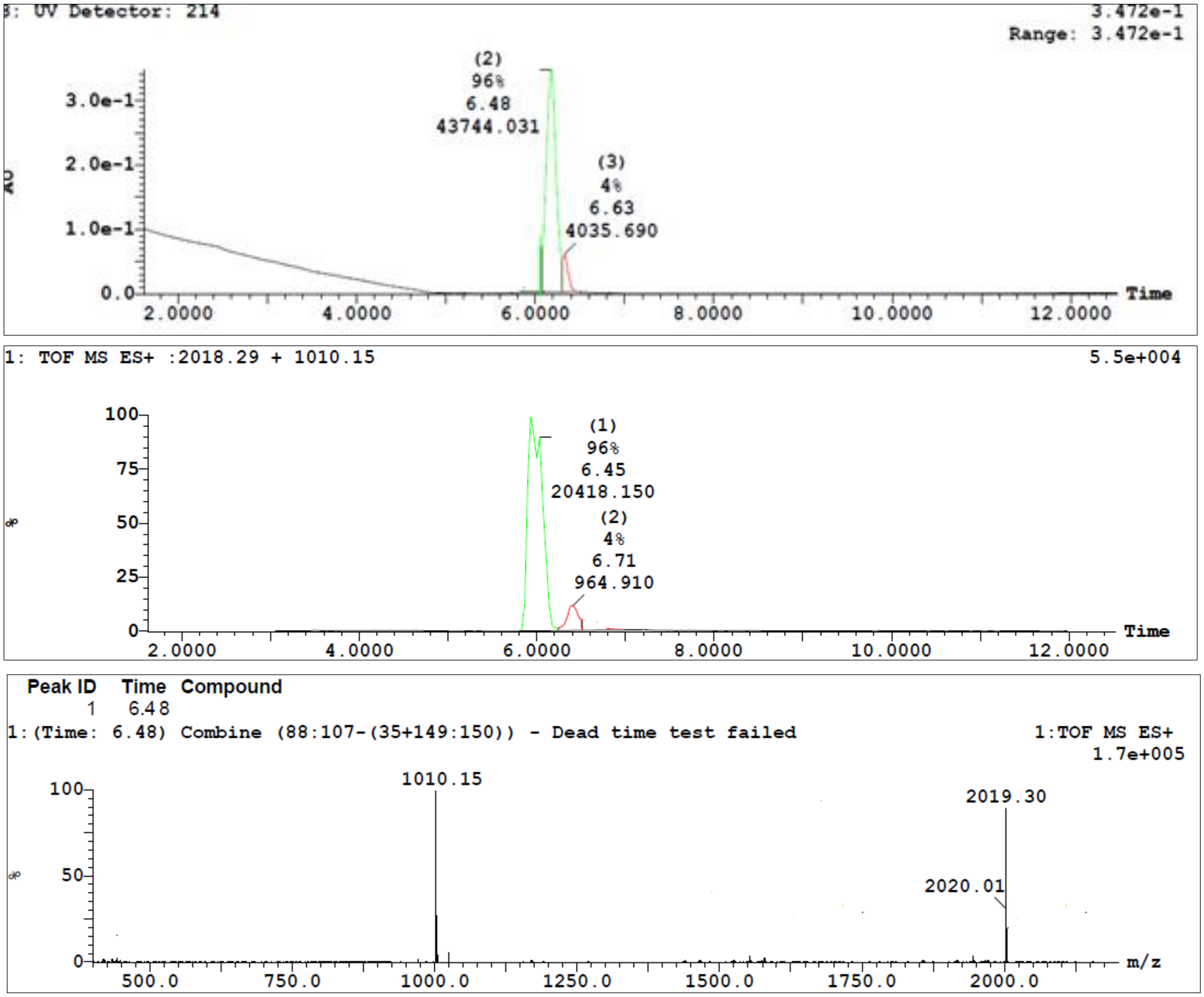


**Figure S26.** LC/TOF-ES-MS spectra of the synthesized peptide **25** is shown in positive mode and the peak at 6.48 min belongs to the peptide **25** (m/z = 2019). The purity of peptide **25** was also determined by HPLC-UV (214 nm)-ESI-MS and was found to be 96%.

**Sample Summary:**

| ID | File | Vial | Found | Time | Area Abs | Peak | Calc Mass | Result Mass | Error mDa |
|---|---|---|---|---|---|---|---|---|---|
| anjay-180117-01 | B180118WT002 | 8:1 | YES | 6.88 | 1132 | 1 | 1963.2402<br>982.1241 | 1963.2386<br>982.0012 | 2.70<br>-3.80 |
| | | | YES | 6.99 | 56460 | 2 | 1963.2402<br>982.1241 | 1963.2386<br>982.0028 | 4.10<br>-2.20 |

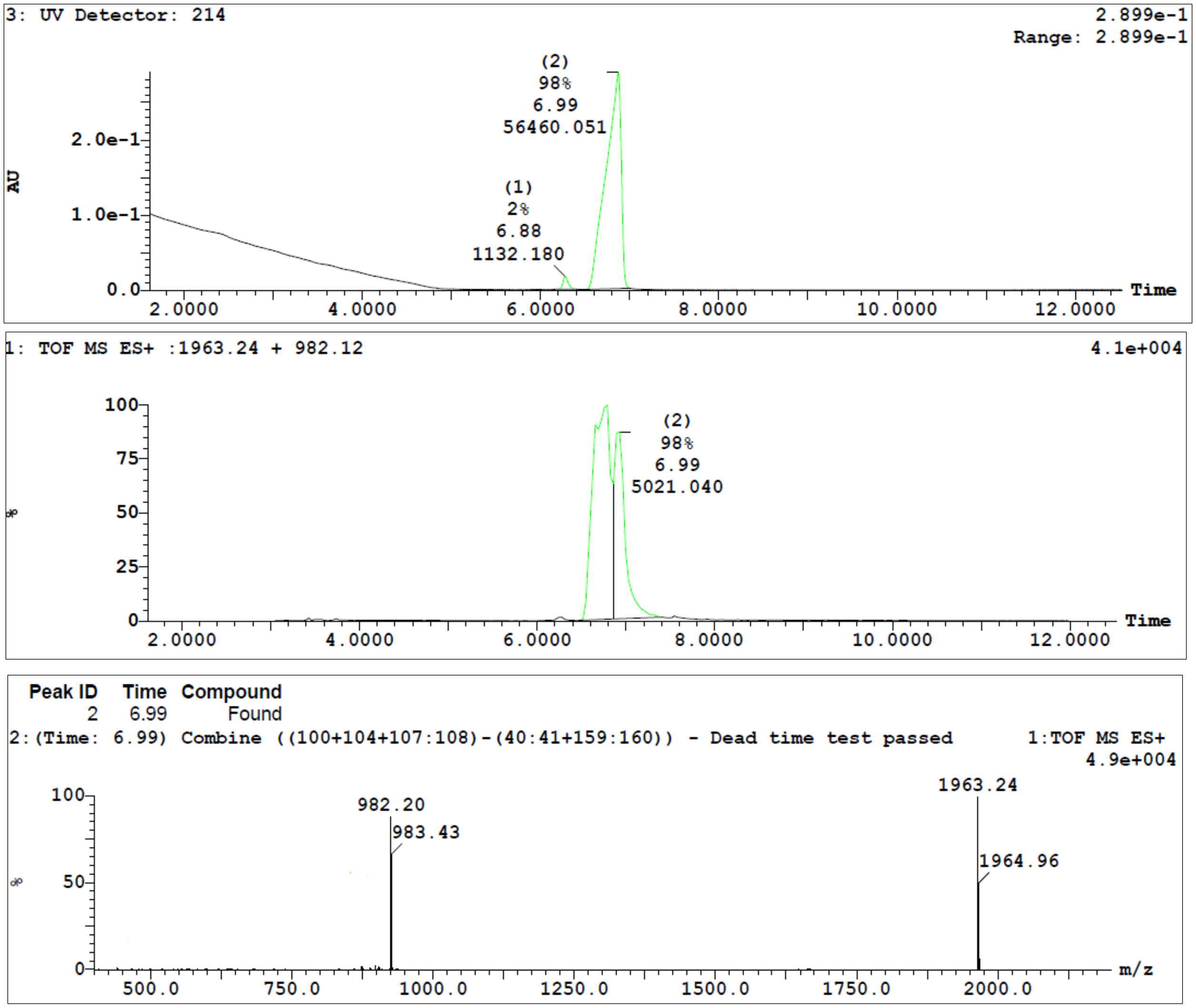


**Figure S27.** LC/TOF-ES-MS spectra of the synthesized peptide **26** is shown in positive mode and the peak at 6.99 min belongs to the peptide **26** (m/z = 1963). The purity of peptide **26** was also determined by HPLC-UV (214 nm)-ESI-MS and was found to be 98%.

**Sample Summary:**

| ID | File | Vial | Found | Time | Area Abs | Peak | Calc Mass | Result Mass | Error mDa |
|---|---|---|---|---|---|---|---|---|---|
| -Anal-170901-03 | B170901WT005 | 8:15 | YES | 4.78 | 6496 | 2 | 1921.1882 | 1921.1803 | -8.20 |
| | | | | | | | 961.0981 | 961.0488 | -2.70 |
| | | | | | | | 531.9971 | 0.0000 | |

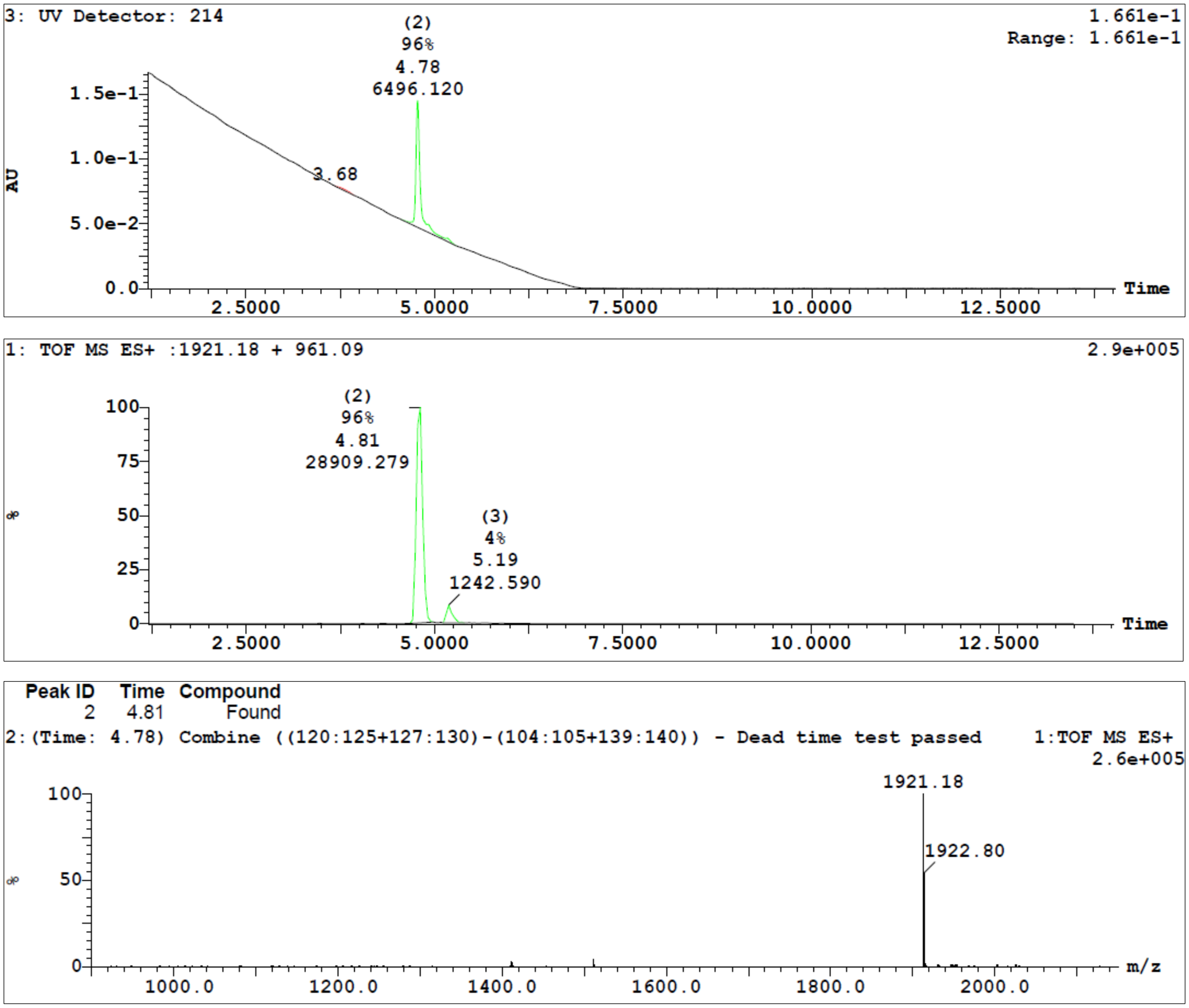


**Figure S28.** LC/TOF-ES-MS spectra of the synthesized peptide **27** is shown in positive mode and the peak at 4.78 min belongs to the peptide **27** (m/z = 1921). The purity of peptide **27** was also determined by HPLC-UV (214 nm)-ESI-MS and was found to be 96%.

**Sample Summary:**

| ID | File | Vial | Found | Time | Area Abs | Peak | Calc Mass | Result Mass | Error mDa |
|---|---|---|---|---|---|---|---|---|---|
| anjay-180117-04 | B180118WT005 | 8:4 | YES | 8.03 | 10108 | 2 | 1865.1306 | 1865.1284 | 0.70 |
| | | | | | | | 933.0693 | 933.0332 | -3.10 |

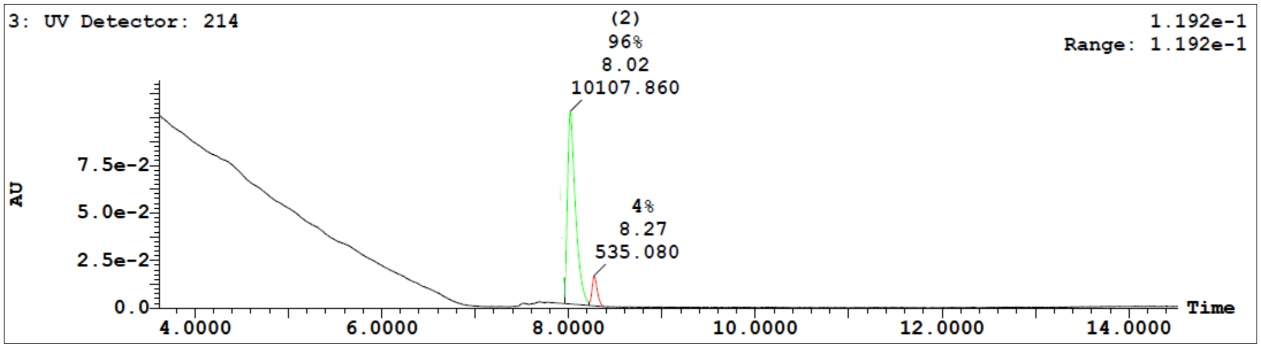


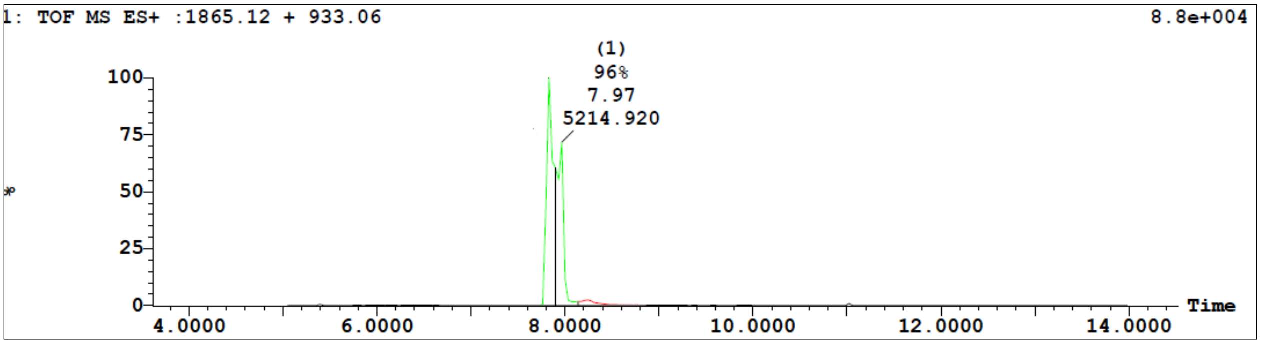


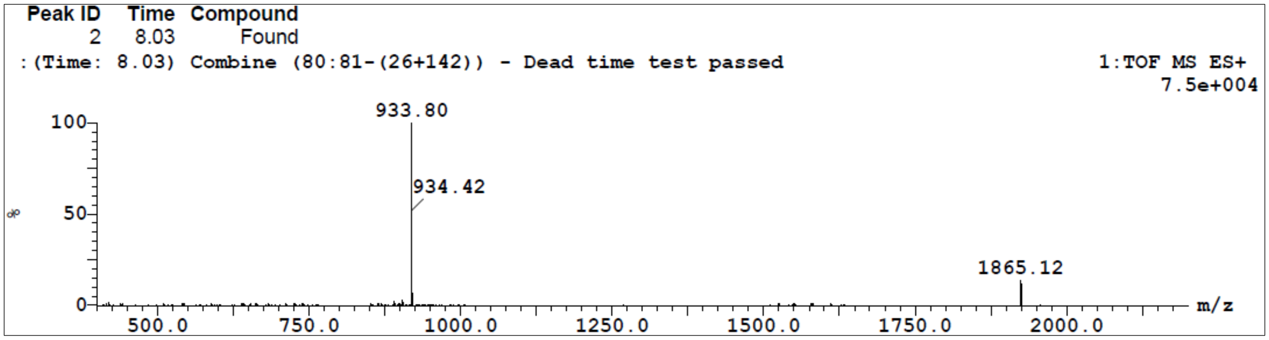


**Figure S29.** LC/TOF-ES-MS spectra of the synthesized peptide **28** is shown in positive mode and the peak at 8.03 min belongs to the peptide **28** (m/z = 1865). The purity of peptide **28** was also determined by HPLC-UV (214 nm)-ESI-MS and was found to be 96%.

**Sample Summary:**

| ID | File | Vial | Found | Time | Area Abs | Peak | Calc Mass | Result Mass | Error mDa |
|---|---|---|---|---|---|---|---|---|---|
| anjay-180117-02 | B180118WT003 | 8:2 | YES | 8.55 | 44559 | 2 | 1921.1932 | 1921.1939 | 1.20 |
| | | | | | | | 961.0050 | 961.0062 | 1.00 |

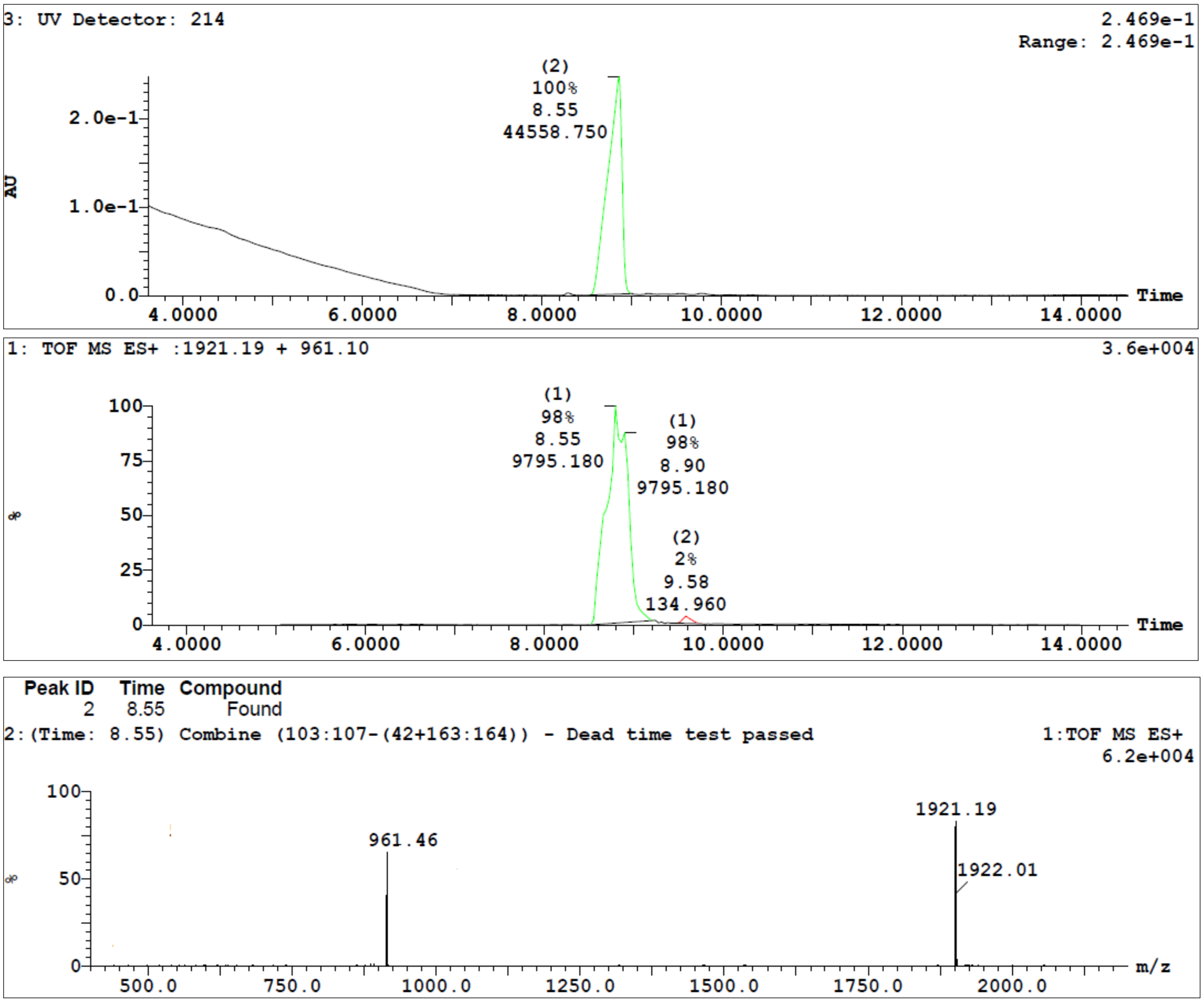


**Figure S30.** LC/TOF-ES-MS spectra of the synthesized peptide **29** is shown in positive mode and the peak at 8.55 min belongs to the peptide **29** (m/z = 1921). The purity of peptide **29** was also determined by HPLC-UV (214 nm)-ESI-MS and was found to be 98%.

**Sample Summary:**

| ID | File | Vial | Found | Time | Area Abs | Peak | Calc Mass | Result Mass | Error mDa |
|---|---|---|---|---|---|---|---|---|---|
| anjay-171113-01 | B171113WT002 | 8:13 | YES | 7.31 | 63218 | 3 | 2191.3737<br>1095.1828 | 2191.3842<br>1095.1828 | 1.90 |

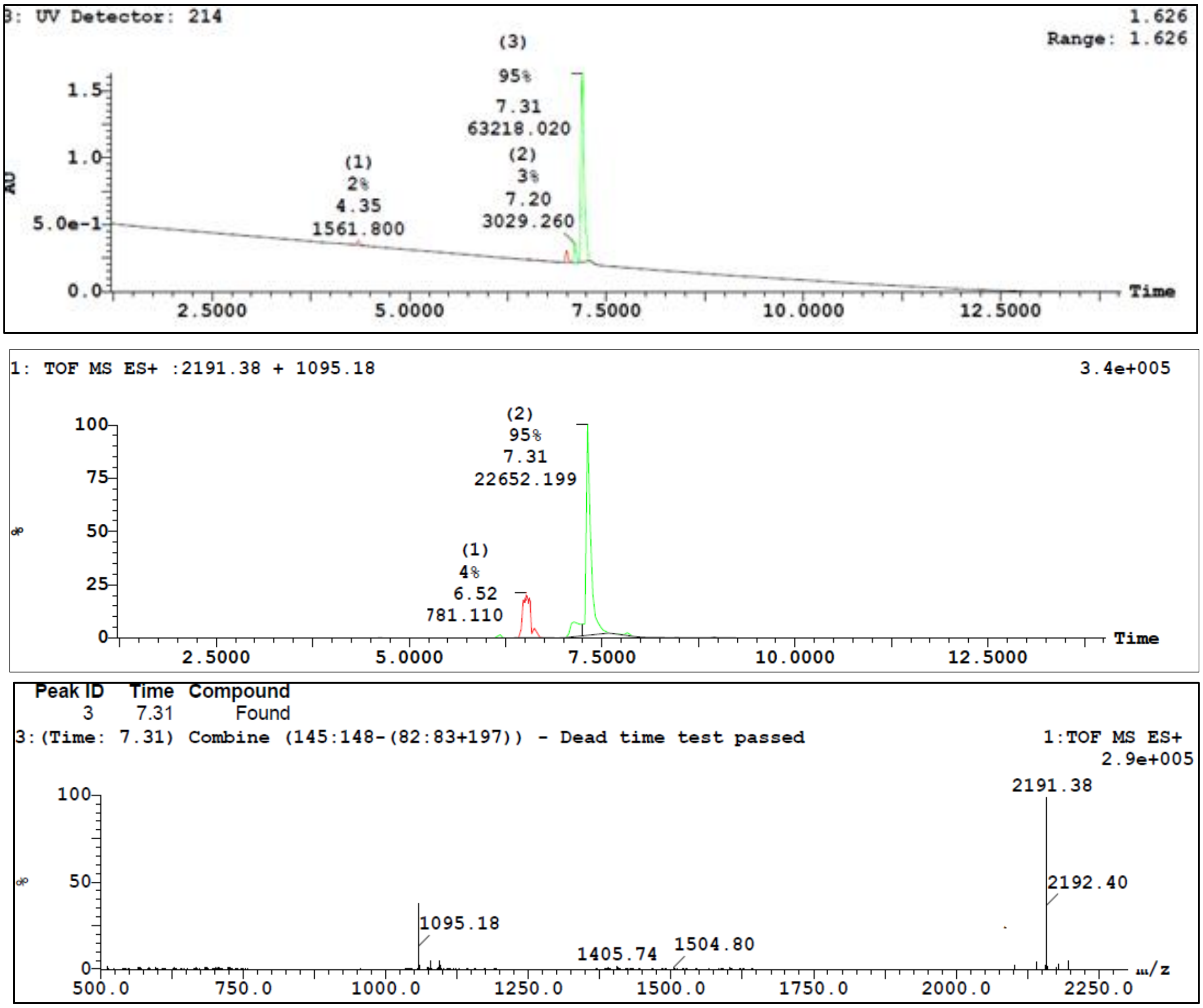


**Figure S31.** LC/TOF-ES-MS spectra of the synthesized peptide **30** is shown in positive mode and the peak at 7.31 min belongs to the peptide **30** (m/z = 2191). The purity of peptide **30** was also determined by HPLC-UV (214 nm)-ESI-MS and was found to be 95%.

**Sample Summary:**

| ID | File | Vial | Found | Time | Area Abs | Peak | Calc Mass | Result Mass | Error mDa |
|---|---|---|---|---|---|---|---|---|---|
| -Anal-180413-03 | B180413WT009 | 8:15 | YES | 10.49 | 26084 | 2 | 1738.0925<br>869.5502 | 1738.1045<br>869.5504 | 2.00 |

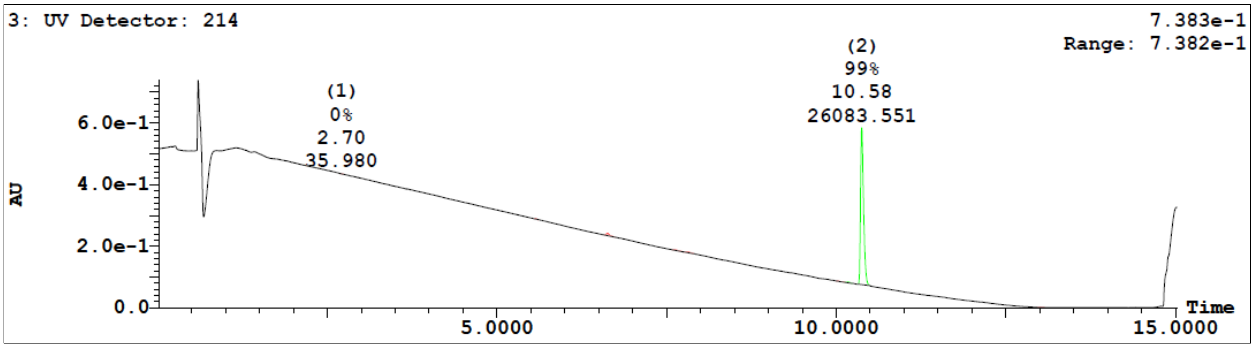


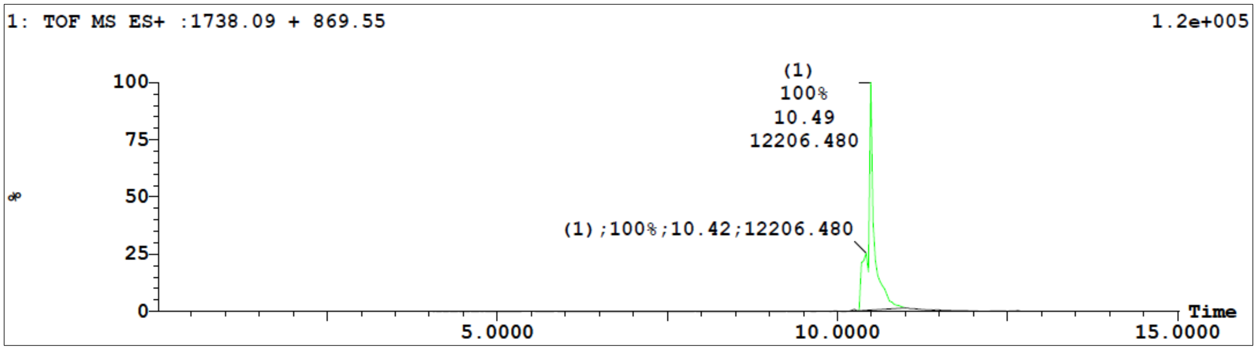


| Peak ID | Time | Compound |
|---|---|---|
| 1 | 10.49 | Found |

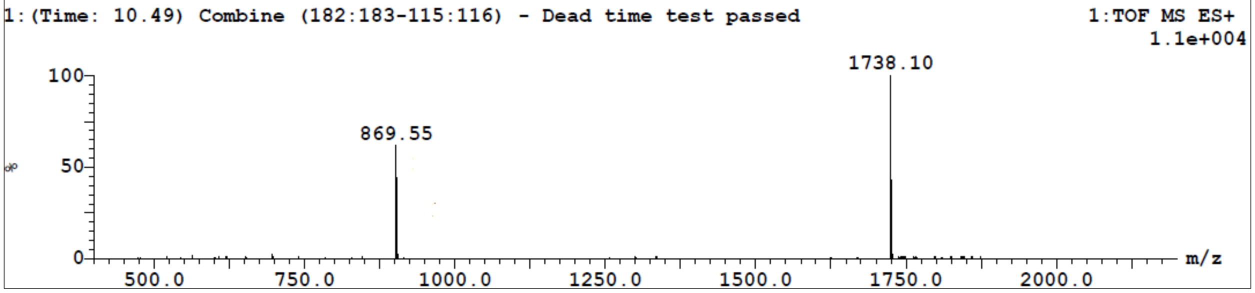


**Figure S33.** LC/TOF-ES-MS spectra of the synthesized peptide **33** is shown in positive mode and the peak at 10.49 min belongs to the peptide **33** (m/z = 1738). The purity of peptide **33** was also determined by HPLC-UV (214 nm)-ESI-MS and was found to be 99%.

**Sample Summary:**

| ID | File | Vial | Found | Time | Area Abs | Peak | Calc Mass | Result Mass | Error mDa |
|---|---|---|---|---|---|---|---|---|---|
| anjay-180213-02 | B180213WT005 | 8:8 | YES | 9.52 | 18069 | 2 | 1809.1296<br>905.0688 | 1809.1274<br>0.0000 | -4.50 |

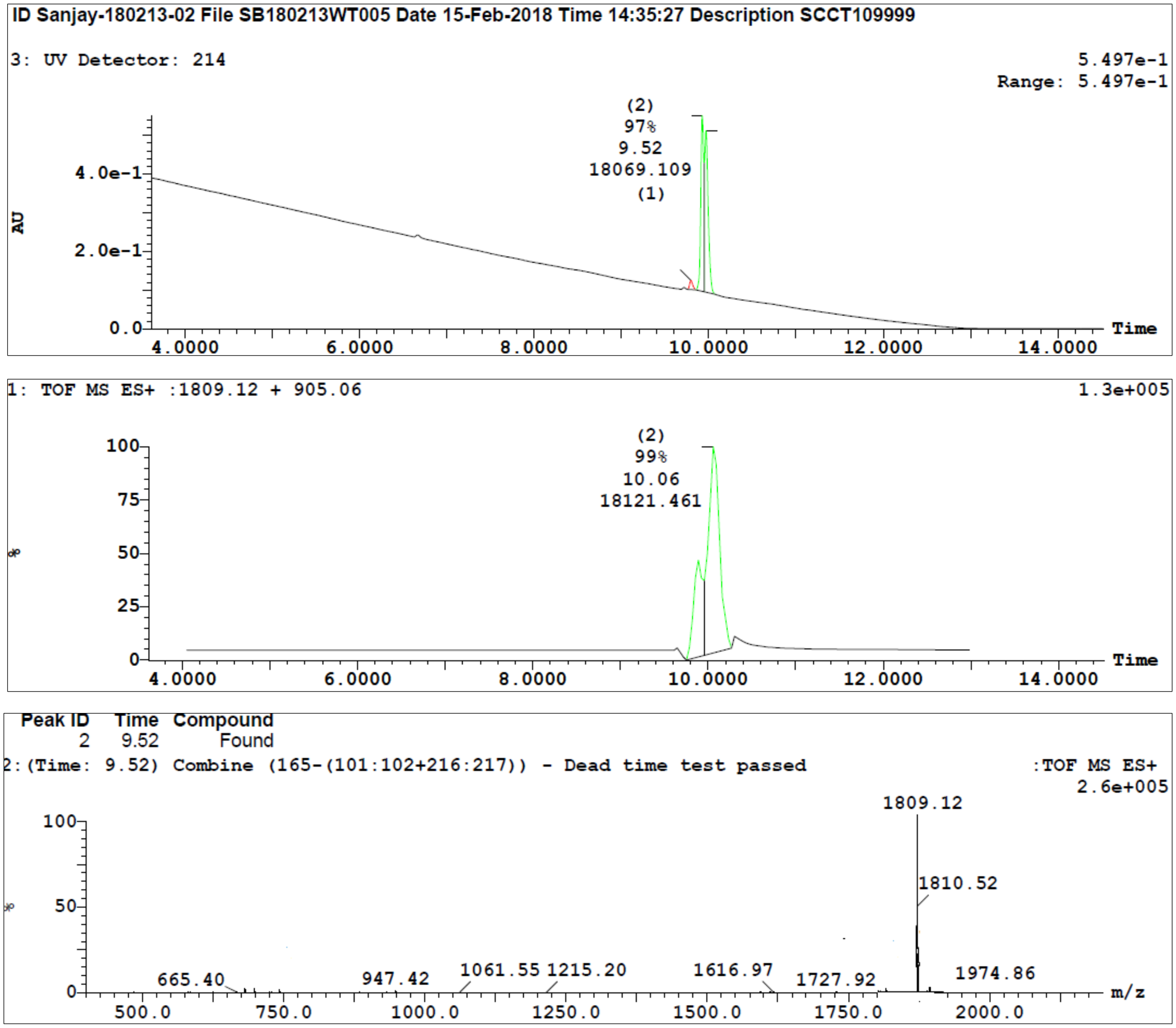


**Figure S34.** LC/TOF-ES-MS spectra of the synthesized peptide **34** is shown in positive mode and the peak at 9.52 min belongs to the peptide **34** (m/z = 1809). The purity of peptide **34** was also determined by HPLC-UV (214 nm)-ESI-MS and was found to be 97%.

**Sample Summary:**

| ID | File | Vial | Found | Time | Area Abs | Peak | Calc Mass | Result Mass | Error mDa |
|---|---|---|---|---|---|---|---|---|---|
| anjay-180117-01 | B180118WT002 | 8:1 | YES | 9.30 | 56460 | 3 | 1837.1609<br>919.0844 | 1837.1654<br>919.0028 | 4.10<br>-2.20 |

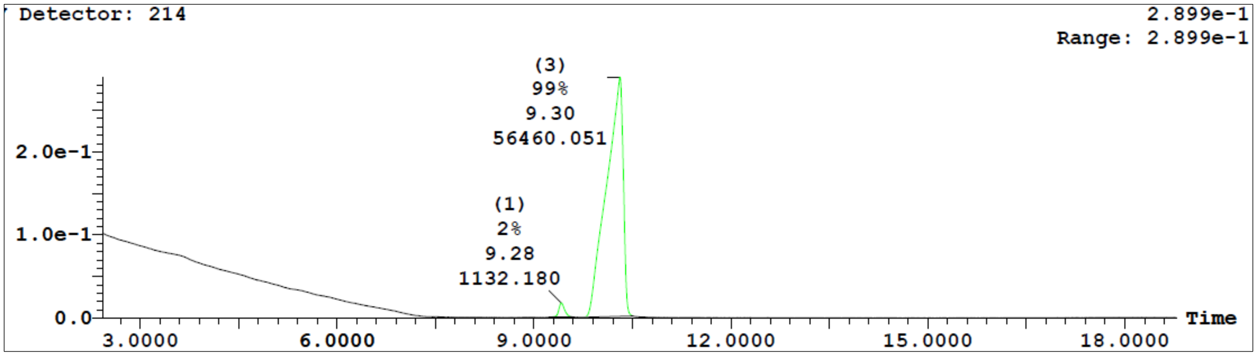


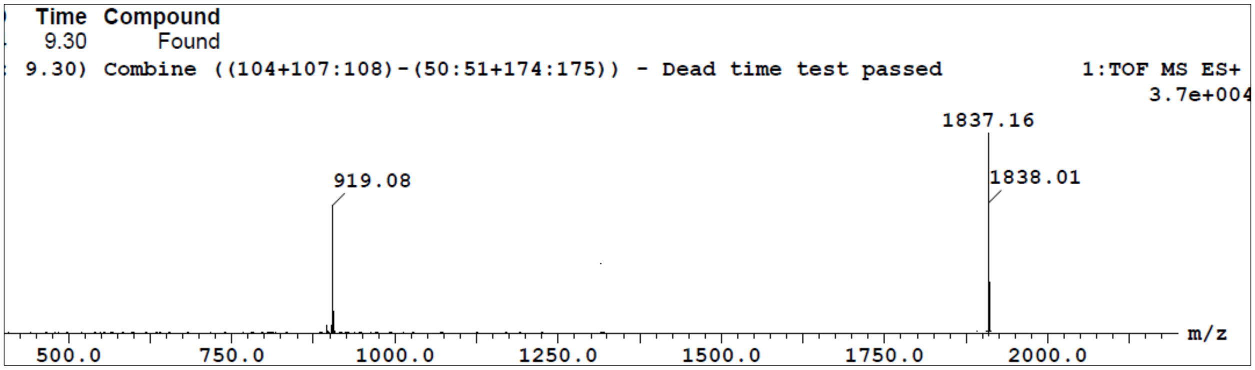


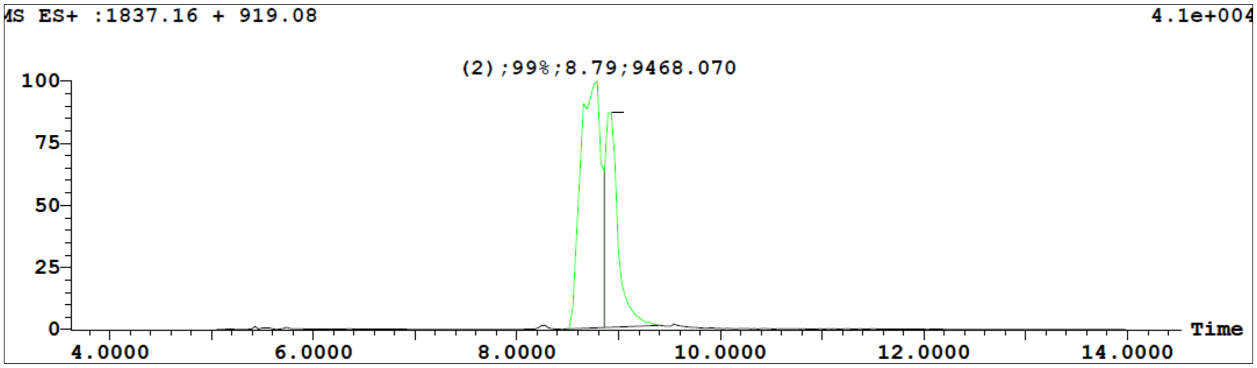


**Figure S35.** LC/TOF-ES-MS spectra of the synthesized peptide **35** is shown in positive mode and the peak at 9.30 min belongs to the peptide **35** (m/z = 1837). The purity of peptide **35** was also determined by HPLC-UV (214 nm)-ESI-MS and was found to be 99%.

**Sample Summary:**

| ID | File | Vial | Found | Time | Area Abs | Peak | Calc Mass | Result Mass | Error mDa |
|---|---|---|---|---|---|---|---|---|---|
| anjay-180116-04 | B180117WT005 | 8:4 | YES | 8.40 | 43744 | 3 | 1850.1813<br>925.5946 | 1850.1381<br>925.6842 | 5.20<br>4.20 |

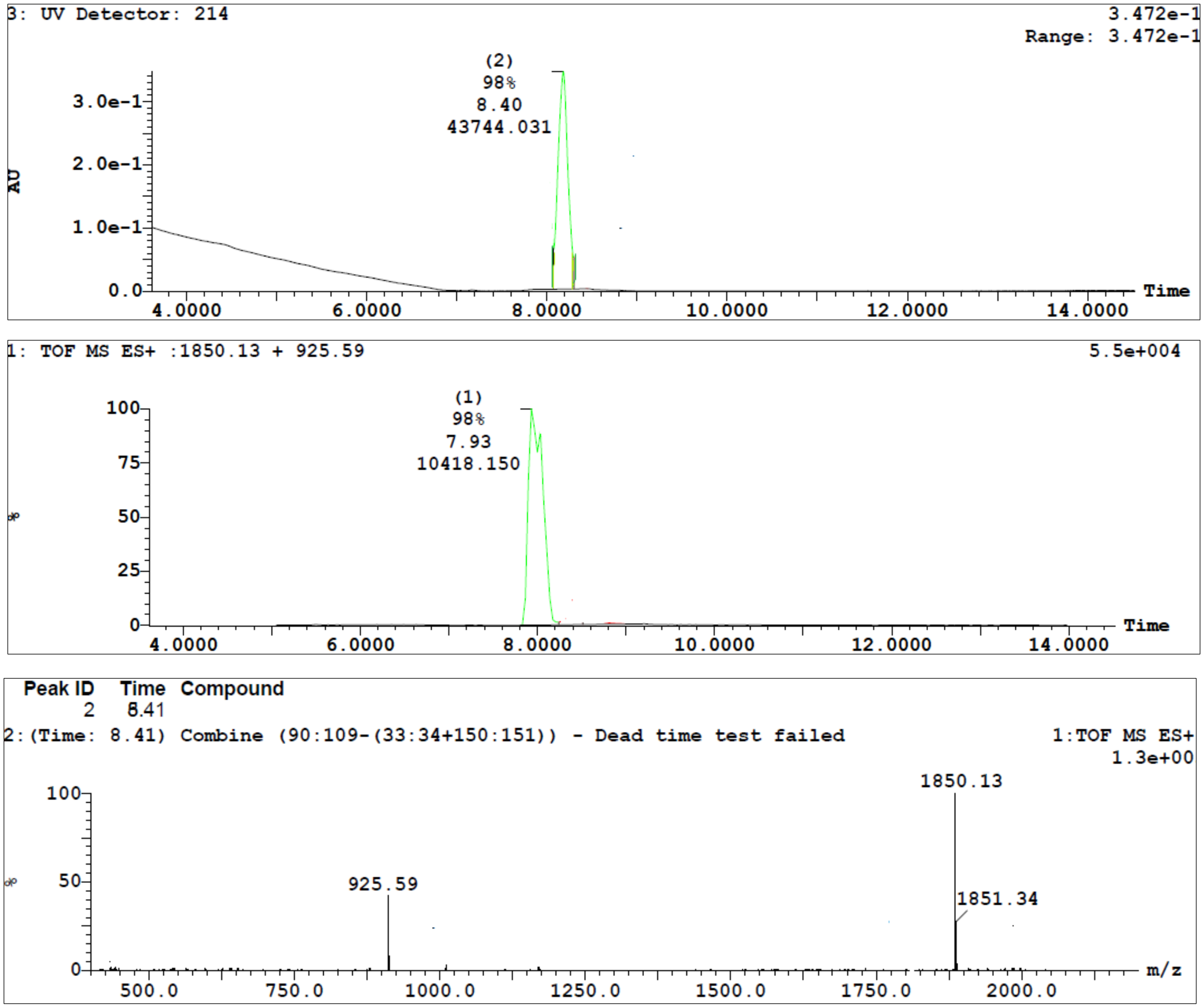


**Figure S36.** LC/TOF-ES-MS spectra of the synthesized peptide **36** is shown in positive mode and the peak at 8.41 min belongs to the peptide **36** (m/z = 1850). The purity of peptide **36** was also determined by HPLC-UV (214 nm)-ESI-MS and was found to be 98%.

**Sample Summary:**

| ID | File | Vial | Found | Time | Area Abs | Peak | Calc Mass | Result Mass | Error mDa |
|---|---|---|---|---|---|---|---|---|---|
| Sanjay-190326-4 | 190326WT004b | 8:10 | YES | 8.24 | 21466 | 2 | 1893.2235<br>947.1157 | 1893.2325<br>947.0673 | 1.40 |

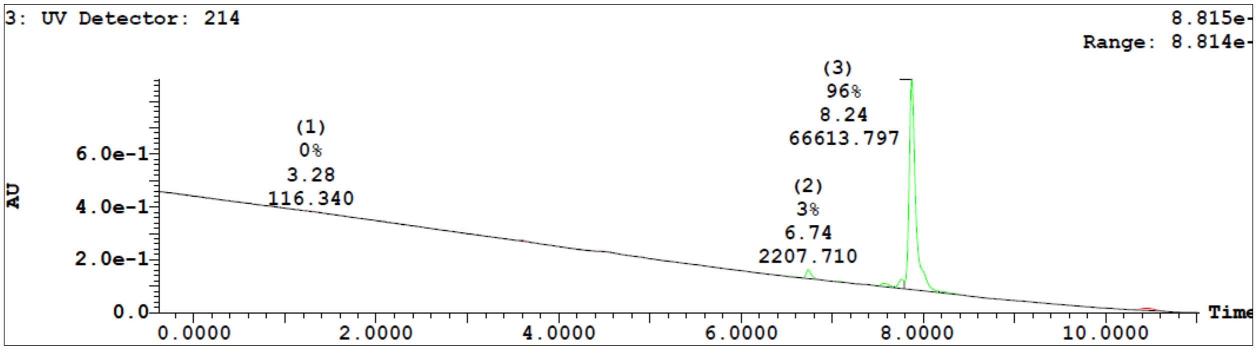


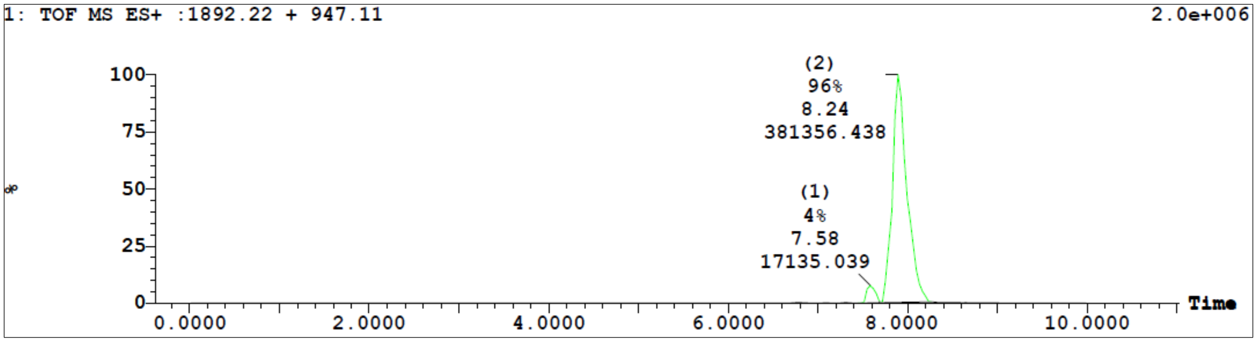


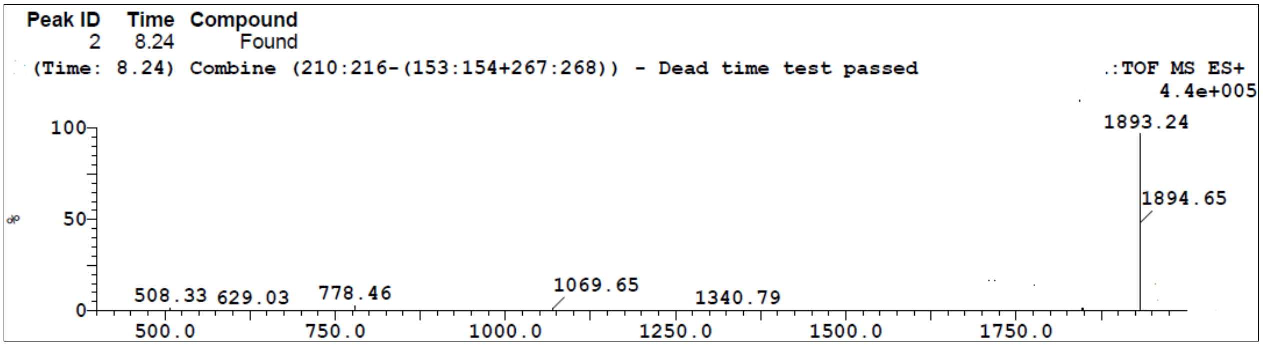


**Figure S37.** LC/TOF-ES-MS spectra of the synthesized peptide **37** is shown in positive mode and the peak at 8.24 min belongs to the peptide **37** (m/z = 1893). The purity of peptide **37** was also determined by HPLC-UV (214 nm)-ESI-MS and was found to be 96%.

**Sample Summary:**

| ID | File | Vial | Found | Time | Area Abs | Peak | Calc Mass | Result Mass | Error mDa |
|---|---|---|---|---|---|---|---|---|---|
| anjay-181212-01 | B181212WT002 | 8:27 | YES | 8.24 | 24175 | 2 | 1921.2548<br>961.1314 | 1921.2845<br>961.6888 | 2.30<br>11.30 |

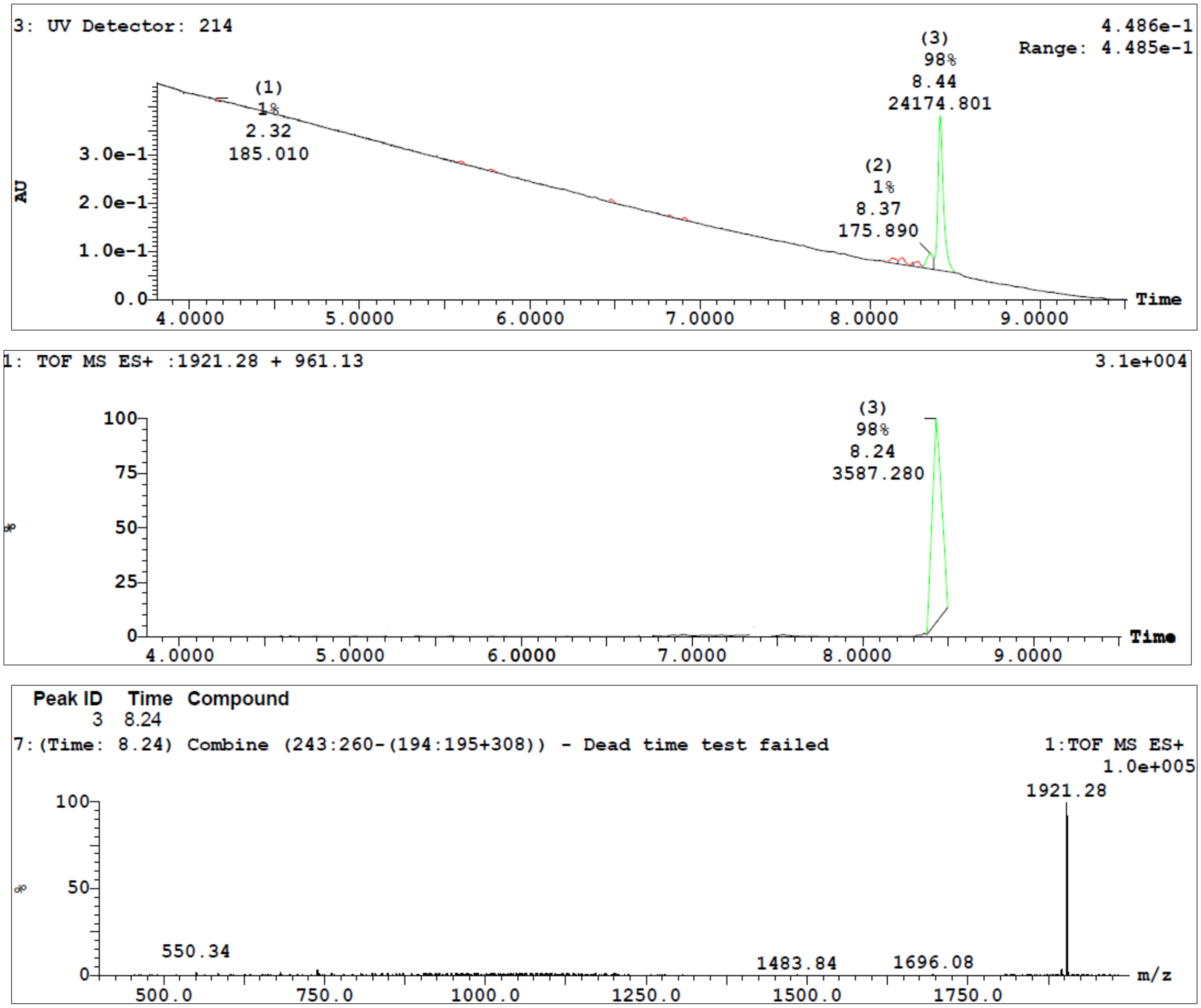


**Figure S38.** LC/TOF-ES-MS spectra of the synthesized peptide **38** is shown in positive mode and the peak at 8.24 min belongs to the peptide **38** (m/z = 1921). The purity of peptide **38** was also determined by HPLC-UV (214 nm)-ESI-MS and was found to be 98%.

**Sample Summary:**

| ID | File | Vial | Found | Time | Area Abs | Peak | Calc Mass | Result Mass | Error mDa |
|---|---|---|---|---|---|---|---|---|---|
| anjay-180618-02 | B180618WT007 | 8:26 | YES | 8.35 | 60891 | 1 | 1935.2704 | 1935.2407 | -7.40 |
| | | | | | | | 968.1352 | 968.1095 | -1.50 |

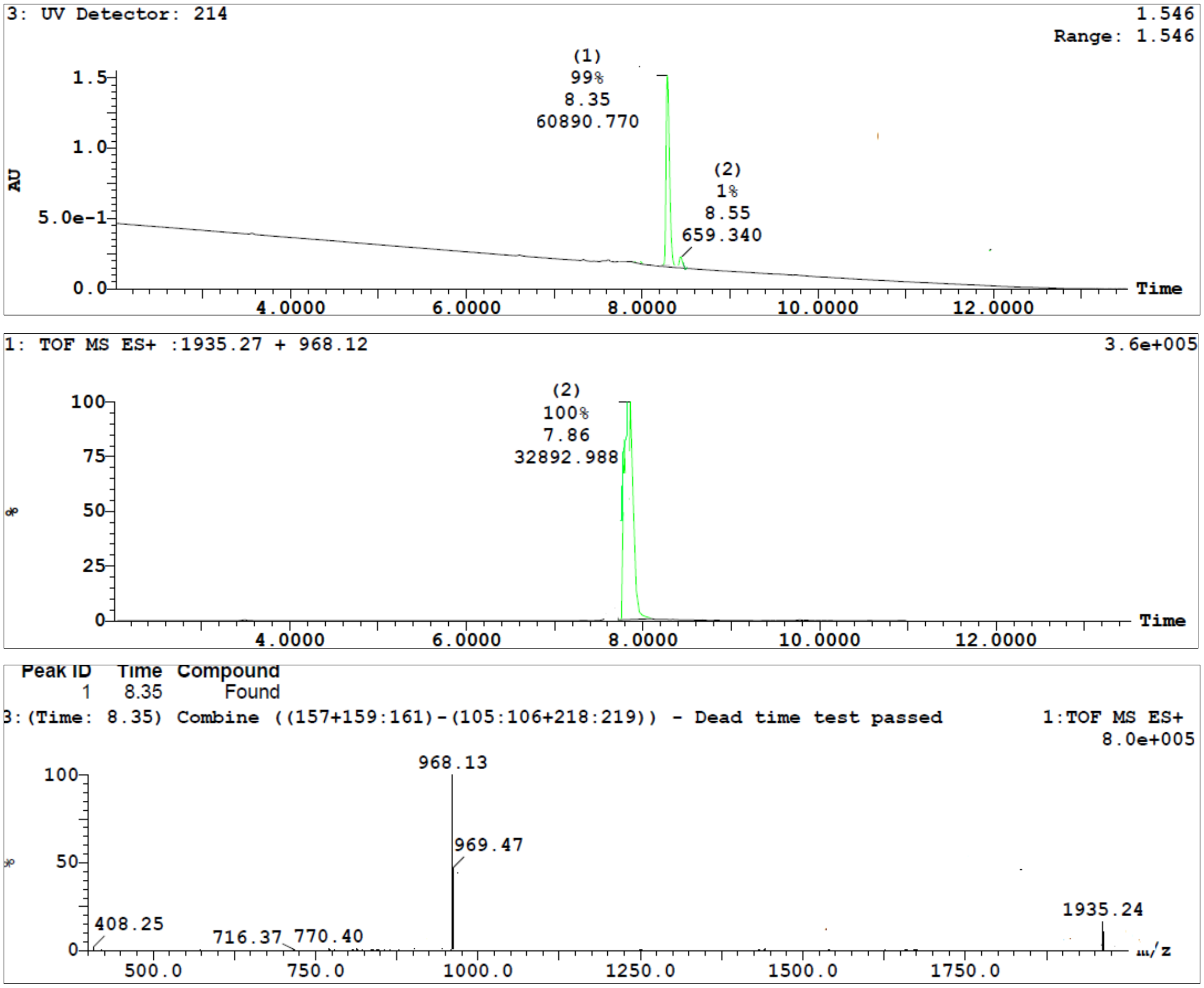


**Figure S39.** LC/TOF-ES-MS spectra of the synthesized peptide **39** is shown in positive mode and the peak at 8.35 min belongs to the peptide **39** (m/z = 1935). The purity of peptide **39** was also determined by HPLC-UV (214 nm)-ESI-MS and was found to be 99%.
